# MSH2 stimulates interfering and inhibits non-interfering crossovers in response to genetic polymorphism

**DOI:** 10.1101/2023.05.03.539183

**Authors:** Julia Dluzewska, Wojciech Dziegielewski, Maja Szymanska-Lejman, Monika Gazecka, Ian R. Henderson, James D. Higgins, Piotr A. Ziolkowski

## Abstract

In Arabidopsis, local sequence polymorphism between homologs can stimulate double-strand break (DSB) repair via crossover formation in a MSH2-dependent manner. To understand how MSH2 regulates crossovers formed by the independent interfering and non-interfering pathways, we combine mutants that elevate non-interfering crossovers with *msh2*. We demonstrate that MSH2 blocks non-interfering crossovers at polymorphic loci, which is the opposite effect to interfering crossovers. We also observe MSH2-independent crossover inhibition at highly polymorphic sites. We measure recombination along the chromosome arms, in lines differing in the heterozygosity pattern, and observe a dramatic crossover increase at the boundaries between heterozygous and homozygous regions, which is MSH2-dependent. Together, we show that MSH2 is a master regulator of meiotic DSB repair, with antagonistic effects on interfering and non-interfering crossovers, that shapes the crossover landscape in relation to interhomolog polymorphism.

## Introduction

Sexual reproduction is based on the fusion of gametes formed by a specific cell division called meiosis^1,2^. During meiosis, chromosomes from both parents pair up and exchange genetic information with each other through crossover recombination^2,3^. Crossovers are initiated with programmed double-strand breaks (DSBs), most of which however are eventually repaired as non-crossovers^1–3^. Crossovers create new allelic combinations, which are crucial for adaptation, diversity and evolution^4–6^. Moreover, they are necessary for proper chromosome segregation in meiosis, hence non-recombining mutants are sterile^2,3,7,8^. In most eukaryotes, two crossover classes exist, that are produced in different pathways^9,10^ and are controlled differently: Class I crossovers arise in the ZMM pathway named after its constituent proteins (ZIP1-4, MSH4/5 and MER3), and inactivation of any protein in this pathway leads to its complete shutdown. Class I crossovers are interfering, so that one crossover prevents the formation of another nearby^11,12^. The number of Class I crossovers is limited by the level of HEI10 expression, synaptonemal complex and HCR1 protein phosphatase^13–19^. In contrast, non-interfering Class II crossovers are dependent on structure-specific nucleases including MUS81 and are strongly inhibited by DNA helicases, mainly FANCM and RECQ4 in plants^20–28^. Therefore, the number of crossovers in both pathways is kept relatively low, e.g., ∼8 Class I and ∼1 Class II crossovers per meiosis in *Arabidopsis thaliana*^29–31^.

The crossover distribution along the chromosomes is not uniform and is largely determined by the chromatin state^32–42^. DNA polymorphisms between homologous chromosomes can also affect crossover placement^43–48^. For example, the juxtaposition of heterozygous and homozygous regions stimulates crossover in the heterozygous region^49^. This effect is specific to the ZMM pathway and depends on MSH2^50,51^. MSH2 is a key subunit of complexes that detect base mismatches occurring both in somatic cells and during meiosis, when recombination is initiated in heterozygous regions^52–58^. On the other hand, a genome-wide comparison of the crossover distribution in *A. thaliana* hybrids and inbreds showed no significant differences^41^.

Here, we further explore the relationship between polymorphism and recombination, and explain the reasons for the apparent contradiction between chromosomal regions and genome-wide data^41,49,50^. To this end, we examine crossover formation in *msh2* mutants depending on the used crossover pathway. We applied both approaches based on genome-wide crossover maps as well as recombination analysis using fluorescent reporter lines (FTLs). We show that inactivation of MSH2 leads to a significant increase in Class II crossovers in *A. thaliana* hybrids. Furthermore, we demonstrate that polymorphism inhibits Class II crossover both in MSH2-dependent and independent manner. By analyzing crossovers using FTLs covering the entire arm of chromosome 3, we showed that the crossover distribution is very similar between inbreds and hybrids. However, a change in the heterozygous/homozygous pattern along the chromosome induces a dramatic Class I crossover redistribution across the heterozygous/homozygous boundary. We argue that the differences in the effects of MSH2 on the Class I and II crossover in response to interhomolog polymorphism are a consequence of distinct biological functions of the two pathways.

## Results

### MSH2 limits fertility of *fancm zip4* hybrids

Earlier studies from us and others have shown that while a mutation in the *FANCM* gene restores the fertility of the *zip4* mutant in inbreds, this effect is largely inhibited in hybrids^25,49,59^. To confirm that the phenotype observed in Col/L*er* hybrid plants was not caused by a L*er* genome-encoded factor, we generated a *zip4* mutation in the L*er fancm* background using CRISPR-Cas9 (Supplementary Fig. 1). The L*er fancm zip4* plants were fully fertile, similarly to their Col counterparts (Supplementary Fig. 1 and 2). Therefore, we hypothesized that the reduced fertility of the Col/L*er fancm zip4* hybrid is due to limited crossover recombination, which in turn is a consequence of the DNA polymorphism between homologues.

Interhomolog polymorphism is typically detected by binding of MSH2 heterodimers to mismatched bases^50,51,54,60–62^. Therefore, we decided to check if inactivation of *MSH2* would increase the fertility of Col/L*er fancm zip4* hybrids. We obtained triple mutants by backcrossing *msh2* to *fancm zip4* double mutants in both Col and L*er* backgrounds. We observed that the triple *msh2 fancm zip4* hybrid produced more seeds than the *fancm zip4* hybrid (Fig. 1a,b and Supplementary Tables 1-3). We also measured the seed set and silique length (Fig. 1c,d). While the *msh2 fancm zip4* plants showed a much lower number of seeds per silique (mean 20.1) than the wild type (mean 56.7), it was significantly higher compared to the *fancm zip4* mutant (mean 7.5, Welch’s *t-*test *P*=4.2×10^−5^) (Fig. 1c).

**Fig. 1.**
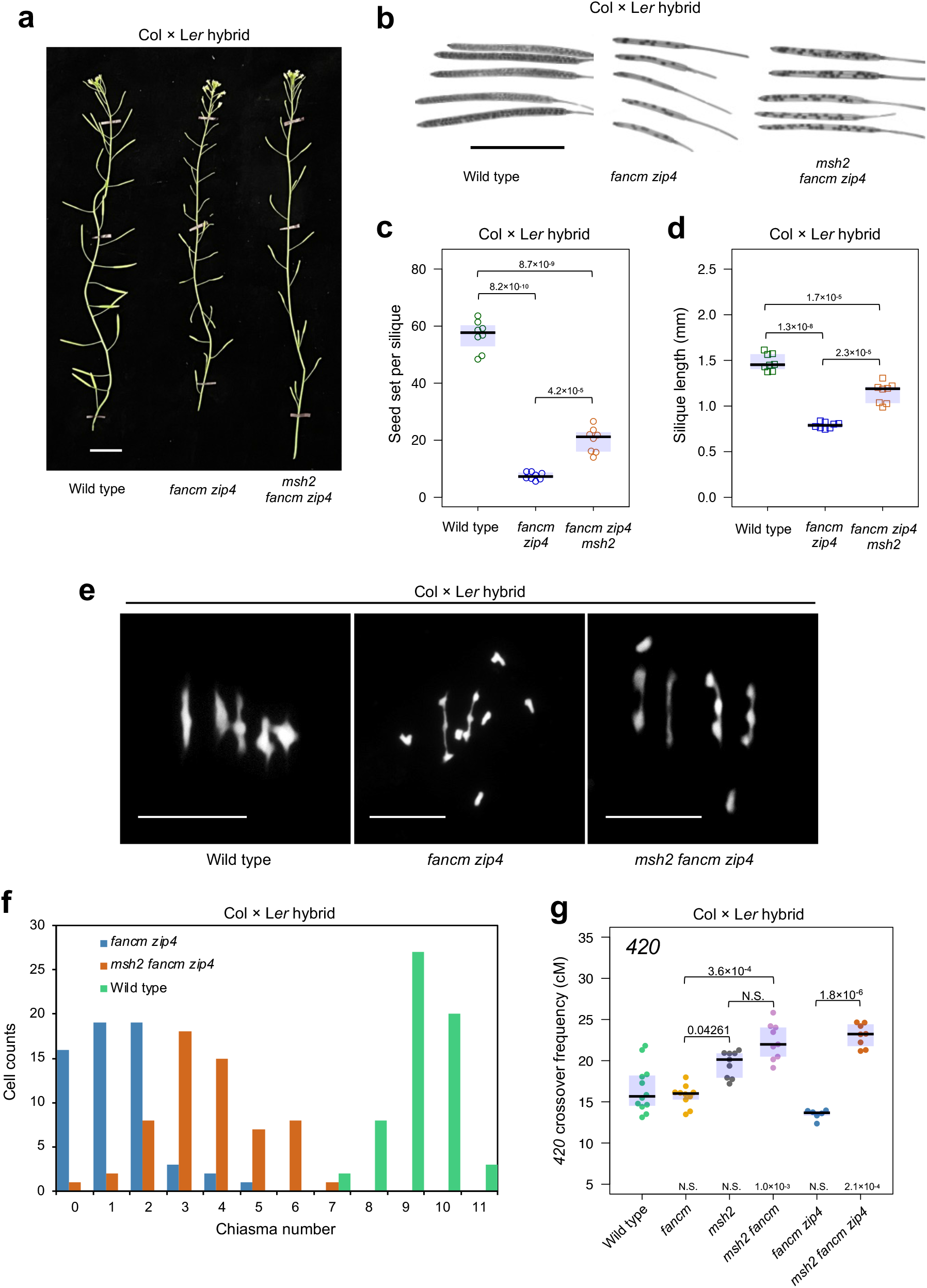
Inactivation of *MSH2* partially restores fertility in Col × L*er fancm zip4* hybrids. **a**, Representative images of wild-type, *fancm zip4* and *msh2 fancm zip4* Col × L*er* hybrids. Scale bar, 2 cm. **b**, Representative cleared siliques of wild-type, *fancm zip4* and *msh2 fancm zip4* Col × L*er* hybrids. Scale bar, 1 cm. **c-d**, Fertility assays in Col × L*er fancm zip4* and *msh2 fancm zip4* as assessed via seed set (**c**) and silique length (**d**). The center line of a boxplot indicates the mean; the upper and lower bounds indicate the 75th and 25th percentiles, respectively. Each dot represents a measurement from five siliques of one plant. Significance was assessed by Welch’s *t*-test. **e**, DAPI-stained metaphase I chromosome spreads from Col × L*er* male meiocytes in wild type, *fancm zip4* and *msh2 fancm zip4*. Scale bar, 5 μm. **f**, Quantification of chiasma count data from Col × L*er* male meiocytes in wild type, *fancm zip4* and *msh2 fancm zip4*. **g**, *420* crossover frequency (cM) in Col × L*er* hybrids of *fancm, msh2, msh2 fancm, fancm zip4* and *msh2 fancm zip4*. Significance was assessed by Kruskal-Wallis H test followed by Dunn’s test with Bonferroni correction. The center line of a boxplot indicates the mean; the upper and lower bounds indicate the 75th and 25th percentiles, respectively. Each dot represents a measurement from one individual.

To examine whether the increased fertility of *msh2 fancm zip4* was associated with an increase in the crossover number, we counted chiasmata in meiotic metaphase I. We observed that the *msh2 fancm zip4* mutant showed significantly more chiasmata than *fancm zip4* (mean 1.32 and 3.70, respectively; Mann-Whitney test *P*=1.72×10^−14^; Fig. 1e,f and Supplementary Table 4). Segregation of linked reporters expressing fluorescent proteins in seeds (fluorescent-tagged lines, FTLs) can be used to assess crossover frequency within defined chromosomal intervals^63,64^. Therefore, we backcrossed combinations of *msh2, fancm* and *zip4* mutants to the Col-*420* reporter line, and then obtained F_1_ plants via crosses to the corresponding mutants in L*er*. Since fertility in *zip4* is drastically reduced by crossover scarcity, the seeds produced in these mutants usually result from gametes that experienced more crossovers than the mutant average^30^. Thus, crossover measurements will be overestimated especially in the *fancm zip4* mutant. Nevertheless, comparing the *420* crossover frequency between *fancm zip4* and *msh2 fancm zip4* showed a statistically significant increase (Welch’s *t*-test *P*=8.6×10^−9^; Fig. 1g and Supplementary Table 5). Based on these observations, we concluded that MSH2 inhibits Class II crossovers in hybrids.

### MSH2 represses FANCM-dependent Class II crossovers

Although the chiasma number in *msh2 fancm zip4* is much lower than in wild type, crossover measurements over a single *420* interval show the opposite effect. This suggests that the chromosomal crossover distribution is different in *msh2 fancm zip4* than in the wild type. Therefore, we investigated the crossover on a genome-wide scale. We generated F_2_ populations from Col × L*er* crosses in the genetic background of *fancm zip4, msh2 fancm*, and *msh2 fancm zip4* mutants. We sequenced genomic DNA from between 172-199 individuals from each population and identified crossovers in each individual (Fig. 2a). Crossovers were defined as genotype switches along the chromosomes and were assigned to the midpoint between pairs of SNPs^65^. On this basis, we obtained data on the number and location of crossovers in the tested mutants, which we compared with the analogous data for wild-type and previously published *msh2* Col × L*er* crosses^50^ (Fig. 2). Both *fancm zip4* and *msh2 fancm zip4* showed lower crossover numbers than wild type (Welch’s *t*-test *P*=4.2×10^−14^ and *P*=4.6×10^−4^, respectively; Fig. 2b). Moreover, *msh2 fancm zip4* showed significantly higher crossover counts than *fancm zip4*, confirming our previous observations based on chiasma counts (*P*=2.2×10^−16^; Fig. 1f and 2b). We also observed elevated crossover numbers in *msh2 fancm* when compared to wild type (11.0 versus 8.0 crossovers per F_2_, *P*=1.1×10^−11^). Previous reports indicated that the crossover frequency in *fancm* Col × L*er* hybrids is not significantly higher than in wild type^25^. Therefore, it can be assumed that the inactivation of *MSH2* in *fancm* also leads to an increase in Class II crossover (Fig. 2b).

**Fig. 2.**
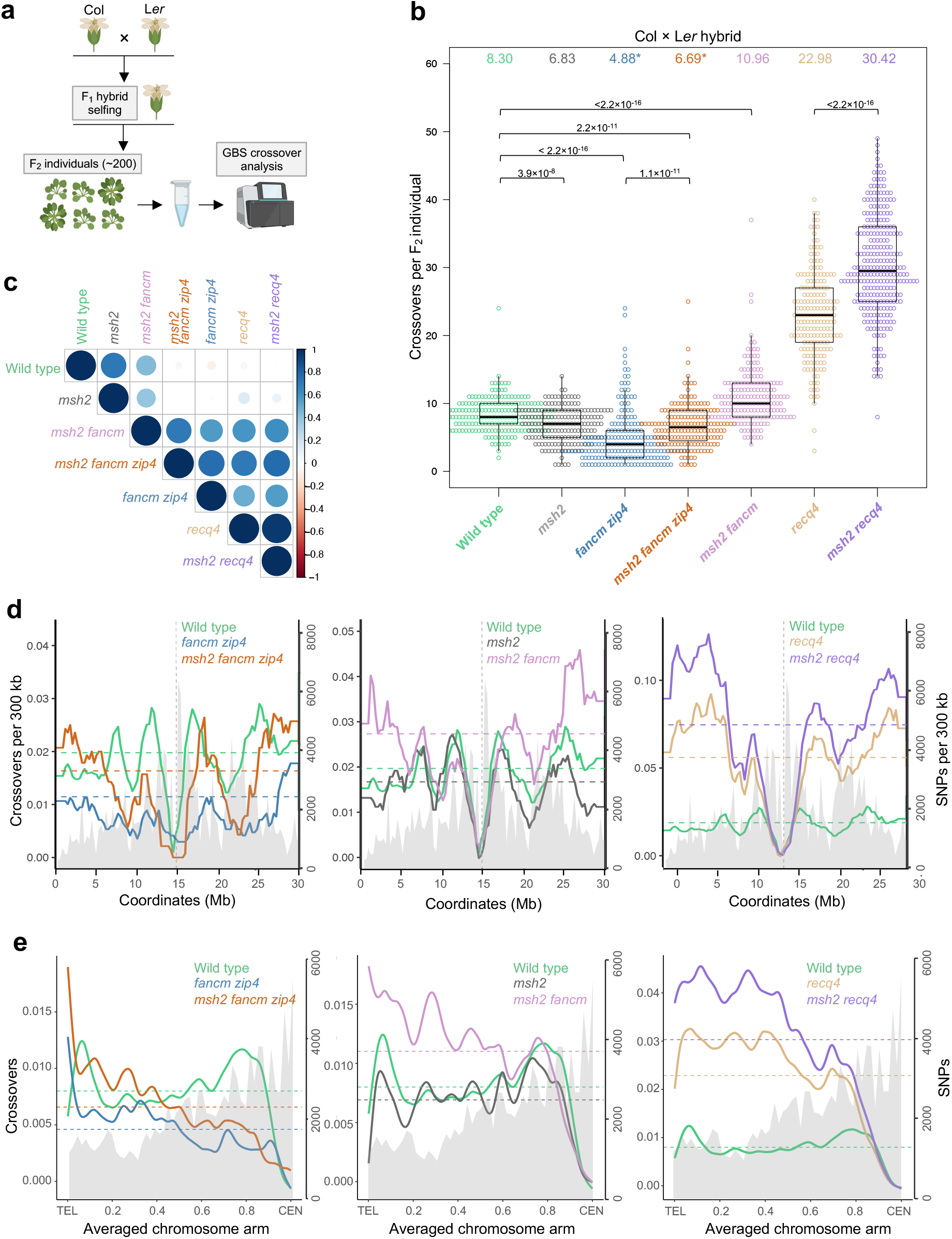
The effect on *MSH2* inactivation on crossover frequency and distribution in different mutant backgrounds. **a**, Diagram illustrating crossover mapping in F_1_ plants based on F_2_ individuals. Created with BioRender.com. **b**, The number of crossovers per F_2_ individual in the indicated populations. Significance was assessed by Kruskal-Wallis H test followed by Mann Whitney U test with Bonferroni correction. The center line of a boxplot indicates the mean; the upper and lower bounds indicate the 75th and 25th percentiles, respectively. The mean is indicated also on the top. **c**, The correlation coefficient matrices among genome-wide crossover distributions as calculated in 0.3 Mb adjacent windows. **d**, Crossovers per 300 kb per F_2_ plotted along the Arabidopsis chromosome 1. Mean values are shown by horizontal dashed lines. SNPs per 300 kb are plotted and shaded in gray. The position of centromere is indicated as vertical dashed line. **e**, Data as for **d**, but analysing crossovers along proportionally scaled chromosome arms, orientated from telomere (TEL) to centromere (CEN). **b-e**, Data for wild type, *msh2* and *recq4* from ref. 42, 50 and 66, respectively.

We then compared crossover distribution along the chromosomes (Fig. 2c-e). The profiles for *fancm zip4* and *msh2 fancm zip4* showed a reduced frequency of recombination in centromeres compared to wild type (Fig. 2c-e) and were strongly correlated (Spearman *Rho*=0.701, *P*<2.2×10^−16^; Fig. 2c). In turn, the crossover pattern for the *msh2 fancm* double mutant was an intermediate form between the pattern observed in the *msh2 fancm zip4* triple mutant (Spearman Rho=0.752, *P*<2.2×10^−16^; Fig. 2c) and the wild type (Rho=0.434, *P*<2.2×10^−16^; Fig. 2c). This likely reflects the similar activity of Class I and II crossover pathways in *msh2 fancm*. In both *msh2 fancm zip4* and *msh2 fancm* we observe an increase in crossovers at the chromosome ends compared to wild type. Together, our results indicate that *MSH2* inactivation increases Class II crossover frequency in the range blocked by the FANCM helicase in Col/L*er* hybrids.

### MSH2 represses RECQ4-dependent Class II crossovers

In contrast to the *fancm* mutation, in which an increase in recombination frequency is observed only in inbreds, the *recq4a recq4b* mutants (hereafter *recq4*) show elevated crossover level in both inbreds and hybrids^21,24,25^. However, the increase observed in the *recq4* hybrids is always lower than that observed in the fully homozygous system^25,66^. Therefore, we investigated whether MSH2 limits recombination in the *recq4* hybrid plants. For this purpose, we obtained the triple *msh2 recq4* mutants in the Col, L*er* and Col/L*er* backgrounds. Then, we sequenced 279 individuals of the F_2_ generation derived from this triple mutant and identified crossover sites. The *msh2 recq4* plants showed an average of 30 crossovers per individual, which is significantly more than the 23 crossovers observed in the *recq4* double mutant (Welch’s t-test P<2.2×10^−16^) (Fig. 2b). This suggests that MSH2 inhibits Class II crossover formation in the pathway controlled by RECQ4 in response to interhomolog polymorphism.

However, the crossover distribution was very similar and strongly correlated in *msh2 recq4* and *recq4* genotypes (Spearman *Rho*=0.958, P<2.2×10^−16^, Fig. 2b-d). While the increase in crossover frequency was observed primarily along the chromosome arms and at the chromosome ends, the difference between the two genotypes was small in the proximal regions (Fig. 2d,e). Therefore, we concluded that MSH2-dependent polymorphism detection is not the major reason for Class II crossover inhibition in pericentromeres.

### MSH2 has the opposite effect on the Class I and Class II crossovers in response to SNP density

To investigate to what extent SNP affects crossover activity in individual mutants, we applied an approach that is less dependent on chromosomal location. We divided the genome into 100 kb nonoverlapping windows for which we determined the SNP density and the crossover frequency. This resulted in 1191 windows, which we sorted according to the SNP density and divided them into 99 groups (Supplementary Fig. 4 and Supplementary Table 6). We plotted the relationship between SNP and crossover for wild type (Fig. 3a). This revealed a parabolic relationship, which was described before^50^. We then plotted the same relationship for the mutants, but after normalizing to wild type (by subtracting recombination frequency for wild type from the given mutant for each SNP density group).

**Fig. 3.**
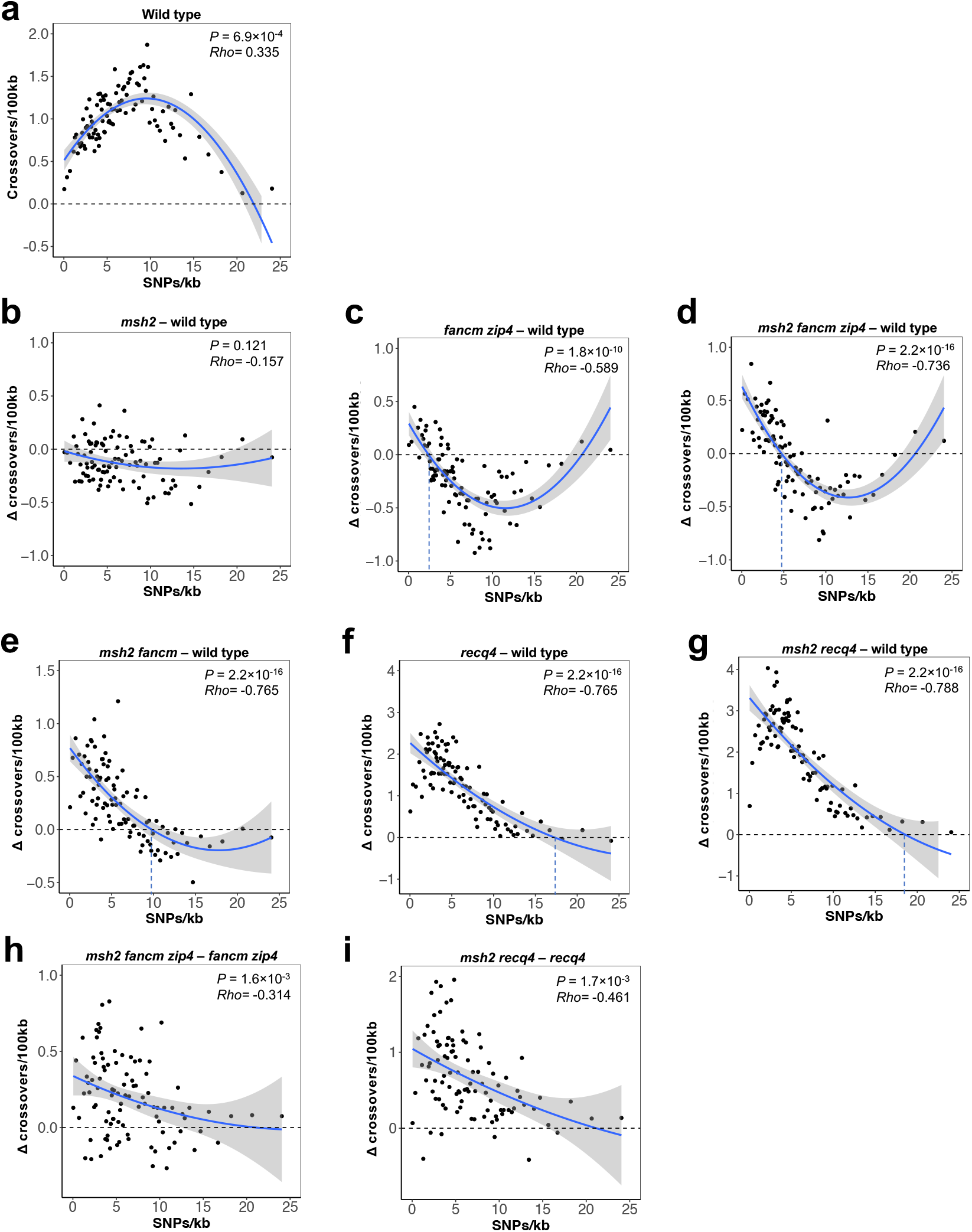
Relationship between SNP density and crossover frequency in different mutant backgrounds. **a**, Crossover frequency as a function of SNP density (SNPs/kb) in wild-type Col/L*er* plants. Crossovers normalized by the number of F_2_ individuals and SNP density in 100 kilobase (kb) adjacent windows were calculated for each population and ranked into percentiles according to SNP density. **b-g**, The difference between crossover frequency in a mutant and wild type (Δ cM) was plotted in relationship to SNP density (SNPs/kb). **h-i**, The difference between crossover frequency in a multiple mutant carrying *msh2* mutation and its counterpart with functional *MSH2* (Δ cM) was plotted in relationship to SNP density (SNPs/kb). The Spearman’s rank correlation coefficient (*Rho*) between SNP density and crossover frequency (**a**) or crossover frequency difference (**b-i**) is printed inset. Trend lines were fitted in ggplot using a generalized additive model (GAM) with the formula y ∼ poly(x,2). Data for wild type, *msh2* and *recq4* from ref. 42, 50 and 66, respectively.

In the *msh2* mutant, we observed a slight decrease in the crossover frequency in almost all SNP density groups (Fig. 3b). However, both Class I and Class II crossovers are formed in the *msh2* mutant, thus conclusions must be drawn with caution. Stronger reductions were observed in the *fancm zip4* and *msh2 fancm zip4* mutants, but only for groups with a high density of SNPs: below 2.5 (*fancm zip4*) and 5 SNPs/kb (*msh2 fancm zip4*) there is a noticeable increase in the crossover frequency (Fig. 3c,d). Despite the inactive ZMM pathway in both mutants (*zip4* mutation), we did not observe a decrease in the crossover frequency in groups with a high SNP density (>18 SNPs/kb), because these regions are practically recombinantly inactive in the wild type (Fig. 3a). The *msh2 fancm* mutant shows a negatively proportional to SNP density increase in crossover frequency in groups below ∼10 SNPs/kb and a decrease above this limit (Fig. 3e). In both *recq4* and *msh2 recq4* we observed a negatively proportional increase in the crossover frequency, which reached the wild-type level at approximately >17 and 18 SNP/kb, respectively (Fig. 3f,g). Although the plots for these two mutants look similar, the different slope of the correlation lines is evident.

To better explore the difference made by *msh2* mutation, we subtracted *recq4* crossover frequency from *msh2 recq4* frequency for each group (Fig. 3i). This showed a proportional inverse relationship between the SNP/kb density and the increase in recombination frequency being a consequence of *MSH2* inactivation (*Rho*=-0.461, *P*=1.74×10^−6^). The effect of MSH2 inactivation is especially strong in regions with a relatively low SNP density (<6 SNPs/kb), where it leads to an increase in recombination frequency by up to 2 cM/100kb (Fig. 3i). The same analysis for *msh2 fancm zip4* showed a similar though weaker correlation (*Rho*=-0.314, *P*=0.0016; Fig. 3h).

A comparison of the analyzes for *msh2* – WT (Fig. 3b) vs. *msh2 fancm zip4* – *fancm zip4* and *msh2 recq4* – *recq4* (Fig. 3h,i) suggests that MSH2 affects crossovers differently in both pathways: In the ZMM pathway, which is responsible for Class I crossovers, MSH2 complexes stimulate recombination, while in pathways leading to the formation of Class II crossovers, MSH2 inhibits recombination. The MSH2 inhibition effect, however, is much stronger in regions with relatively low polymorphism densities.

### Local effects of DNA polymorphism on Class II crossover

In our previous studies, we observed that heterozygous regions show elevated crossover numbers when they are adjacent to homozygous regions on the same chromosome^49^. This effect depends on MSH2 heterodimers detecting interhomolog polymorphisms^50^. Interestingly, the heterozygosity/homozygosity juxtaposition effect applies only to Class I crossovers: with blocked ZMM pathway via *zip4* mutation and increased activity of the non-interfering pathway via *fancm* mutation, we observed an opposite trend with a drastic decrease in crossover frequency in the heterozygous region. To investigate the genetic basis of this distinct response to polymorphism, we measured crossover frequency in plants with different heterozygosity contexts. For this purpose, we used Col/Ct recombinant lines, which differ in the heterozygosity pattern on chromosome 3 and allow us to measure the crossover frequency in the *420* interval, also located on chromosome 3^49^.

Four Col/Ct polymorphism configurations were used: (i) “HOM-HOM” that are Col/Col inbred throughout the genome, (ii) “HET-HET” that are Col/Ct hybrids throughout the genome, (iii) “HET-HOM” where the *420* region is Col/Ct heterozygous and the remainder of chromosome 3 is Col/Col homozygous and (iv) “HOM-HET” where *420* is Col/Col homozygous and the remainder of chromosome 3 is Col/Ct heterozygous (Fig. 4a).

**Fig. 4.**
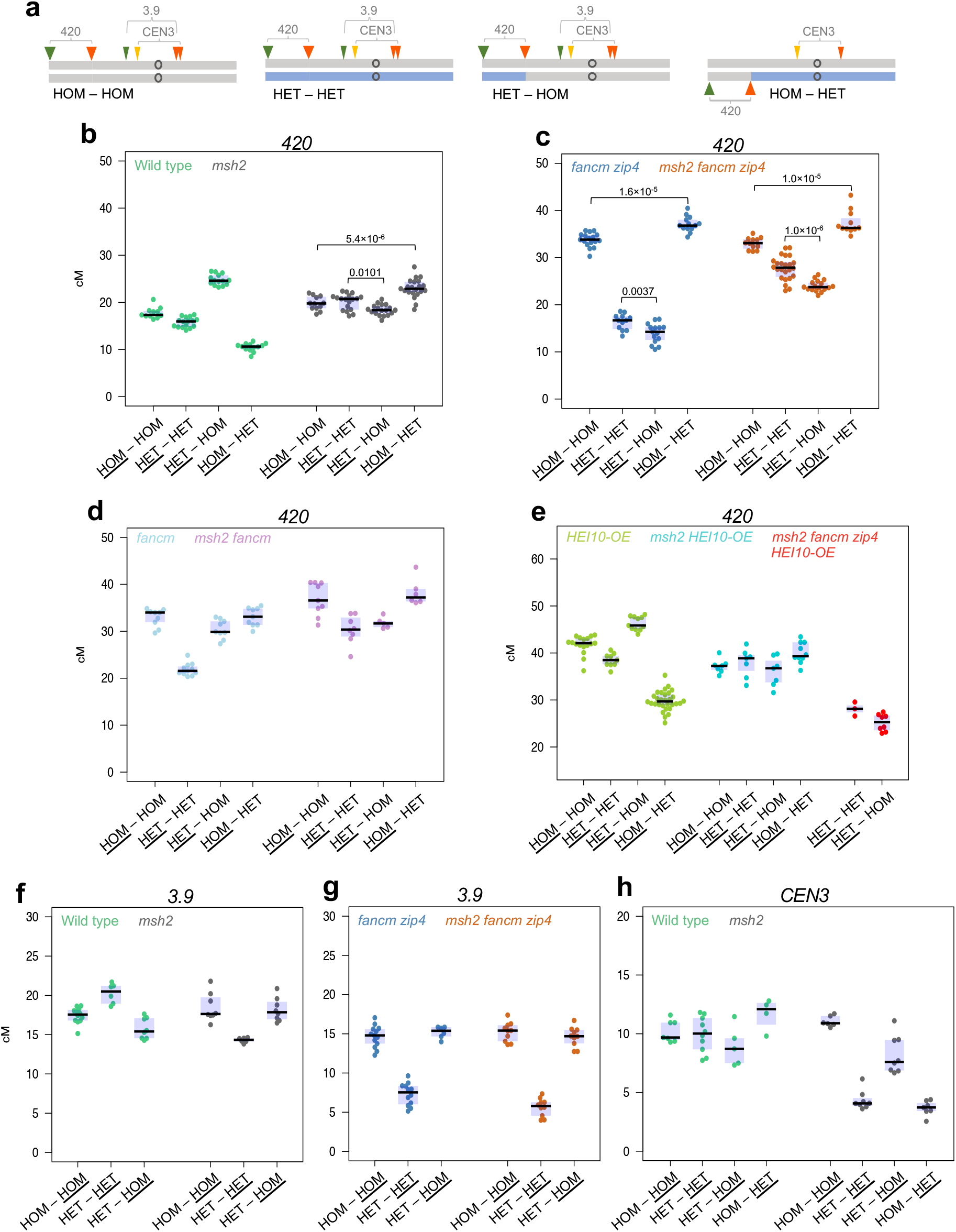
MSH2-dependent and independent crossover redistribution in response to heterozygosity pattern. **a**, Ideograms of chromosome 3 in lines differing in heterozygosity pattern. Gray corresponds to Col while blue corresponds to Ct genotype. Location of fluorescent reporters defining three different intervals (*420, 3*.*9* and *CEN3*) are indicated together, for simplicity. **b**, *420* crossover frequency (cM) in the HOM-HOM, HET-HET, HET-HOM, HOM-HET genotypes shown in **a**, in either wild type or *msh2*. Underlined is the homo- or heterozygosity state of the *420* interval. The center line of a boxplot indicates the mean; the upper and lower bounds indicate the 75th and 25th percentiles, respectively. Each dot represents a measurement from one individual. Welch’s t-test was used to calculate statistical significance. **c**, as in **b**, but for *fancm zip4* and *msh2 fancm zip4*. **d**, as in **b**, but for *fancm* and *msh2 fancm*. **e**, as in **b**, but for *HEI10-OE, msh2 HEI10-OE* and *msh2 fancm zip4 HEI10-OE* (only HET-HET and HET-HOM genotypes). **f**, *3*.*9* crossover frequency (cM) in the HOM-HOM, HET-HET and HET-HOM genotypes shown in **a**, in either wild type or *msh2*. **g**, as in **f**, but for *fancm zip4* and *msh2 fancm zip4*. Underlined is the homo- or heterozygosity state of the *3*.*9* interval. **h**, *CEN3* crossover frequency (cM) in the HOM-HOM, HET-HET, HET-HOM, HOM-HET genotypes shown in **a**, in either wild type or *msh2*. Underlined is the homo- or heterozygosity state of the *CEN3* interval. Each dot represents a measurement from a pool of 5-8 individuals. Data in **b** from ref. 50.

We observed a high recombination frequency in the *fancm zip4* mutant whenever the *420* region was homozygous and very low whenever *420* was heterozygous (Fig. 4b,c and Supplementary Table 7), which is consistent with previous observations^49^. When we additionally inactivated the *MSH2* gene in these lines, obtaining the triple mutants *msh2 fancm zip4*, the *420* crossover frequency for HET-HOM (24.0 cM) and HET-HET (27.8 cM) increases substantially compared to their counterparts in the *fancm zip4* background (Welch’s *t*-test *P*=1.1×10^−13^ and *P*<2.2×10^−16^, respectively), while not changing for the HOM-HET and HOM-HOM lines. The increase in *420* crossover frequency in *msh2 fancm zip4* HET-HET is also significant compared to its wild-type HET-HET counterpart (Welch’s *t*-test *P*<2.2×10^−16^). However, both the HET-HET and HET-HOM lines in *msh2 fancm zip4* remain colder in *420* than HOM-HOM and HOM-HET in *msh2 fancm zip4*, indicating that some genetic or epigenetic factor still limits the formation of crossovers in heterozygous state (Fig. 4c). This shows a MSH2-independent effect of DNA polymorphism on Class II crossover formation.

Interestingly, the *420* crossover frequency was significantly higher in HOM-HET versus HOM-HOM (*P* values indicated on Fig. 4b,c) both in *msh2* and in *fancm zip4* and in *msh2 fancm zip4*, even though the *420* interval was homozygous in both cases. Conversely, the *420* crossover frequency is lower in HET-HOM compared to HET-HET (Fig. 4b,c), while in both lines the *420* interval was heterozygous. This may suggest the existence of a weak modifier of the chromosomal crossover distribution differentiating Col and Ct genotypes, located on chromosome 3 beyond the *420* region.

We created combinations carrying only *fancm* or *msh2 fancm* mutations, thus having both active Class I and boosted Class II crossovers (Fig. 4d). As expected, *420* crossover frequencies for the juxtaposition lines in *fancm* are compilations of values observed in wild type and *fancm zip4* (Fig. 4d and Supplementary Table 7). Similarly, the values measured for *msh2 fancm* appear to be the resultant of the values measured for *msh2* and *msh2 fancm zip4* (Fig. 4d and Supplementary Table 7).

Previously, we showed that overexpression of *HEI10* preserved the juxtaposition effect (Fig. 4e)^50^. When we combined *HEI10* overexpression with *MSH2* inactivation, this effect disappeared following the trends observed in the single *msh2* mutant, though all the lines showed increases in *420* crossover rate (compare Fig. 4b and 4e; Supplementary Table 8). In addition, *HEI10* overexpression was not able to increase crossover frequency above the level observed in *msh2 fancm zip4* (Fig. 4d). Altogether these results confirm that HEI10 has no role in crossover formation outside the ZMM pathway.

### Pericentromeric region of chromosome 3 shows a strong MSH2-independent crossover inhibition when Col/Ct heterozygous

We decided to check if the mutation in *MSH2* is able to increase *fancm zip4* recombination also in much more polymorphic pericentromeric regions. To this end, we backcrossed the *msh2, fancm zip4* and *msh2 fancm zip4* mutations to the CTL-*3*.*9* line (hereafter *3*.*9*), in which crossover-measuring interval spans the centromere of chromosome 3^67^. We obtained HOM-HOM (Col/Col inbred), HET-HET (Col/Ct hybrid) and HET-HOM (Col/Ct hybrid in *420* but Col/Col inbred in *3*.*9*) lines in mutant and wild type backgrounds (Fig. 4a and Supplementary Table 9). In wild-type *3*.*9* lines, recombination was higher in hybrids than in inbreds (20.24 and 17.44 cM, respectively; Welch’s *t*-test *P*=1.5×10^−3^; Fig. 4f), which is consistent with previous observations that Col/Col inbreds have a relatively low crossover frequency in the centromere regions^41^. HET-HOM lines, where the *3*.*9* interval is in the homozygous region while the *420* chromosome fragment is heterozygous, showed a decrease of *3*.*9* crossover frequency compared to the inbred (15.74 cM, *P*=9.6×10^−3^; Fig. 4f). This decrease is due to crossover redistribution from the homozygous region spanning *3*.*9* interval, to the heterozygous region in subtelomeric region^49^. In the *msh2* background, we observed that the *3*.*9* crossover frequency is significantly lower in the hybrid than in the inbred (14.30 and 18.47 cM, *P*=2.8×10^−4^), whereas HOM-HOM was not different from HET-HOM (Fig. 4f). This shows that when MSH2 is inactive, crossover recombination is inhibited at pericentromeres in the heterozygous state. One possibility is that this is triggered by a high number of mismatches formed during strand invasion, because pericentromeres are substantially more polymorphic than distal regions^68^. Other explanations are also possible, for instance potential differences in chromatin states between Col and Ct accessions within pericentromeres^69^.

For any of the heterozygosity combinations in the *fancm zip4* double and *msh2 fancm zip4* triple mutants tested, we did not observe a significant increase in *3*.*9* recombination relative to their wild-type counterparts (Fig. 4f,g). In both multiple mutants, recombination rate in the lines differing in the pattern of heterozygosity is very similar to that observed in the *msh2* mutant, but the decrease in HET-HET relative to HOM-HOM and HET-HOM is deepened. This confirms previous observations that a mutation in the *FANCM* gene is unable to restore recombination in the heterozygous pericentromeric regions when the ZMM pathway is off^49^. However, it also shows that this inability to act in polymorphic regions is mainly MSH2 independent. These results are consistent with our genome-wide data for the Col × L*er* crosses (Fig. 2d,e).

We confirmed the results for *msh2* in the *CEN3* interval, which also has a pericentromeric location (Fig. 4a and Supplementary Table 10). In the case of *CEN3*, recombination is measured by segregation of fluorescent reporters expressed in the pollen, therefore it is specific for male meiosis^64,70^. These results are consistent with those observed for *3*.*9*, except that the decrease in *msh2* HET-HET recombination relative to *msh2* HOM-HOM is even more drastic (Fig. 4h; 4.3 and 11.06 cM, respectively *P* = 4.6 × 10^−10^). This may be related to a more distal crossover location in Arabidopsis male meiosis than in female meiosis^71,72^.

Altogether these results show that MSH2 is effective in stimulating Class I crossovers in pericentromeric regions when heterozygous. In contrast, non-interfering Class II crossovers in the FANCM-regulated repair pathway cannot be formed in the polymorphic regions near the centromere, whether MSH2 is active or not.

### Crossover redistribution along the chromosome arm in response to heterozygosity pattern

One of the advantages of using reporter systems for measuring crossover frequency is that they allow to study recombination in both hybrid and inbred contexts. We decided to use this property to investigate whether interhomolog polymorphism-dependent crossover redistribution (heterozygosity/homozygosity juxtaposition effect) is uniform along heterozygous regions. For this purpose, we used a set of eight additional FTLs covering the longer arm of chromosome 3 (Fig. 5a and Supplementary Table 11)^67^. We previously obtained lines with a changed pattern of heterozygosity along this chromosome arm (Fig. 5a)^49,50^.

**Fig. 5.**
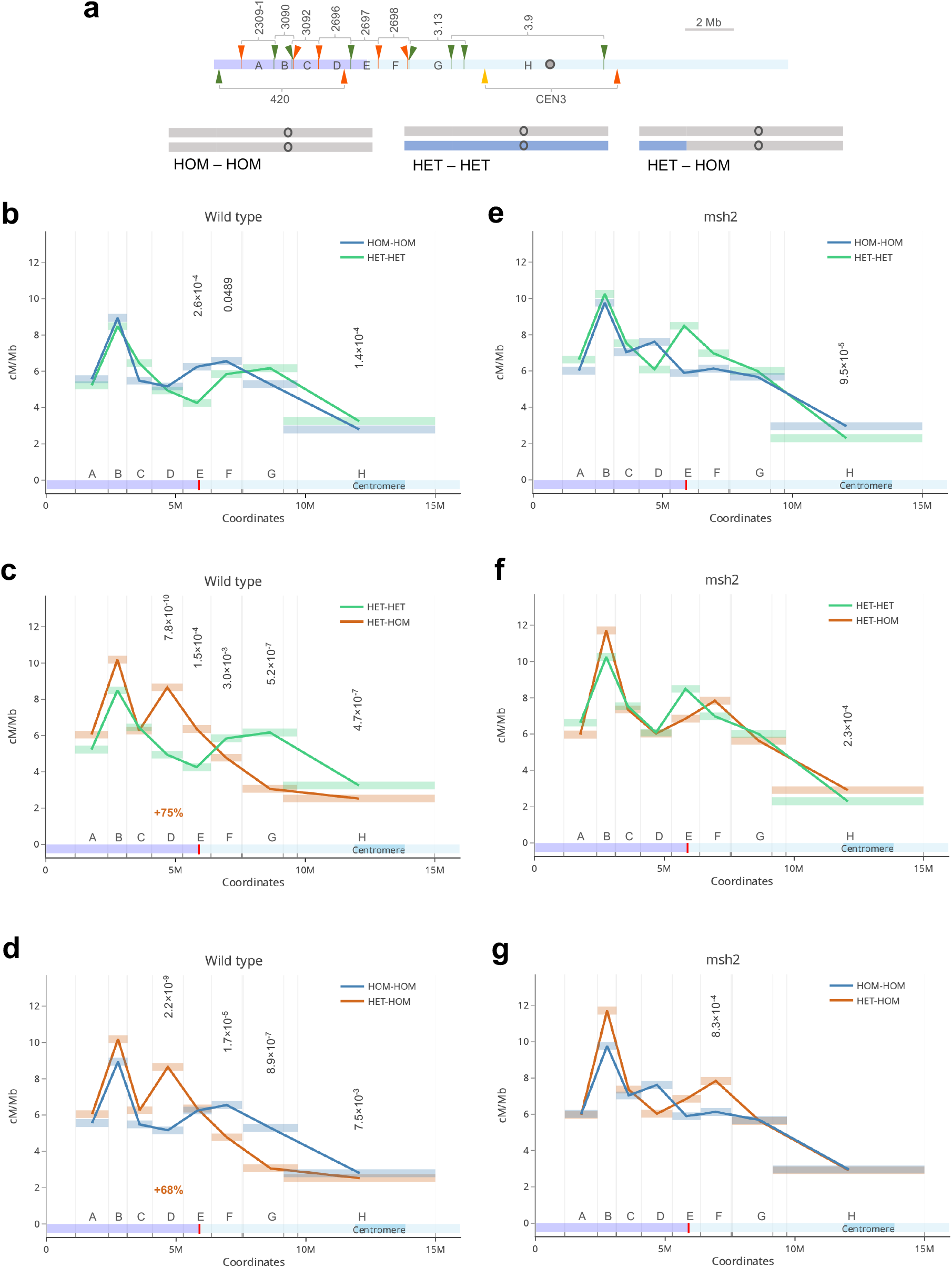
Crossover distribution across the chromosome arm in response to polymorphism. **a**, (upper panel) Location of eight FTL intervals across the Arabidopsis chromosome 3 shown in addition to *420* and *CEN3* intervals. Fluorescent reporters are indicated by tick marks and arrowheads with colors corresponding to eGFP (green), dsRed (red) and eYFP (yellow). Capital letters ‘A’ – ‘H’ were used instead of the original FTL names (indicated on the top), for simplicity. The violet and light blue shading represents ‘HET’ and ‘HOM’ regions in HET-HOM line, respectively. Genetically defined centromere^95^ is indicated as gray circle. (lower panel) Ideograms of chromosome 3 showing heterozygosity pattern in lines used in **b-g**. Gray corresponds to Col while blue corresponds to Ct genotype. **b-d**, Comparison of mean crossover frequency (cM/Mb) in eight intervals along the chromosome arm in inbred (HOM-HOM) vs. hybrid (HET-HET) (**b**), hybrid vs. recombinant line (HET-HOM) (**c**) and inbred vs. recombinant line (**d**). Rectangles represent the length of the intervals, interval names as in **a**. One-way ANOVA with Tukey HSD was used to calculate statistical significance. Not significant values were not shown. The relative percentage increase in the interval D crossover frequency is indicated in red. **e-g**, as in **b-d**, but for the *msh2* mutant.

We first compared the distribution of recombination along the arm in hybrids (HET-HET) versus inbreds (HOM-HOM) (Fig. 5b, Supplementary Fig. 6 and Supplementary Table 12). We observed a slight increase in HET-HET recombination frequency in the proximal regions with a concomitant reduction in the arm (Fig. 5b). This modification corresponds well to the effects recently described in the genome-wide comparison of Col/Col inbreds with Col/Ler hybrids^41^. In the HET-HOM line, where the distal fragment of the chromosome was heterozygous, while the rest of the chromosome was homozygous, there was a strong increase in the crossover frequency in the HET region compared to both inbreds and hybrids, but only at the very HET-HOM border (intervals D and partly E, which is ∼2Mb from the HOM/HET border; Fig. 5c,d, Supplementary Fig. 6 and Supplementary Table 12). As expected, this increase was at the expense of a decrease in the neighboring HOM region. Interestingly, however, a statistically significant decrease was observed not only in the immediate vicinity of the HET/HOM border, but over a longer section of the chromosome, including the “H’ pericentromeric interval, which starts 3.47 Mb from the HET-HOM breakpoint (Fig. 5c,d).

To investigate how MSH2 complexes influence crossover chromosomal distributions in different heterozygosity contexts, we backcrossed the eight FTLs to the *msh2* mutant. Comparisons between *msh2* inbreds and hybrids revealed that only the pericentromeric interval ‘H’ showed significant differences in recombination frequency (*P*=9.5×10^−5^; Fig. 5e, Supplementary Fig. 6 and Supplementary Table 12). This may be due to potential structural variations between Col and Ct within pericentromeres. Crossover distribution does not differ in HET-HOM from HET-HET outside this region (Fig. 5f). However, we observed a significant difference between HET-HOM and HOM-HOM in the ‘F’ interval, which is the first homozygous segment from the HET-HOM border. Contrary to wild type, we observed an increase in crossover frequency in the HET-HOM line within this interval (*P* = 8.3 × 10^−4^; Fig. 5g). Overall, line-to-line differences were much smaller in the *msh2* background than in the wild type. In conclusion, our analyzes confirm that the chromosomal crossover distribution is similar in hybrids and inbreds, as recently described by genome-wide analysis^41^. Only the introduction of a heterozygous chromosomal segment into an otherwise homozygous chromosome causes a drastic local redistribution of recombination, and this effect is MSH2 dependent. This demonstrates that MSH2 has a stimulating effect on Class I crossover in response to polymorphism primarily in the immediate boundary between heterozygous and homozygous regions. However, in the case of a gradual decrease in the polymorphism density, which exists in hybrids, the effect is hardly noticeable.

### Crossover interference is maintained both in *msh2* hybrids and inbreds

Linked fluorescent reporters expressed in pollen can be used to locally measure crossover interference represented as coefficient of coexistence (1 – CoC). This provides a unique opportunity to compare the strength of interference both in hybrids and inbreds, either in a wild-type or *msh2* mutant backgrounds. Consistent with previous reports^49^, we observed that interference is stronger in hybrids than in inbreds (Supplementary Fig. 5 and Supplementary Tables 13,14). In contrast, interference remained unchanged in both *msh2* inbreds and hybrids relative to the values measured for the wild type (Supplementary Fig. 5). This shows that although the presence of interhomolog polymorphism increases the strength of crossover interference, this effect is not dependent on the detection of mismatches involving MSH2 complexes.

## Discussion

In *A. thaliana*, the ZMM pathway that creates Class I crossover is dominant, while Class II crossovers are rare and therefore unable to secure the proper course of meiosis^73,74^. To investigate the potential differences between the effect of polymorphism on both crossover classes, we used mutants in which Class II is hyperactivated (*fancm* and *recq4*), also in conjunction with the *zip4* mutation that completely disables Class I crossovers^21,24,25^. Previous studies have shown that while the combination of *fancm* and *zip4* mutations leads to normal fertility of inbreds, repeating this experiment in hybrids results in sterile plants^25^. An intuitive explanation for this is that the interhomolog polymorphism blocks Class II crossovers^6,25^. Indeed, the combination of the *fancm zip4* mutation with the *msh2* mutation resulted in an increase in fertility, but not to the level observed in the wild type (Fig. 1). Therefore, we performed genome-wide crossover analysis for *fancm zip4, msh2 fancm zip4, recq4* and *msh2 recq4* Col/L*er* hybrids. This revealed that *msh2* mutations invariably leads to increases in Class II crossover numbers (Fig. 2), indicating that MSH2, presumably via binding mismatches during strand invasion, blocks Class II crossover either by recruiting FANCM and RECQ4 or by limiting MUS81 activity (Fig. 6a). In budding yeast, the MSH2 complexes recruit SGS1, the RECQ4 homologue, to heteroduplexes, thus prevents crossover repair in somatic cells^75,76^. Interestingly, in both *fancm* and *recq4*, the *msh2* mutation is unable to force Class II crossovers in regions with high polymorphism density (Fig. 3). This suggests that DNA polymorphism also inhibits Class II crossovers in an MSH2-independent manner, possibly limiting the stability of D-loops during strand invasion.

**Fig. 6.**
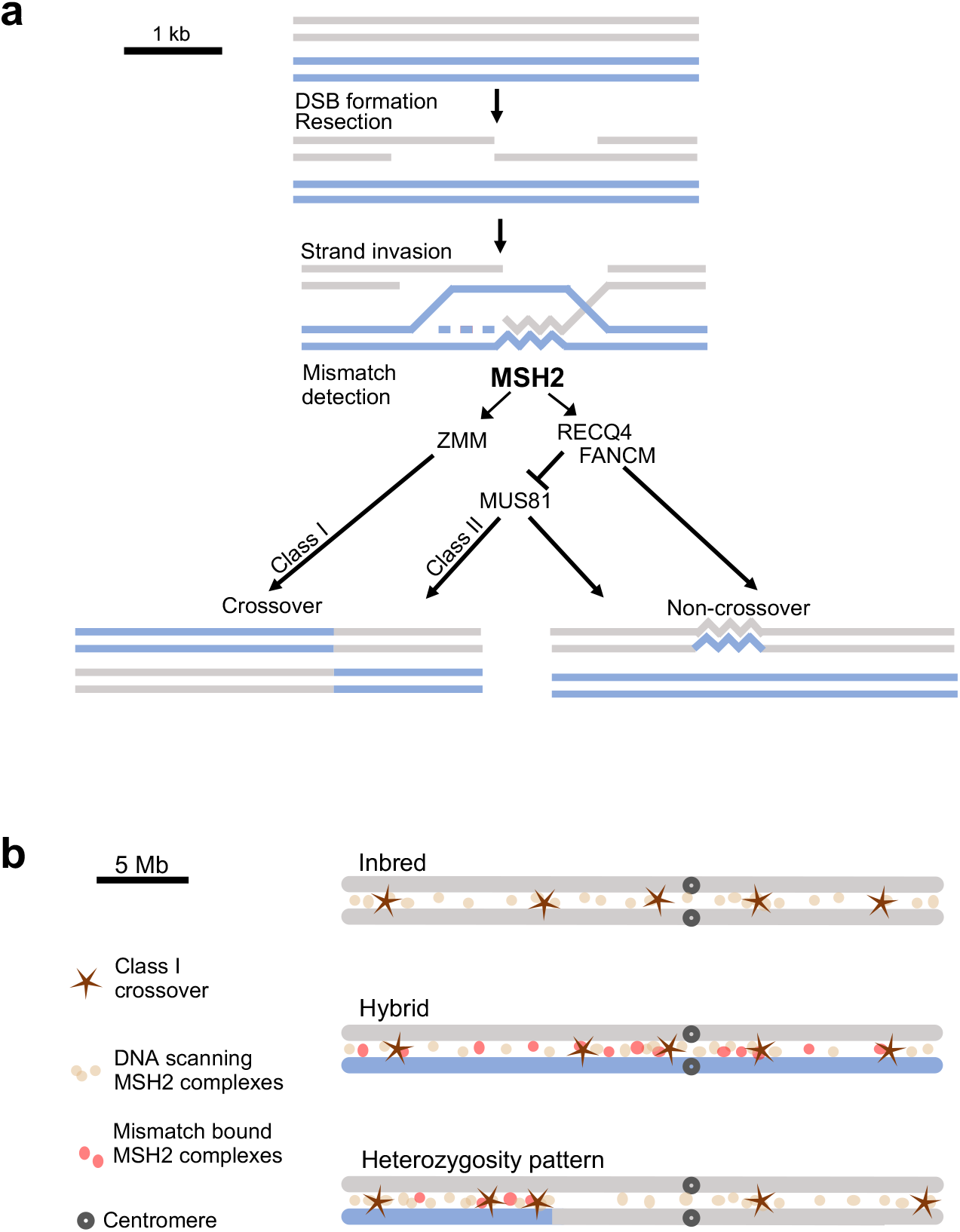
Models showing the impact of DNA polymorphism on crossover formation during Arabidopsis meiosis. **a**, Two recombining homologous DNA molecules are depicted in gray and blue, over a region of several kilobases. Following DSB formation on the gray molecule, resection occurs to form 3’-single-stranded DNA. This ssDNA undergoes strand invasion into the homologous molecule, forming a displacement loop. MSH2 heterodimers detect mismatches at the invasion site. Two scenarios are proposed for the situation when mismatches are detected: (i) MSH2 promotes ZMM pathway leading to Class I crossover, or (ii) MSH2 recruits or stimulates DNA helicases, including FANCM and RECQ4, resulting in D-loop displacement and non-crossover repair. In the absence of mismatches or MSH2, MUS81 endonuclease repairs the DSB via Class II crossover or noncrossover. Alternatively, MSH2 can directly stimulate MUS81-dependent crossover formation (not shown). **b**, Two recombining homologues are depicted with gray and blue colors representing sequence divergence. MSH2 complexes scan DNA to detect mismatches in heteroduplexes. In inbreds, when there are no mismatches, the position of crossovers is determined mainly by the chromatin structure. In hybrids, the mismatches along the entire chromosome length trigger a fairly even distribution of mismatch bound MSH2 complexes, which, combined with interference, also results in Class I crossover placement determined mainly by chromatin. However, the presence of a single heterozygous region on an otherwise homozygous chromosome results in a local concentration of mismatch bound MSH2 complexes that stimulate Class I crossover in the heterozygous region.

In addition, we investigated how MSH2 inactivation affects the distribution of Class II crossovers when the same chromosome fragment is either homozygous or heterozygous. This allows elimination of other effects besides polymorphism, including DNA methylation and chromatin state. We confirmed that FANCM-associated Class II crossovers are strongly repressed when the region is heterozygous and *msh2* mutation only partially restores recombination activity (Fig. 4). At the centromere region in the heterozygous state, *msh2* is unable to increase Class II crossover frequency, remaining two-fold lower than when this region is homozygous (Fig. 4f-h). Again, this confirms our observation that polymorphism inhibits Class II crossovers also in an MSH2-independent manner.

Although we do not have mutants in which only the ZMM pathway is active, the Class I crossovers predominate both in wild type and in *msh2*^2,3,8,29^. Our genome-wide analyzes showed a limited effect of polymorphism on Class I crossover distribution (Fig. 2-3, ref. ^50^), which is consistent with previous observations^41^. Using reporter lines covering the entire arm of chromosome 3 we also showed that the differences between inbreds and hybrids are small (Fig. 5). However, the juxtaposition of the heterozygous and homozygous regions causes a MSH2-dependent redistribution of the Class I crossover towards the former, regardless of the chromosomal location^49,50^ (Fig. 4b,f,h, Fig. 5). Recently, we showed that this applies not only to large chromosomal regions, but also to individual recombination hotspots^51^. This proves the stimulating role of MSH2 on Class I crossover in response to interhomolog polymorphism, primarily in the situation of a different pattern of polymorphism along the chromosome. In line with that in vitro assays show that yeast MSH2 complexes stimulate MLH1-MLH3, the major endonuclease in ZMM pathway^77^.

However, Arabidopsis hybrids naturally have regions of high and low polymorphism density (Fig. 2d, Supplementary Fig. 3), so why don’t we see significant crossover redistribution between inbreds and hybrids? A possible explanation is the limited availability of MSH2 heterodimers. In hybrids, the SNP density is so high that not all mismatches can be bound simultaneously by MSH2 complexes. The crossover interference remains at a similar level in wild type and *msh2*, so tight crossover control is maintained (Supplementary Fig. 5, ref. ^50^). As a consequence, other factors, such as chromatin structure or DNA methylation, become dominant, while polymorphism-dependent changes in crossover distribution are small (Fig. 5b, ref. ^41^). On the contrary, when only a single chromosomal region is heterozygous and the rest of the genome is homozygous, MSH2 saturation occurs in this region and strong local crossover stimulation occurs (Fig. 6b). Similarly, in a homozygous region on an otherwise heterozygous chromosome, the absence of MSH2 results in a decrease in crossovers. This hypothesis is confirmed by the fact that we practically do not observe differences between the tested lines in *msh2* (Fig. 5e-g).

Our results show that the two crossover pathways exhibit dramatic differences in response to interhomolog polymorphisms and that the effect of MSH2 complexes on these pathways is opposite (Fig. 6a). Observed differences are likely related to different biological functions of the two pathways: The ZMM pathway leading to the formation of Class I crossover is dedicated exclusively to meiotic recombination during gamete formation. As sexual reproduction involves mixing genetic material from non-identical parental individuals, the detection of polymorphisms in this pathway cannot block crossovers. Moreover, targeting recombination to regions that differ between individuals allows for the formation of new allelic combinations. This is different for the Class II crossovers, which are formed via pathways also involved in DNA repair in somatic cells, where recombination between non-identical sequences threatens genome stability^78,79^. Therefore, the detection of mismatches blocks Class II crossover repair, leading invariably to heteroduplex rejection and noncrossover repair. We have shown here that this is done with the active participation of MSH2 complexes.

Lack of meiotic recombination is one of the causes of infertility of distant hybrids^80,81^. Our results indicate that the inactivation of the mismatch detection system in combination with the inactivation of anti-recombination factors, enables an increase in the frequency of Class II crossover in polymorphic regions. Similar effect was recently reported in hybrids between diverged *Saccharomyces* species^82^. The use of such an approach may allow to break reproductive isolation between plant species, where the limitation is DNA divergence, which prevents the exchange of genetic material.

## Methods

### Growth conditions and plant material

Plants were grown in controlled environment chambers at 21°C with long day 16/8h light/dark photoperiods with 70% humidity and 150-μmol light intensity. Prior to germination seeds were kept for 48h in the dark at 4°C to stratify germination.

The *msh2-1* T-DNA insertion line (SALK_002708) and Arabidopsis accessions Col, Ler and Ct were obtained from the Nottingham Arabidopsis Stock Centre (NASC). The *msh2-2, msh2-3, msh2-4, msh2-5* (in Col/Ct recombinant lines) and *msh2-6* (in L*er*-0) deletion mutants were previously generated in our laboratory via CRISPR/Cas9 mutagenesis^50,51^, as well as Ler *zip4-3* mutant generated in this study. The *HEI10-OE* line corresponds to transgenic line “C2”, previously described in (^14^). For measuring recombination frequency fluorescence-tagged lines were used: *420* (kindly provided by Avraham Levy^63^), *Cen3* and *I3bc* (kindly provided by Gregory Copenhaver^64^), CTLs *3*.*9, 3*.*13, 2309-1, 3090, 3092, 2696, 2697, 2698* (kindly provided by Scott Poethig^67^). The Col mutants *fancm-1, zip4-2, recq4a-4, recq4b-2* and the L*er* mutants *recq4a* and *fancm-10* were kindly provided by Raphael Mercier^21,24,59,83^. All primer sequences used for genotyping of mutant lines are described in Supplementary Table 15.

### Genotyping-by-sequencing library preparation

DNA was extracted from leaves of F_2_ plants obtained from Col × L*er* cross. DNA extraction was performed as described^65^ and the quality of the DNA was checked in 1% agarose gel. Tagmentation was performed by mixing 1μL of 5ng/ μL of the DNA with 1 μL of Tagmentation Buffer (40mM Tris-HCl pH=7.5, 40 mM MgCl2), 0.5 μL of DMF (Sigma), 2.35 μL of Nuclease-free water (Thermo Fisher) and 0.05 μL of loaded, in-house produced Tn5. Loading Tn5 with the annealed linker oligonucleotides was previously described^84^. The tagmentation step was carried out at 55°C for 2 min and then stopped by adding 1 μL 0.1% SDS and incubating at 65°C for 10 min. Amplification of the tagmented DNA was performed using the KAPA2G Robust PCR kit (Sigma) and custom P5 and P7 indexing primers. Each sample was amplified with the unique set of P5 and P7 primers as described^42^. The successful libraries were pooled and size selected in 2% agarose gel, after which DNA fragments in a range of 400-700 bp were excised and extracted using Gel Extraction Kit (A&A Biotechnology) The quality and quantity of the libraries were verified with TapeStation system (Agilent) and Qubit 2.0 fluorometer. Paired-end sequencing of libraries was performed on HiSeq X-10 instrument (Illumina).

### Genotyping-by-sequencing bioinformatics analysis

To identify Single-nucleotide polymorphisms (SNPs) within the tested populations, demultiplexed paired-end forward and reverse reads have been pooled and aligned to Col-0 genome reference sequence with use of BowTie2^85^. Resulting BAM files have been sorted and indexed with use of SAMtools v1.2^86^. SNPs were called using SAMtools and BCFtools^87^. Subsequently, individual sequencing libraries have been aligned to Col-0 genome reference sequence with default parameters in BowTie2 and compared to previously generated SNP list with SAMtools and BCFtools. Later, the resulting tables of SNPs have been filtered to keep only SNPs with high mapping quality (>100) and high coverage (>2.5×) in R. Individual libraries with less than 100,000 reads were discarded from the analysis. To call crossovers TIGER pipeline has been used on filtered files^65^. Summary of GBS results is presented in Supplementary Table 16. To investigate CO distribution, crossover frequencies have been binned into scaled windows and summed across chromosome arms.

For analysis of the relationship between crossover recombination and SNP density, the genome was divided into 100 kb nonoverlapping windows and for each of them SNP density was determined based on published Col/L*er* polymorphism data^88^. The crossover frequency per each window was normalized to the number of analyzed individuals. This resulted in 1191 windows, which were sorted according to the SNP density and grouped into 99 groups, so that each group consisted of 12 windows with a similar polymorphism level (Supplementary Fig. 4 and Supplementary Table 6). Spearman rank correlation was used to assess relationship between SNPs and crossovers.

### Crossover frequency measurement using FTL seed-based system

Crossover rate measurements using seed-based system was performed as described previously^49,89^. Briefly, pictures of seeds were acquired using epifluorescent microscope in bright field, ultraviolet (UV) + dsRed filter, and UV + GFP filter. The images were later processed by CellProfiler software^90^ to identify seed boundaries and to assign a dsRed and eGFP fluorescence intensity value to each seed object. Thresholds between fluorescent and non-fluorescent seed were set manually using fluorescence histograms for each colour. The crossover frequency is calculated as cM = 100 × (1–(1−2(NG+NR)/NT)/2), where NG is the number of green alone seeds, NR is the number of red alone seeds, and NT is the total number of seeds.

### Crossover frequency and interference measurements using FTL pollen-based system

Samples for flow cytometry analysis were prepared as described previously^91^. Inflorescences from 5-8 individual plants were pooled for each experimental variant, with at least three biological replicates. The flow cytometry was performed on Guava easyCyte 8HT Cytometer (Millipore). The samples were analysed using GuavaSoft 3.3 programme (Millipore). Obtained events were separated based on forward and side scatter and hydrated pollen was gated to exclude dead or damaged material. For crossover frequency measurements in *CEN3* interval the events were gated into four classes based on their fluorescence emission signals: red (R), yellow (Y), double-colour (RY) and non-colour (N). Crossover frequency (cM) was calculated as 100 × (Y/(Y + RY)). For *I3bc* interval crossover interference measurements events were divided into eight classes. *I3b* and *I3c* genetic distances were calculated by dividing the sum of recombinant gametes in particular interval by the total number of pollen grains. Crossover interference was calculated by counting the coefficient of coincidence (CoC), which is the ratio between the expected and the observed double crossover (DCO) number. The expected DCO frequency is obtained by dividing by hundredths the genetic distances in *I3b* and *I3c* intervals and further multiplying them by the total number of pollen grains. The observed DCO is the sum of pollen grains that have experienced a double crossover (B-R and -Y-classes). Interference is then calculated as follows: 1− CoC.

### CRISPR/Cas9 mutagenesis of *ZIP4* in L*er*-0

CRISPR/Cas9 mutagenesis on Arabidopsis plants was performed according to the protocol^92,93^. To obtain *zip4* mutant line in L*er*-0 background, a pair of gRNAs targeted within exon 1 of *ZIP4* were designed. A vector containing the *ZIP4* gRNA pair under the U3 and U6 promoters, and a *ICU2::Cas9* transgene was used for *Agrobacterium* transformation. Transformants were genotyped by PCR amplification with primers flanking the *ZIP4* gRNA target sites. Sanger sequencing was performed to detect deletions - mutants with heritable deletions causing a frame shift in *ZIP4*, and not carrying the CRISPR-Cas9 construct, were identified for further experiments.

### Fertility assays

Seed set and silique length were assessed from five fruits, located at positions 6 through 10 of the main stem, in eight plants per genotype. Collected siliques were incubated in 70% EtOH for 72h and later photographed in the bright field using bottom light source, enabling seed set calculations, which were performed manually. Silique length was calculated using ImageJ software^94^.

### MSH2 protein overexpression

A DNA fragment containing *MSH2* genomic sequence including endogenous promoter was amplified from Col genomic DNA using msh2gf and msh2gr primers. The PCR product was cloned into the pFGC binary vector containing *bar* resistance gene using one step cloning kit (Vazyme). The vector was introduced into Col-*420* plants via *Agrobacterium* floral-dip transformation. T_1_ plants were treated with Basta for recombinant selection and later genotyped for construct presence. Three independent T_1_s were crossed with Col, Ct and I7-A13 plants. The resulting F_1_s were tested for the construct presence by PCR and used for *420* crossover frequency measurements using seed-based system.

### Chiasmata counting

Metaphase I chromosome spread preparations were prepared from ethanol: acetic acid (3:1) fixed material. Chiasma counts were based on the shape of bivalents. Significance was assessed by Kruskal-Wallis H test with Mann Whitney U test using Bonferroni correction.

## Data Availability

The GBS sequence data generated in this study have been deposited in the NCBI Sequence Read Archive (SRA) under the BioProject accession codes PRJNA952840. Seed scoring and pollen scoring raw data generated in this study are provided in the Supplementary Information. Source data are provided with this paper.

## Acknowledgements

We thank Prof. Raphael Mercier (Max Planck Institute for Plant Breeding Research, Cologne) for sharing *fancm zip4, recq4a recq4b* mutants in Col and L*er* backgrounds. The computations were performed at the Poznan Supercomputing and Networking Center (grant 312). This work was supported by the National Science Center, Poland (NCN) grants 2016/22/E/NZ2/00455 and 2020/39/I/NZ2/02464 to P.A.Z., 2021/41/N/NZ2/01226 to J.D., the Foundation for Polish Science grant (POIR.04.04.00-00-5C0F/17-00) to P.A.Z.

## Author Contributions Statement

J.D. and P.A.Z. designed research; J.D., M.Sz.-L., M.G. and J.H. performed research; I.R.H. contributed computational pipelines and materials; W.D. performed the computational analyses; J.D., W.D., M.Sz.-L., J.H. and P.A.Z. analyzed data; and J.D. and P.A.Z. wrote the paper with the aid of all authors.

