## Supplementary Figures and Tables for "MSH2 stimulates interfering and inhibits non-interfering crossovers in response to genetic polymorphism"

### Supplementary Information

### Supplementary Figures

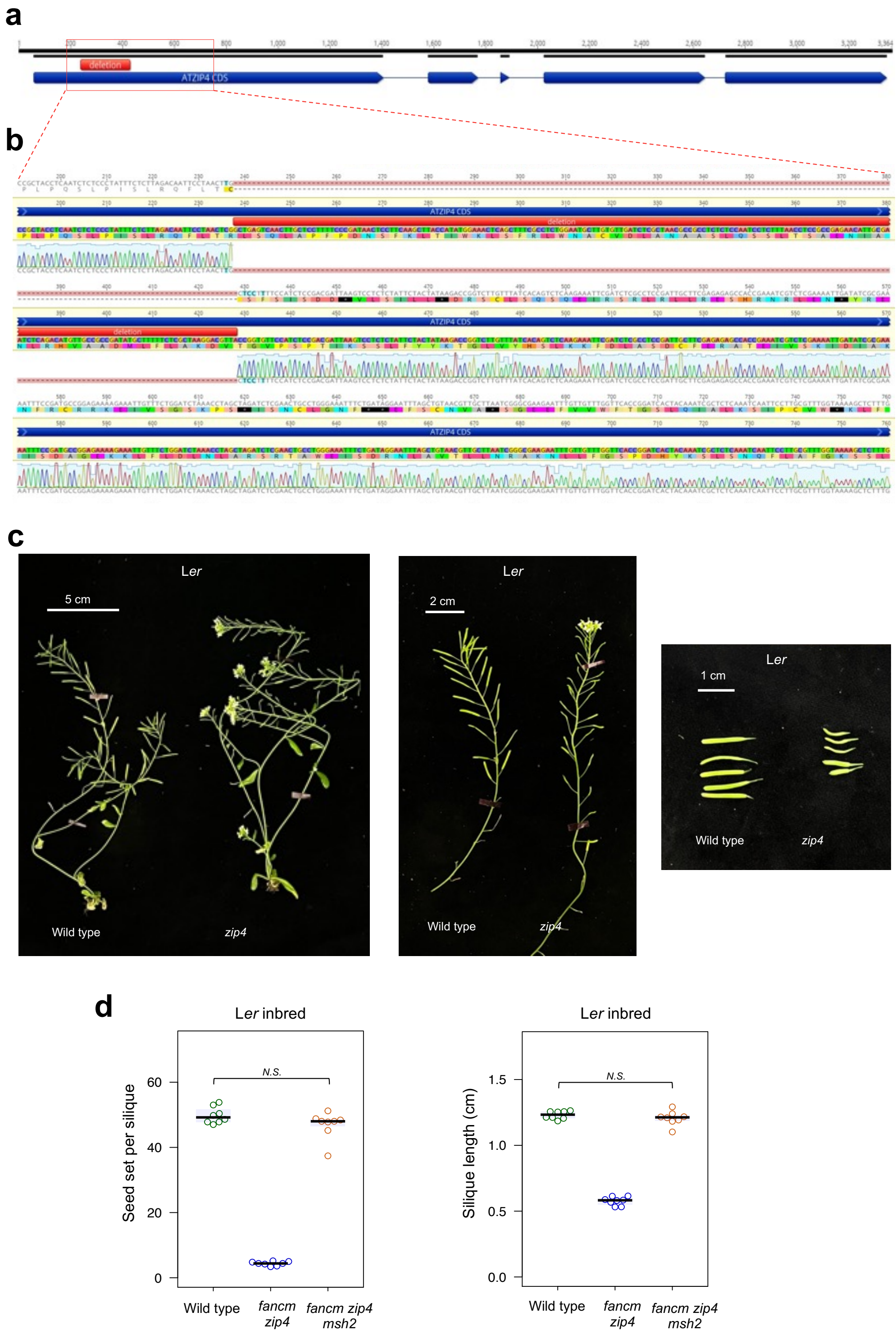

**Supplementary Fig. 1.** *ZIP4* CRISPR/Cas9-mediated mutagenesis in L*er* accession. **a**, ZIP4 coding sequence with exons represented in dark blue arrows. Mutagenesis by CRISPR/Cas9 resulted in 191 bp deletion within the first exon (annotated as a red square) and introduction of 5 SNPs within the Cas9 cutting sites. **b**, Close-up view of the L*er* *zip4* sequencing aligned to the wild-type reference shows that the 191 bp deletion results in a frameshift and multiple STOP codons, represented in black. **c**, The L*er* *zip4* mutant obtained by CRISPR/Cas9 shows dramatically reduced fertility. Whole plants (left) are shown next to primary shoots (center) and exemplary siliques (right). **d**, Fertility assays in L*er zip4* and *fancm zip4* as assessed via seed set and silique length. The center line of a boxplot indicates the mean; the upper and lower bounds indicate the 75th and 25th percentiles, respectively. Each dot represents a measurement from five siliques of one plant. Significance was assessed by Welch's *t*-test.

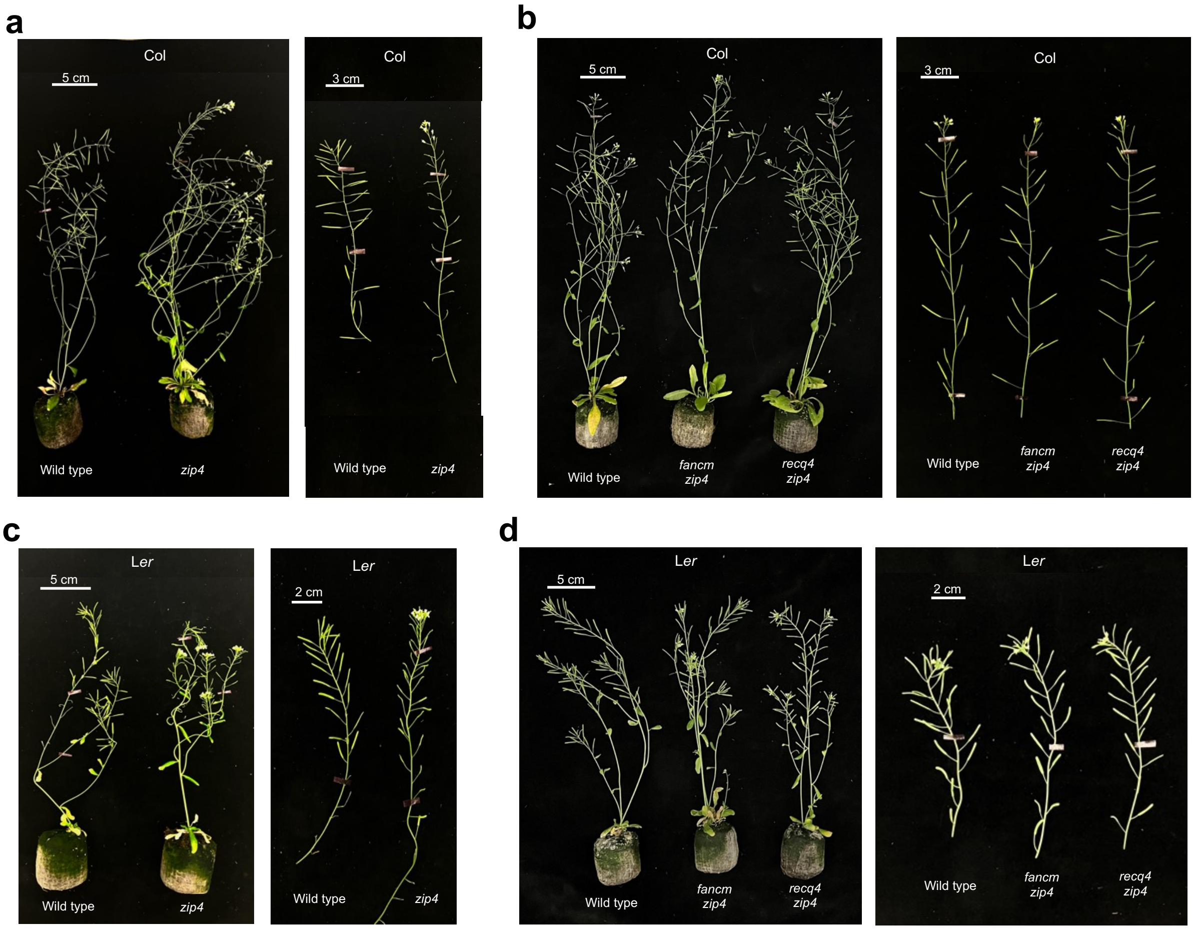

**Supplementary Fig. 2.** Fertility, which is lost in Col and L*er zip4* mutants, is restored in *fancm* or *recq4* (*recq4a recq4b*) mutant backgrounds. **a**, comparison of wild type and *zip4* plants (left) and shoots (right) in Col background. **b**, as in **a**, but for the *fancm zip4* and *recq4 zip4* mutants. **c**, as in **a**, but in the L*er* background. **d**, as in **a**, but for the *fancm zip4* and *recq4 zip4* mutants in L*er* background.

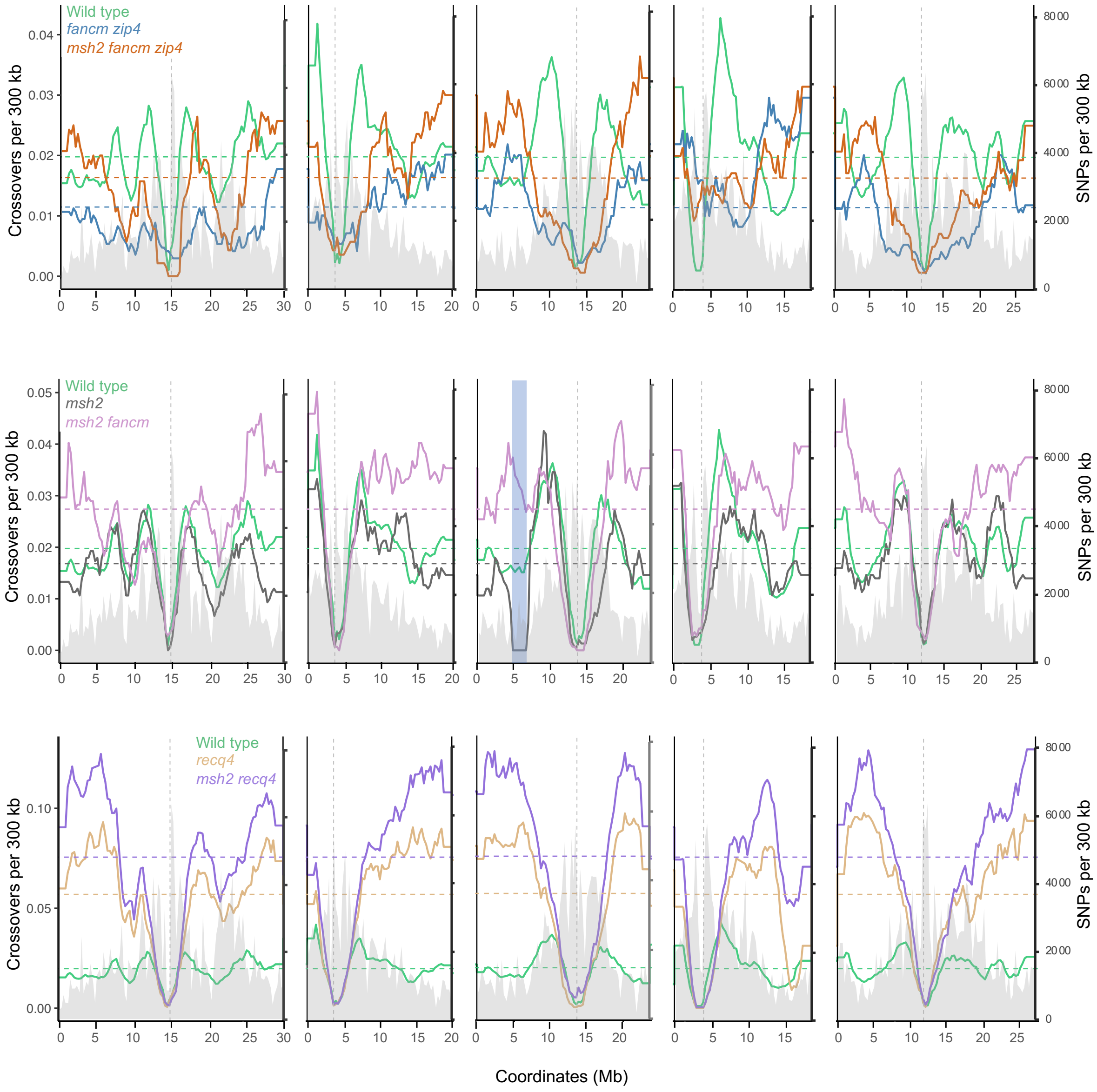

**Supplementary Fig. 3.** Crossovers per 300 kb per $F_2$ plotted along the Arabidopsis chromosomes. Mean values are shown by horizontal dashed lines. SNPs per 300 kb are plotted and shaded in gray. The position of centromeres is indicated as vertical dashed line. The introgression of Col in *msh2* L*er* due to *msh2* backcrossing results in failure to detect crossovers, which is indicated as a blue rectangle.

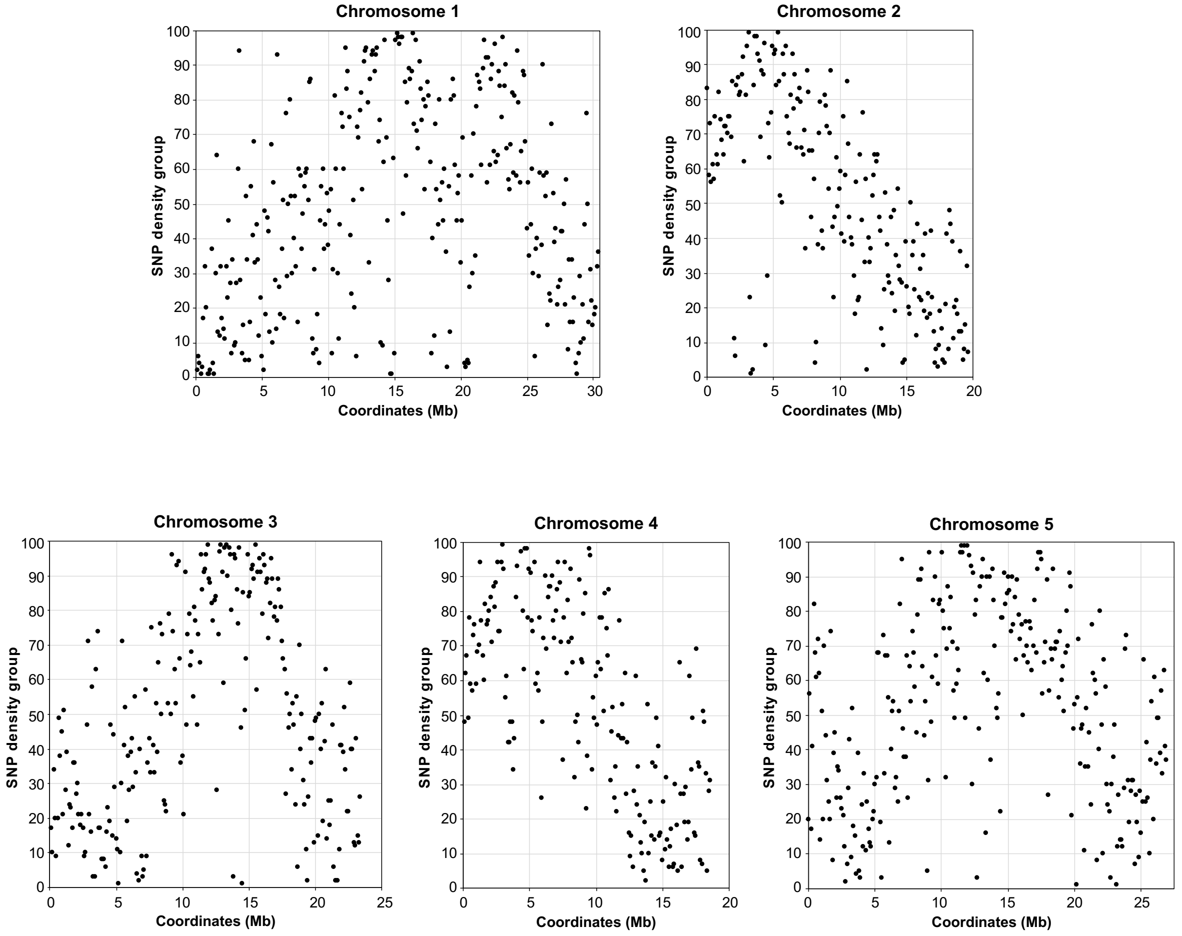

**Supplementary Fig. 4.** Chromosomal distribution of 100 kb windows for each SNP density group used in Figure 3. Each dot corresponds to one window, each group (Y-axis) consists of 12 windows.

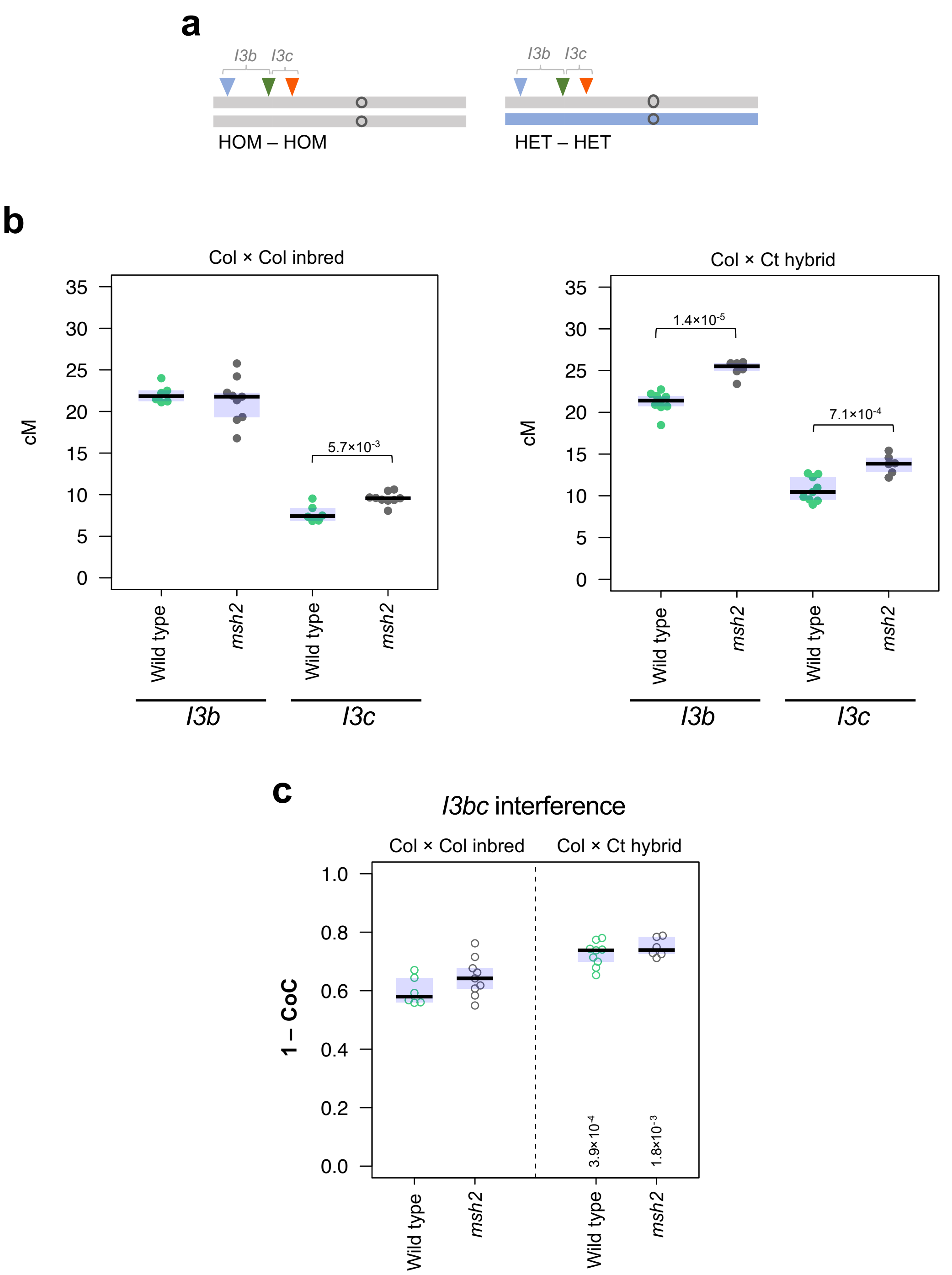

**Supplementary Fig. 5.** Crossover interference is not changed in *msh2* inbreds and hybrids. **a**, Ideograms of chromosome 3 showing the location of *I3b* and *I3c* intervals in Col/Col inbred and Col/Ct hybrid plants used in the experiment. Gray corresponds to Col while blue corresponds to Ct genotype. Location of fluorescent reporters defining *I3b* and *I3c* interval is indicated by blue, green and red triangles. **b**, *I3b* and *I3c* crossover rate as measured in Col/Col inbred and Col/Ct hybrid plants as measured in wild type and *msh2* plants. Welch's t-test was used to calculate statistical significance. **c**, Crossover interference (1 – CoC) as measured over *I3bc* intervals for Col/Col inbred and Col/Ct hybrid plants in wild type and *msh2* plants. Two-sided *P* values indicate the statistical difference in crossover interference in hybrids versus inbreds (Welch's t-test). Each dot on **b** and **c** represents a measurement from a pool of 5-8 individuals.

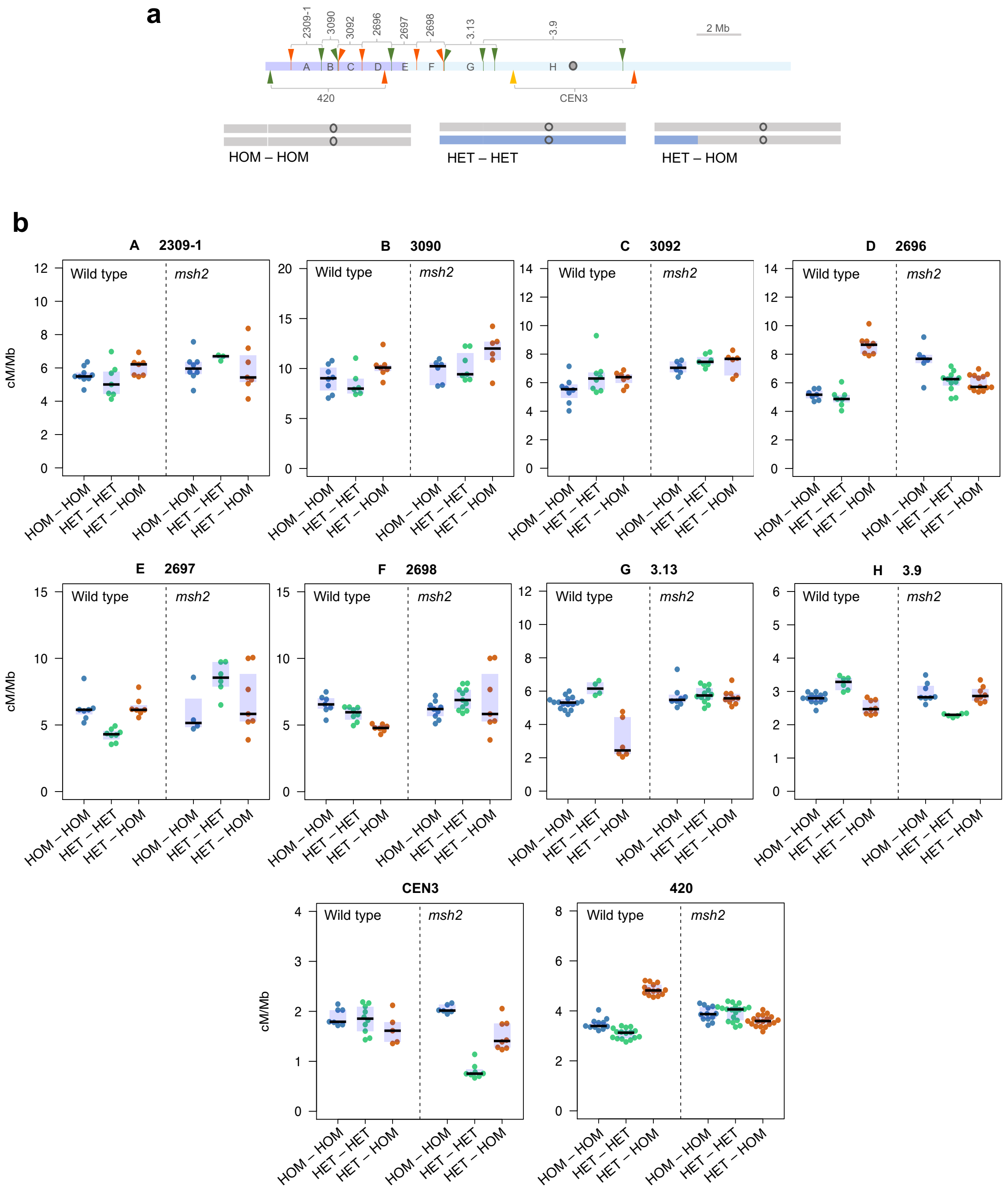

**Supplementary Fig. 6.** Crossover distribution across the chromosome arm in response to polymorphism. The plots show the same data as in Figure 5, but separately for each FTL. **a**, (upper panel) Location of eight FTL intervals across the Arabidopsis chromosome 3 shown in addition to *420* and *CEN3* intervals. Fluorescent reporters are indicated by tick marks and arrowheads with colors corresponding to eGFP (green), dsRed (red) and eYFP (yellow). The violet and light blue shading represents 'HET' and 'HOM' regions in HET-HOM line, respectively. Genetically defined centromere[1] is indicated as gray circle. (lower panel) Ideograms of chromosome 3 showing heterozygosity pattern in lines used in **b**. Gray corresponds to Col while blue corresponds to Ct genotype. **b**, Crossover frequency (cM/Mb) in the HOM-HOM, HET-HET and HET-HOM genotypes shown separately for ten FTLs, in either wild type or *msh2*. The center line of a boxplot indicates the mean; the upper and lower bounds indicate the 75th and 25th percentiles, respectively. Each dot represents a measurement from one individual.

**Supplementary Tables**

**Supplementary Table 1.** Seed set per silique in Col/L*er* hybrid in the wild type, *fancm zip4* and *msh2 fancm zip4* mutant lines. The number of seeds was counted manually in eight plants per genotype and five fruits per plant, located at positions 6 through 10 of the main stem. Mean values per individual are shown.

| | | **Seeds per silique in Col/L*er* hybrid, plant #** | | | | | | | |
|---|---|---|---|---|---|---|---|---|---|
| **Genotype** | **Silique** | **1** | **2** | **3** | **4** | **5** | **6** | **7** | **8** |
| **Wild type** | **1** | 58 | 25 | 56 | 65 | 60 | 60 | 57 | 18 |
| | **2** | 57 | 64 | 55 | 46 | 66 | 56 | 67 | 66 |
| | **3** | 54 | 61 | 70 | 55 | 59 | 61 | 67 | 68 |
| | **4** | 50 | 69 | 63 | 64 | 60 | 56 | 66 | 71 |
| | **5** | 65 | 29 | 37 | 66 | 62 | 9 | 61 | 70 |
| | **Mean** | 56.8 | 49.6 | 56.2 | 59.2 | 61.4 | 48.4 | 63.6 | 58.6 |
| ***fancm zip4*** | **1** | 9 | 6 | 8 | 11 | 1 | 9 | 9 | 8 |
| | **2** | 4 | 10 | 7 | 9 | 13 | 4 | 7 | 7 |
| | **3** | 12 | 13 | 6 | 9 | 10 | 8 | 4 | 4 |
| | **4** | 10 | 8 | 3 | 10 | 7 | 7 | 7 | 7 |
| | **5** | 4 | 8 | 4 | 3 | 14 | 6 | 6 | 6 |
| | **Mean** | 7.8 | 9 | 5.6 | 8.4 | 9 | 6.8 | 6.6 | 6.4 |
| ***msh2 fancm zip4*** | **1** | 28 | 10 | 13 | 21 | 22 | 23 | 26 | 15 |
| | **2** | 19 | 16 | 11 | 29 | 24 | 22 | 28 | 14 |
| | **3** | 16 | 20 | 22 | 22 | 21 | 20 | 25 | 9 |
| | **4** | 20 | 11 | 15 | 25 | 19 | 25 | 32 | 17 |
| | **5** | 20 | 24 | 18 | 21 | 23 | 20 | 22 | 15 |
| | **Mean** | 20.6 | 16.2 | 15.8 | 23.6 | 21.8 | 22 | 26.6 | 14 |

**Supplementary Table 2.** Silique length in Col/L*er* hybrid in the wild type, *fancm zip4* and *msh2 fancm zip4* mutant lines. Silique length was measured in eight plants per genotype and five fruits per plant, located at positions 6 through 10 of the main stem. Measurements were done with ImageJ software. Mean values per individual are shown.

| | | Silique length (cm) in Col/Ler hybrid, plant # | | | | | | | |
|---|---|---|---|---|---|---|---|---|---|
| **Genotype** | **Silique** | **1** | **2** | **3** | **4** | **5** | **6** | **7** | **8** |
| **Wild type** | **1** | 1.404 | 0.959 | 1.429 | 1.582 | 1.501 | 1.569 | 1.602 | 0.872 |
| | **2** | 1.383 | 1.71 | 1.473 | 1.173 | 1.703 | 1.545 | 1.618 | 1.712 |
| | **3** | 1.318 | 1.55 | 1.726 | 1.412 | 1.52 | 1.639 | 1.633 | 1.659 |
| | **4** | 1.237 | 1.798 | 1.531 | 1.55 | 1.474 | 1.47 | 1.553 | 1.803 |
| | **5** | 1.532 | 1.132 | 1.084 | 1.565 | 1.669 | 0.675 | 1.664 | 1.778 |
| | **Mean** | 1.3748 | 1.4298 | 1.4486 | 1.4564 | 1.5734 | 1.3796 | 1.614 | 1.5648 |
| ***fancm zip4*** | **1** | 0.831 | 0.704 | 0.783 | 0.932 | 0.492 | 0.835 | 0.845 | 0.819 |
| | **2** | 0.655 | 0.889 | 0.784 | 0.844 | 0.92 | 0.641 | 0.775 | 0.831 |
| | **3** | 0.92 | 0.943 | 0.822 | 0.876 | 0.891 | 0.79 | 0.659 | 0.701 |
| | **4** | 0.911 | 0.812 | 0.695 | 0.864 | 0.772 | 0.804 | 0.818 | 0.741 |
| | **5** | 0.723 | 0.837 | 0.644 | 0.6 | 0.935 | 0.729 | 0.783 | 0.72 |
| | **Mean** | 0.808 | 0.837 | 0.7456 | 0.8232 | 0.802 | 0.7598 | 0.776 | 0.7624 |
| ***msh2 fancm zip4*** | **1** | 1.342 | 0.877 | 1.019 | 1.138 | 1.223 | 1.289 | 1.315 | 1.04 |
| | **2** | 1.156 | 1.034 | 0.926 | 1.315 | 1.233 | 1.198 | 1.315 | 0.919 |
| | **3** | 1.035 | 1.161 | 1.102 | 1.129 | 1.165 | 1.187 | 1.297 | 0.852 |
| | **4** | 1.184 | 0.887 | 1.036 | 1.22 | 1.17 | 1.228 | 1.398 | 1.133 |
| | **5** | 1.212 | 1.226 | 1.049 | 1.168 | 1.309 | 1.118 | 1.204 | 0.996 |
| | **Mean** | 1.1858 | 1.037 | 1.0264 | 1.194 | 1.22 | 1.204 | 1.3058 | 0.988 |

**Supplementary Table 3.** Fertility analysis in wild type and *zip4* mutant in the L*er* background. The number of seeds per silique (seed set) was counted manually in eight plants per genotype and five fruits per plant, located at positions 6 through 10 of the main stem. Silique length was measured in eight plants per genotype and five fruits per plant, located at positions 6 through 10 of the main stem. Measurements were done with ImageJ software. Mean values per individual are shown.

| | **L*er* wt, plant #** | | | | | | | |
|---|---|---|---|---|---|---|---|---|
| **Silique length (cm)** | | | | | | | | |
| **Silique** | **1** | **2** | **3** | **4** | **5** | **6** | **7** | **8** |
| **1** | 1.25 | 1.24 | 1.23 | 1.20 | 1.24 | 1.20 | 1.22 | 1.17 |
| **2** | 1.26 | 1.30 | 1.17 | 1.19 | 1.26 | 1.19 | 1.11 | 1.24 |
| **3** | 1.27 | 1.27 | 1.23 | 1.19 | 1.30 | 1.21 | 1.19 | 1.25 |
| **4** | 1.21 | 1.25 | 1.24 | 1.20 | 1.28 | 1.25 | 1.23 | 1.29 |
| **5** | 1.29 | 1.23 | 1.19 | 1.14 | 1.25 | 1.22 | 1.28 | 1.33 |
| **Mean** | **1.25** | **1.26** | **1.21** | **1.19** | **1.26** | **1.21** | **1.21** | **1.26** |
| **Seeds per silique** | | | | | | | | |
| **1** | 47 | 47 | 51 | 51 | 51 | 44 | 48 | 49 |
| **2** | 53 | 50 | 57 | 48 | 55 | 46 | 46 | 43 |
| **3** | 54 | 45 | 58 | 50 | 58 | 52 | 51 | 50 |
| **4** | 44 | 47 | 52 | 46 | 50 | 49 | 51 | 48 |
| **5** | 51 | 50 | 47 | 48 | 55 | 44 | 56 | 49 |
| **Mean** | **49.8** | **47.8** | **53** | **48.6** | **53.8** | **47** | **50.4** | **47.8** |
| | **L*er zip4*, plant #** | | | | | | | |
| **Silique length (cm)** | | | | | | | | |
| **Silique** | **1** | **2** | **3** | **4** | **5** | **6** | **7** | **8** |
| **1** | 0.48 | 0.56 | 0.60 | 0.57 | 0.59 | 0.63 | 0.68 | 0.50 |
| **2** | 0.59 | 0.62 | 0.53 | 0.62 | 0.63 | 0.49 | 0.51 | 0.68 |
| **3** | 0.59 | 0.56 | 0.63 | 0.64 | 0.53 | 0.62 | 0.52 | 0.66 |
| **4** | 0.47 | 0.75 | 0.68 | 0.56 | 0.54 | 0.55 | 0.46 | 0.45 |
| **5** | 0.53 | 0.59 | 0.64 | 0.55 | 0.64 | 0.62 | 0.49 | 0.55 |
| **Mean** | **0.53** | **0.61** | **0.62** | **0.59** | **0.58** | **0.58** | **0.53** | **0.57** |
| **Seeds per silique** | | | | | | | | |
| **1** | 2 | 3 | 1 | 5 | 5 | 8 | 5 | 3 |
| **2** | 5 | 6 | 4 | 3 | 3 | 2 | 5 | 9 |
| **3** | 5 | 4 | 6 | 7 | 2 | 7 | 4 | 7 |
| **4** | 2 | 11 | 7 | 3 | 3 | 2 | 1 | 2 |
| **5** | 3 | 2 | 4 | 3 | 9 | 6 | 3 | 3 |
| **Mean** | **3.4** | **5.2** | **4.4** | **4.2** | **4.4** | **5** | **3.6** | **4.8** |

**Supplementary Table 4.** Chiasma frequency in Col/L*er* hybrid meiocytes in the wild type, *fancm zip4* and *msh2 fancm zip4* mutant lines. Total of 60 meiocytes were calculated for each genotype. Mean values for each genotype are shown.

| | **Genotype** | | |
|---|---|---|---|
| **Chiasmata per meiocyte** | **Wild type** | ***fancm zip4*** | ***msh2 fancm zip4*** |
| 0 | 0 | 16 | 1 |
| 1 | 0 | 19 | 2 |
| 2 | 0 | 19 | 8 |
| 3 | 0 | 3 | 18 |
| 4 | 0 | 2 | 15 |
| 5 | 0 | 1 | 7 |
| 6 | 0 | 0 | 8 |
| 7 | 2 | 0 | 1 |
| 8 | 8 | 0 | 0 |
| 9 | 27 | 0 | 0 |
| 10 | 20 | 0 | 0 |
| 11 | 3 | 0 | 0 |
| | Mean: 9.23 | Mean: 1.32 | Mean: 3.7 |

**Supplementary Table 5.** *420* fluorescent seed count data for the wild type, *msh2, fancm, msh2 fancm, fancm zip4* and *msh2 fancm zip4* mutants in Col/L*er* hybrid. Genetic distance in centimorgans (cM) is calculated as $cM = 100\times(1-(1-2(NG+NR)/NT)^{1/2})$, where NG is the number of green alone seeds, NR is the number of red alone seeds and NT is the total number of seeds of all classes analysed.

| Genotype | Total | Green alone | Red alone | Red+Green | Non-colour | cM |
|---|---|---|---|---|---|---|
| Wild type | 2110 | 156 | 152 | 1432 | 370 | 15.85 |
| Wild type | 1871 | 131 | 123 | 1304 | 313 | 14.65 |
| Wild type | 2243 | 130 | 145 | 1564 | 404 | 13.12 |
| Wild type | 2035 | 156 | 183 | 1364 | 332 | 18.34 |
| Wild type | 1770 | 152 | 192 | 1165 | 261 | 21.81 |
| Wild type | 2334 | 142 | 152 | 1639 | 401 | 13.51 |
| Wild type | 2140 | 143 | 143 | 1506 | 348 | 14.40 |
| Wild type | 2038 | 141 | 150 | 1405 | 342 | 15.48 |
| Wild type | 1849 | 159 | 133 | 1255 | 302 | 17.29 |
| Wild type | 1566 | 150 | 148 | 1024 | 244 | 21.30 |
| Wild type | 1723 | 134 | 149 | 1128 | 312 | 18.05 |
| Wild type | 2030 | 141 | 137 | 1381 | 371 | 14.79 |
| *msh2* | 2271 | 189 | 168 | 1491 | 423 | 17.20 |
| *msh2* | 2385 | 212 | 235 | 1561 | 377 | 20.93 |
| *msh2* | 2104 | 160 | 221 | 1361 | 362 | 20.14 |
| *msh2* | 2074 | 183 | 205 | 1351 | 335 | 20.89 |
| *msh2* | 1857 | 142 | 161 | 1221 | 333 | 17.92 |
| *msh2* | 2177 | 182 | 225 | 1424 | 346 | 20.87 |
| *msh2* | 2053 | 179 | 153 | 1386 | 335 | 17.75 |
| *msh2* | 1732 | 140 | 163 | 1168 | 261 | 19.37 |
| *msh2* | 1872 | 205 | 151 | 1218 | 298 | 21.28 |
| *fancm* | 2538 | 160 | 198 | 1743 | 437 | 15.27 |
| *fancm* | 2417 | 192 | 182 | 1626 | 417 | 16.90 |
| *fancm* | 1995 | 145 | 151 | 1345 | 354 | 16.14 |
| *fancm* | 2599 | 149 | 186 | 1876 | 388 | 13.85 |
| *fancm* | 2426 | 138 | 167 | 1695 | 426 | 13.48 |
| *fancm* | 2261 | 163 | 163 | 1560 | 375 | 15.64 |
| *fancm* | 996 | 81 | 82 | 660 | 173 | 17.98 |
| *fancm* | 2222 | 153 | 177 | 1502 | 390 | 16.16 |
| *fancm* | 1944 | 150 | 135 | 1297 | 362 | 15.93 |
| *fancm* | 2196 | 164 | 161 | 1505 | 366 | 16.09 |
| *msh2 fancm* | 1485 | 190 | 144 | 947 | 204 | 25.83 |
| *msh2 fancm* | 1148 | 101 | 107 | 721 | 219 | 20.15 |
| *msh2 fancm* | 1825 | 193 | 185 | 1148 | 299 | 23.47 |
| *msh2 fancm* | 1528 | 165 | 160 | 978 | 225 | 24.20 |
| *msh2 fancm* | 2081 | 236 | 204 | 1332 | 309 | 24.03 |
| *msh2 fancm* | 1479 | 150 | 122 | 946 | 261 | 20.49 |
| *msh2 fancm* | 1705 | 144 | 151 | 1129 | 281 | 19.13 |
| *msh2 fancm* | 1375 | 137 | 132 | 884 | 222 | 21.98 |

| | | | | | | |
|---|---|---|---|---|---|---|
| *msh2 fancm* | 1984 | 202 | 182 | 1247 | 353 | 21.71 |
| *fancm zip4* | 1337 | 82 | 73 | 934 | 248 | 12.36 |
| *fancm zip4* | 1247 | 76 | 86 | 840 | 245 | 13.97 |
| *fancm zip4* | 1219 | 84 | 72 | 830 | 233 | 13.74 |
| *fancm zip4* | 1133 | 75 | 66 | 768 | 224 | 13.33 |
| *fancm zip4* | 1026 | 66 | 64 | 707 | 189 | 13.59 |
| *fancm zip4* | 1221 | 84 | 74 | 858 | 205 | 13.91 |
| *msh2 fancm zip4* | 868 | 93 | 85 | 524 | 166 | 23.20 |
| *msh2 fancm zip4* | 1134 | 111 | 113 | 746 | 164 | 22.22 |
| *msh2 fancm zip4* | 1027 | 125 | 97 | 641 | 164 | 24.66 |
| *msh2 fancm zip4* | 987 | 95 | 115 | 628 | 149 | 24.21 |
| *msh2 fancm zip4* | 1334 | 164 | 124 | 847 | 199 | 24.62 |
| *msh2 fancm zip4* | 759 | 78 | 78 | 480 | 123 | 23.26 |
| *msh2 fancm zip4* | 1421 | 139 | 130 | 924 | 228 | 21.17 |
| *msh2 fancm zip4* | 1482 | 151 | 131 | 955 | 245 | 21.30 |

**Supplementary Table 6.** SNP density ranges in 99 groups used for SNP/crossover correlation study. Each group consists of 12 windows of 100 kb.

| Group | SNP/kb | | Group | SNP/kb | |
|---|---|---|---|---|---|
| | From | To | | From | To |
| 1 | 0.0 | 4.7 | 51 | 543.7 | 553.7 |
| 2 | 4.7 | 36.0 | 52 | 553.7 | 568.2 |
| 3 | 36.0 | 70.6 | 53 | 568.2 | 576.5 |
| 4 | 70.6 | 112.6 | 54 | 576.5 | 588.9 |
| 5 | 112.6 | 127.8 | 55 | 588.9 | 604.0 |
| 6 | 127.8 | 142.3 | 56 | 604.0 | 618.0 |
| 7 | 142.3 | 160.6 | 57 | 618.0 | 630.3 |
| 8 | 160.6 | 176.9 | 58 | 630.3 | 640.3 |
| 9 | 176.9 | 190.0 | 59 | 640.3 | 647.4 |
| 10 | 190.0 | 202.9 | 60 | 647.4 | 668.0 |
| 11 | 202.9 | 212.0 | 61 | 668.0 | 682.8 |
| 12 | 212.0 | 221.4 | 62 | 682.8 | 698.3 |
| 13 | 221.4 | 234.8 | 63 | 698.3 | 714.4 |
| 14 | 234.8 | 245.5 | 64 | 714.4 | 724.9 |
| 15 | 245.5 | 253.6 | 65 | 724.9 | 741.5 |
| 16 | 253.6 | 262.4 | 66 | 741.5 | 758.1 |
| 17 | 262.4 | 270.1 | 67 | 758.1 | 771.2 |
| 18 | 270.1 | 276.3 | 68 | 771.2 | 788.6 |
| 19 | 276.3 | 285.9 | 69 | 788.6 | 804.1 |
| 20 | 285.9 | 294.4 | 70 | 804.1 | 827.8 |
| 21 | 294.4 | 300.8 | 71 | 827.8 | 846.0 |
| 22 | 300.8 | 309.8 | 72 | 846.0 | 867.3 |
| 23 | 309.8 | 316.4 | 73 | 867.3 | 882.6 |
| 24 | 316.4 | 322.3 | 74 | 882.6 | 903.4 |
| 25 | 322.3 | 328.4 | 75 | 903.4 | 919.3 |
| 26 | 328.4 | 337.6 | 76 | 919.3 | 930.0 |
| 27 | 337.6 | 345.3 | 77 | 930.0 | 947.6 |
| 28 | 345.3 | 354.0 | 78 | 947.6 | 962.3 |
| 29 | 354.0 | 365.7 | 79 | 962.3 | 972.8 |
| 30 | 365.7 | 374.2 | 80 | 972.8 | 990.8 |
| 31 | 374.2 | 383.7 | 81 | 990.8 | 1019.6 |
| 32 | 383.7 | 394.3 | 82 | 1019.6 | 1035.1 |
| 33 | 394.3 | 399.2 | 83 | 1035.1 | 1051.3 |
| 34 | 399.2 | 404.3 | 84 | 1051.3 | 1079.3 |
| 35 | 404.3 | 411.0 | 85 | 1079.3 | 1108.4 |
| 36 | 411.0 | 419.3 | 86 | 1108.4 | 1144.6 |
| 37 | 419.3 | 427.8 | 87 | 1144.6 | 1170.3 |
| 38 | 427.8 | 436.7 | 88 | 1170.3 | 1209.5 |

| | | | | | |
|---|---|---|---|---|---|
| 39 | 436.7 | 447.4 | 89 | 1209.5 | 1239.6 |
| 40 | 447.4 | 453.3 | 90 | 1239.6 | 1261.1 |
| 41 | 453.3 | 461.8 | 91 | 1261.1 | 1287.3 |
| 42 | 461.8 | 466.9 | 92 | 1287.3 | 1341.3 |
| 43 | 466.9 | 472.3 | 93 | 1341.3 | 1400.2 |
| 44 | 472.3 | 482.2 | 94 | 1400.2 | 1469.3 |
| 45 | 482.2 | 498.0 | 95 | 1469.3 | 1564.9 |
| 46 | 498.0 | 509.8 | 96 | 1564.9 | 1670.3 |
| 47 | 509.8 | 517.0 | 97 | 1670.3 | 1821.6 |
| 48 | 517.0 | 524.3 | 98 | 1821.6 | 2062.8 |
| 49 | 524.3 | 533.3 | 99 | 2062.8 | 2404.5 |
| 50 | 533.3 | 543.7 | | | |

86

**Supplementary Table 7.** *420* fluorescent seed count data for the wild type and *msh2* mutants in recombinant lines with differing pattern of heterozygosity. The following recombinant backgrounds were used: 'HOM-HOM' that are Col/Col inbred throughout the genome, 'HET-HET' that are Col/Ct heterozygous throughout the genome, 'HET-HOM' where the *420* region is Col/Ct heterozygous and the remainder of chromosome 3 is Col/Col homozygous, and 'HOM-HET' where *420* is Col/Col homozygous and the remainder of chr3 is Col/Ct heterozygous. Genetic distance in centimorgans (cM) is calculated as cM = $100\times(1-(1-2(NG+NR)/NT)^{1/2})$, where NG is the number of green alone seeds, NR is the number of red alone seeds and NT is the total number of seeds of all classes analysed.

| Cross | Genotype | Total | Green alone | Red alone | Red+ Green | Non-colour | cM |
|---|---|---|---|---|---|---|---|
| HOM-HOM | Wild type | 1997 | 164 | 176 | 1301 | 356 | 18.79 |
| HOM-HOM | Wild type | 1469 | 122 | 108 | 968 | 271 | 17.12 |
| HOM-HOM | Wild type | 2031 | 165 | 155 | 1344 | 367 | 17.24 |
| HOM-HOM | Wild type | 2167 | 186 | 153 | 1448 | 380 | 17.11 |
| HOM-HOM | Wild type | 2287 | 224 | 199 | 1470 | 394 | 20.62 |
| HOM-HOM | Wild type | 2226 | 188 | 177 | 1451 | 410 | 18.02 |
| HOM-HOM | Wild type | 2012 | 152 | 151 | 1332 | 377 | 16.41 |
| HOM-HOM | Wild type | 1743 | 150 | 126 | 1147 | 320 | 17.34 |
| HOM-HOM | Wild type | 1802 | 131 | 145 | 1202 | 324 | 16.71 |
| HOM-HOM | Wild type | 1929 | 171 | 149 | 1269 | 340 | 18.26 |
| HOM-HOM | Wild type | 1593 | 149 | 112 | 1058 | 274 | 18.01 |
| HET-HET | Wild type | 2056 | 146 | 135 | 1390 | 385 | 14.76 |
| HET-HET | Wild type | 2127 | 155 | 164 | 1400 | 408 | 16.33 |
| HET-HET | Wild type | 2116 | 147 | 149 | 1406 | 414 | 15.13 |
| HET-HET | Wild type | 1847 | 115 | 159 | 1248 | 325 | 16.14 |
| HET-HET | Wild type | 1850 | 151 | 135 | 1225 | 339 | 16.88 |
| HET-HET | Wild type | 1674 | 113 | 134 | 1126 | 301 | 16.04 |
| HET-HET | Wild type | 2096 | 140 | 145 | 1454 | 357 | 14.67 |
| HET-HET | Wild type | 1954 | 133 | 137 | 1296 | 388 | 14.93 |
| HET-HET | Wild type | 2140 | 161 | 136 | 1445 | 398 | 15.00 |
| HET-HET | Wild type | 2387 | 158 | 162 | 1589 | 478 | 14.45 |
| HET-HET | Wild type | 2232 | 167 | 184 | 1463 | 418 | 17.21 |
| HET-HET | Wild type | 1864 | 138 | 106 | 1263 | 357 | 14.08 |
| HET-HET | Wild type | 1340 | 101 | 108 | 920 | 211 | 17.05 |
| HET-HET | Wild type | 2024 | 140 | 156 | 1337 | 391 | 15.89 |
| HET-HET | Wild type | 1884 | 145 | 154 | 1260 | 325 | 17.38 |
| HET-HET | Wild type | 2174 | 179 | 143 | 1450 | 402 | 16.11 |
| HET-HOM | Wild type | 2001 | 231 | 193 | 1267 | 310 | 24.09 |
| HET-HOM | Wild type | 1807 | 186 | 205 | 1153 | 263 | 24.68 |
| HET-HOM | Wild type | 1693 | 178 | 208 | 1063 | 244 | 26.24 |
| HET-HOM | Wild type | 1782 | 176 | 198 | 1127 | 281 | 23.83 |

| | | | | | | | |
|---|---|---|---|---|---|---|---|
| HET-HOM | Wild type | 1891 | 210 | 199 | 1211 | 271 | 24.67 |
| HET-HOM | Wild type | 2152 | 231 | 231 | 1349 | 341 | 24.46 |
| HET-HOM | Wild type | 1907 | 238 | 200 | 1198 | 271 | 26.47 |
| HET-HOM | Wild type | 2013 | 231 | 185 | 1302 | 295 | 23.40 |
| HET-HOM | Wild type | 2016 | 220 | 194 | 1286 | 316 | 23.24 |
| HET-HOM | Wild type | 1409 | 150 | 175 | 906 | 178 | 26.61 |
| HET-HOM | Wild type | 2291 | 251 | 242 | 1475 | 323 | 24.53 |
| HET-HOM | Wild type | 1355 | 141 | 141 | 896 | 177 | 23.60 |
| HET-HOM | Wild type | 1823 | 195 | 214 | 1185 | 229 | 25.75 |
| HET-HOM | Wild type | 1520 | 173 | 162 | 993 | 192 | 25.22 |
| HOM-HET | Wild type | 2049 | 101 | 104 | 1417 | 427 | 10.56 |
| HOM-HET | Wild type | 2027 | 97 | 114 | 1419 | 397 | 11.02 |
| HOM-HET | Wild type | 1842 | 123 | 81 | 1269 | 369 | 11.77 |
| HOM-HET | Wild type | 1908 | 92 | 100 | 1318 | 398 | 10.63 |
| HOM-HET | Wild type | 2069 | 82 | 123 | 1437 | 427 | 10.45 |
| HOM-HET | Wild type | 1365 | 65 | 57 | 970 | 273 | 9.38 |
| HOM-HET | Wild type | 2186 | 99 | 107 | 1529 | 451 | 9.92 |
| HOM-HET | Wild type | 2051 | 83 | 114 | 1423 | 431 | 10.12 |
| HOM-HET | Wild type | 1935 | 105 | 100 | 1321 | 409 | 11.22 |
| HOM-HET | Wild type | 1605 | 69 | 62 | 1149 | 325 | 8.53 |
| HOM-HET | Wild type | 1436 | 64 | 89 | 999 | 284 | 11.29 |
| HOM-HET | Wild type | 2182 | 100 | 123 | 1511 | 448 | 10.80 |
| HOM-HET | Wild type | 1042 | 52 | 55 | 738 | 197 | 10.86 |
| HOM-HOM | *msh2* | 1187 | 102 | 100 | 796 | 189 | 18.78 |
| HOM-HOM | *msh2* | 1946 | 162 | 186 | 1288 | 310 | 19.85 |
| HOM-HOM | *msh2* | 1946 | 165 | 166 | 1258 | 357 | 18.77 |
| HOM-HOM | *msh2* | 1618 | 152 | 137 | 1042 | 287 | 19.83 |
| HOM-HOM | *msh2* | 1775 | 164 | 170 | 1155 | 286 | 21.03 |
| HOM-HOM | *msh2* | 2142 | 165 | 177 | 1460 | 340 | 17.50 |
| HOM-HOM | *msh2* | 2014 | 194 | 200 | 1322 | 298 | 21.98 |
| HOM-HOM | *msh2* | 1818 | 155 | 142 | 1198 | 323 | 17.95 |
| HOM-HOM | *msh2* | 1808 | 149 | 172 | 1190 | 297 | 19.69 |
| HOM-HOM | *msh2* | 2305 | 191 | 208 | 1547 | 359 | 19.14 |
| HOM-HOM | *msh2* | 2055 | 220 | 177 | 1303 | 355 | 21.67 |
| HOM-HOM | *msh2* | 1442 | 166 | 109 | 929 | 238 | 21.35 |
| HET-HET | *msh2* | 968 | 89 | 86 | 648 | 145 | 20.10 |
| HET-HET | *msh2* | 1364 | 137 | 116 | 888 | 223 | 20.69 |
| HET-HET | *msh2* | 1447 | 140 | 136 | 960 | 211 | 21.35 |
| HET-HET | *msh2* | 809 | 68 | 72 | 532 | 137 | 19.14 |
| HET-HET | *msh2* | 1097 | 112 | 103 | 696 | 186 | 22.02 |
| HET-HET | *msh2* | 1291 | 105 | 100 | 877 | 209 | 17.39 |

| | | | | | | | |
|---|---|---|---|---|---|---|---|
| HET-HET | *msh2* | 1265 | 89 | 109 | 854 | 213 | 17.12 |
| HET-HET | *msh2* | 1646 | 130 | 175 | 1080 | 261 | 20.66 |
| HET-HET | *msh2* | 1646 | 136 | 134 | 1123 | 253 | 18.03 |
| HET-HET | *msh2* | 1761 | 143 | 149 | 1195 | 274 | 18.25 |
| HET-HET | *msh2* | 1467 | 133 | 113 | 981 | 240 | 18.48 |
| HET-HET | *msh2* | 1658 | 154 | 157 | 1065 | 282 | 20.95 |
| HET-HET | *msh2* | 1041 | 107 | 100 | 660 | 174 | 22.39 |
| HET-HET | *msh2* | 1305 | 128 | 115 | 859 | 203 | 20.78 |
| HET-HET | *msh2* | 911 | 86 | 86 | 610 | 129 | 21.11 |
| HET-HET | *msh2* | 954 | 85 | 96 | 628 | 145 | 21.23 |
| HET-HET | *msh2* | 932 | 91 | 93 | 590 | 158 | 22.21 |
| HET-HET | *msh2* | 1376 | 125 | 131 | 916 | 204 | 20.76 |
| HET-HOM | *msh2* | 2004 | 184 | 150 | 1316 | 354 | 18.35 |
| HET-HOM | *msh2* | 2223 | 198 | 194 | 1475 | 356 | 19.54 |
| HET-HOM | *msh2* | 2339 | 154 | 194 | 1608 | 383 | 16.19 |
| HET-HOM | *msh2* | 2281 | 200 | 180 | 1489 | 412 | 18.34 |
| HET-HOM | *msh2* | 2053 | 163 | 158 | 1413 | 319 | 17.10 |
| HET-HOM | *msh2* | 2245 | 177 | 176 | 1506 | 386 | 17.20 |
| HET-HOM | *msh2* | 2187 | 173 | 181 | 1447 | 386 | 17.76 |
| HET-HOM | *msh2* | 2053 | 197 | 159 | 1326 | 371 | 19.18 |
| HET-HOM | *msh2* | 2040 | 200 | 160 | 1366 | 314 | 19.56 |
| HET-HOM | *msh2* | 1771 | 162 | 166 | 1141 | 302 | 20.65 |
| HET-HOM | *msh2* | 2293 | 195 | 195 | 1549 | 354 | 18.77 |
| HET-HOM | *msh2* | 2297 | 175 | 216 | 1545 | 361 | 18.79 |
| HET-HOM | *msh2* | 2096 | 172 | 166 | 1414 | 344 | 17.69 |
| HET-HOM | *msh2* | 2365 | 199 | 174 | 1578 | 414 | 17.26 |
| HET-HOM | *msh2* | 2343 | 181 | 208 | 1534 | 420 | 18.27 |
| HET-HOM | *msh2* | 2082 | 160 | 190 | 1399 | 333 | 18.53 |
| HET-HOM | *msh2* | 2022 | 179 | 152 | 1362 | 329 | 17.99 |
| HET-HOM | *msh2* | 2013 | 174 | 187 | 1297 | 355 | 19.92 |
| HOM-HET | *msh2* | 1411 | 164 | 148 | 897 | 202 | 25.32 |
| HOM-HET | *msh2* | 2193 | 218 | 233 | 1386 | 356 | 23.27 |
| HOM-HET | *msh2* | 1454 | 142 | 156 | 946 | 210 | 23.18 |
| HOM-HET | *msh2* | 2074 | 213 | 196 | 1350 | 315 | 22.18 |
| HOM-HET | *msh2* | 1701 | 170 | 170 | 1121 | 240 | 22.53 |
| HOM-HET | *msh2* | 1912 | 220 | 234 | 1223 | 235 | 27.54 |
| HOM-HET | *msh2* | 1673 | 191 | 169 | 1063 | 250 | 24.53 |
| HOM-HET | *msh2* | 2279 | 200 | 246 | 1477 | 356 | 21.99 |
| HOM-HET | *msh2* | 1677 | 175 | 173 | 1085 | 244 | 23.52 |
| HOM-HET | *msh2* | 1651 | 155 | 178 | 1083 | 235 | 22.76 |
| HOM-HET | *msh2* | 2210 | 248 | 227 | 1432 | 303 | 24.49 |

| | | | | | | | |
|---|---|---|---|---|---|---|---|
| HOM-HET | *msh2* | 1704 | 171 | 207 | 1074 | 252 | 25.41 |
| HOM-HET | *msh2* | 1860 | 192 | 192 | 1213 | 263 | 23.38 |
| HOM-HET | *msh2* | 1875 | 209 | 192 | 1171 | 303 | 24.35 |
| HOM-HET | *msh2* | 1092 | 120 | 98 | 699 | 175 | 22.49 |
| HOM-HET | *msh2* | 947 | 94 | 84 | 604 | 165 | 21.00 |
| HOM-HET | *msh2* | 1885 | 149 | 167 | 1222 | 347 | 18.47 |
| HOM-HET | *msh2* | 2051 | 165 | 195 | 1334 | 357 | 19.44 |
| HOM-HET | *msh2* | 1327 | 127 | 142 | 857 | 201 | 22.89 |
| HOM-HET | *msh2* | 1405 | 142 | 130 | 897 | 236 | 21.72 |
| HOM-HET | *msh2* | 1064 | 96 | 115 | 678 | 175 | 22.32 |
| HOM-HOM | *fancm* | 2055 | 279 | 303 | 1270 | 203 | 34.15 |
| HOM-HOM | *fancm* | 2322 | 293 | 330 | 1479 | 220 | 31.93 |
| HOM-HOM | *fancm* | 1939 | 272 | 275 | 1205 | 187 | 33.99 |
| HOM-HOM | *fancm* | 2207 | 274 | 349 | 1339 | 245 | 34.01 |
| HOM-HOM | *fancm* | 2135 | 245 | 294 | 1391 | 205 | 29.64 |
| HOM-HOM | *fancm* | 2324 | 291 | 306 | 1475 | 252 | 30.27 |
| HOM-HOM | *fancm* | 2375 | 323 | 328 | 1434 | 290 | 32.78 |
| HOM-HOM | *fancm* | 1974 | 247 | 321 | 1190 | 216 | 34.84 |
| HOM-HOM | *fancm* | 982 | 126 | 155 | 617 | 84 | 34.60 |
| HET-HET | *fancm* | 1358 | 129 | 167 | 861 | 201 | 24.90 |
| HET-HET | *fancm* | 2203 | 195 | 222 | 1430 | 356 | 21.17 |
| HET-HET | *fancm* | 2298 | 234 | 230 | 1502 | 332 | 22.79 |
| HET-HET | *fancm* | 1972 | 171 | 199 | 1292 | 310 | 20.96 |
| HET-HET | *fancm* | 2118 | 189 | 212 | 1369 | 348 | 21.17 |
| HET-HET | *fancm* | 1937 | 193 | 163 | 1256 | 325 | 20.48 |
| HET-HET | *fancm* | 2175 | 210 | 214 | 1398 | 353 | 21.89 |
| HET-HET | *fancm* | 1903 | 185 | 181 | 1249 | 288 | 21.56 |
| HET-HET | *fancm* | 2162 | 224 | 216 | 1366 | 356 | 23.00 |
| HET-HET | *fancm* | 1197 | 111 | 108 | 807 | 171 | 20.37 |
| HET-HET | *fancm* | 1678 | 149 | 182 | 1072 | 275 | 22.19 |
| HET-HOM | *fancm* | 1756 | 185 | 239 | 1103 | 229 | 28.09 |
| HET-HOM | *fancm* | 1987 | 211 | 258 | 1241 | 277 | 27.34 |
| HET-HOM | *fancm* | 2056 | 251 | 312 | 1252 | 241 | 32.74 |
| HET-HOM | *fancm* | 1306 | 164 | 188 | 841 | 113 | 32.11 |
| HET-HOM | *fancm* | 1485 | 199 | 176 | 929 | 181 | 29.65 |
| HET-HOM | *fancm* | 909 | 96 | 135 | 549 | 129 | 29.88 |
| HET-HOM | *fancm* | 1134 | 130 | 157 | 719 | 128 | 29.73 |
| HET-HOM | *fancm* | 1882 | 211 | 289 | 1218 | 164 | 31.54 |
| HET-HOM | *fancm* | 1484 | 187 | 222 | 879 | 196 | 33.01 |
| HOM-HET | *fancm* | 1366 | 201 | 193 | 804 | 168 | 34.95 |
| HOM-HET | *fancm* | 2149 | 283 | 293 | 1306 | 267 | 31.89 |

| | | | | | | | |
|---|---|---|---|---|---|---|---|
| HOM-HET | *fancm* | 2314 | 267 | 345 | 1430 | 272 | 31.37 |
| HOM-HET | *fancm* | 2215 | 301 | 345 | 1334 | 235 | 35.45 |
| HOM-HET | *fancm* | 2199 | 313 | 316 | 1320 | 250 | 34.58 |
| HOM-HET | *fancm* | 1996 | 272 | 302 | 1184 | 238 | 34.82 |
| HOM-HET | *fancm* | 2096 | 257 | 277 | 1311 | 251 | 29.97 |
| HOM-HET | *fancm* | 1477 | 184 | 224 | 909 | 160 | 33.10 |
| HOM-HET | *fancm* | 1539 | 206 | 201 | 912 | 220 | 31.36 |
| HOM-HOM | *msh2 fancm* | 1774 | 300 | 230 | 1013 | 231 | 36.56 |
| HOM-HOM | *msh2 fancm* | 825 | 126 | 140 | 466 | 93 | 40.41 |
| HOM-HOM | *msh2 fancm* | 1055 | 162 | 178 | 616 | 99 | 40.38 |
| HOM-HOM | *msh2 fancm* | 2025 | 305 | 278 | 1200 | 242 | 34.87 |
| HOM-HOM | *msh2 fancm* | 1741 | 237 | 223 | 1080 | 201 | 31.33 |
| HOM-HOM | *msh2 fancm* | 2017 | 308 | 332 | 1198 | 179 | 39.55 |
| HOM-HOM | *msh2 fancm* | 1772 | 252 | 263 | 1051 | 206 | 35.29 |
| HOM-HOM | *msh2 fancm* | 1464 | 177 | 224 | 944 | 119 | 32.76 |
| HOM-HOM | *msh2 fancm* | 1790 | 293 | 283 | 1041 | 173 | 40.30 |
| HET-HET | *msh2 fancm* | 1424 | 131 | 216 | 923 | 154 | 28.40 |
| HET-HET | *msh2 fancm* | 1896 | 198 | 211 | 1211 | 276 | 24.60 |
| HET-HET | *msh2 fancm* | 367 | 49 | 45 | 229 | 44 | 30.16 |
| HET-HET | *msh2 fancm* | 2224 | 287 | 270 | 1370 | 297 | 29.35 |
| HET-HET | *msh2 fancm* | 1846 | 247 | 231 | 1123 | 245 | 30.56 |
| HET-HET | *msh2 fancm* | 1923 | 225 | 293 | 1144 | 261 | 32.08 |
| HET-HET | *msh2 fancm* | 1747 | 260 | 231 | 1049 | 207 | 33.83 |
| HET-HET | *msh2 fancm* | 1646 | 239 | 223 | 968 | 216 | 33.77 |
| HET-HOM | *msh2 fancm* | 1399 | 198 | 194 | 854 | 153 | 33.70 |
| HET-HOM | *msh2 fancm* | 1746 | 232 | 239 | 1071 | 204 | 32.14 |
| HET-HOM | *msh2 fancm* | 1970 | 267 | 258 | 1193 | 252 | 31.66 |
| HET-HOM | *msh2 fancm* | 1298 | 164 | 175 | 786 | 173 | 30.89 |
| HET-HOM | *msh2 fancm* | 1007 | 125 | 136 | 621 | 125 | 30.60 |
| HOM-HET | *msh2 fancm* | 1493 | 199 | 245 | 894 | 155 | 36.34 |
| HOM-HET | *msh2 fancm* | 2411 | 329 | 384 | 1465 | 233 | 36.08 |
| HOM-HET | *msh2 fancm* | 2038 | 268 | 341 | 1232 | 197 | 36.57 |
| HOM-HET | *msh2 fancm* | 2273 | 336 | 378 | 1356 | 203 | 39.03 |
| HOM-HET | *msh2 fancm* | 2248 | 367 | 323 | 1323 | 235 | 37.86 |
| HOM-HET | *msh2 fancm* | 1975 | 352 | 322 | 1106 | 195 | 43.66 |
| HOM-HOM | *fancm zip4* | 1420 | 192 | 205 | 854 | 169 | 33.60 |
| HOM-HOM | *fancm zip4* | 1086 | 126 | 153 | 679 | 128 | 30.27 |
| HOM-HOM | *fancm zip4* | 1434 | 200 | 199 | 874 | 161 | 33.40 |
| HOM-HOM | *fancm zip4* | 1473 | 195 | 200 | 913 | 165 | 31.91 |
| HOM-HOM | *fancm zip4* | 1434 | 191 | 206 | 872 | 165 | 33.19 |
| HOM-HOM | *fancm zip4* | 1410 | 186 | 215 | 851 | 158 | 34.33 |

| | | | | | | | |
|---|---|---|---|---|---|---|---|
| HOM-HOM | *fancm zip4* | 1651 | 246 | 229 | 989 | 187 | 34.84 |
| HOM-HOM | *fancm zip4* | 1506 | 220 | 222 | 912 | 152 | 35.73 |
| HOM-HOM | *fancm zip4* | 1154 | 155 | 161 | 710 | 128 | 32.74 |
| HOM-HOM | *fancm zip4* | 1099 | 165 | 151 | 679 | 104 | 34.81 |
| HOM-HOM | *fancm zip4* | 1381 | 184 | 221 | 829 | 147 | 35.70 |
| HOM-HOM | *fancm zip4* | 1854 | 252 | 265 | 1125 | 212 | 33.50 |
| HOM-HOM | *fancm zip4* | 1215 | 164 | 184 | 730 | 137 | 34.64 |
| HOM-HOM | *fancm zip4* | 981 | 129 | 135 | 612 | 105 | 32.05 |
| HOM-HOM | *fancm zip4* | 1743 | 261 | 232 | 1049 | 201 | 34.10 |
| HOM-HOM | *fancm zip4* | 2098 | 287 | 326 | 1272 | 213 | 35.53 |
| HOM-HOM | *fancm zip4* | 993 | 125 | 154 | 600 | 114 | 33.81 |
| HET-HET | *fancm zip4* | 1307 | 67 | 96 | 922 | 222 | 13.36 |
| HET-HET | *fancm zip4* | 1921 | 183 | 141 | 1270 | 327 | 18.60 |
| HET-HET | *fancm zip4* | 1967 | 124 | 141 | 1364 | 338 | 14.53 |
| HET-HET | *fancm zip4* | 1261 | 101 | 110 | 840 | 210 | 18.43 |
| HET-HET | *fancm zip4* | 1778 | 162 | 110 | 1206 | 300 | 16.69 |
| HET-HET | *fancm zip4* | 1487 | 88 | 112 | 1005 | 282 | 14.50 |
| HET-HET | *fancm zip4* | 1394 | 106 | 117 | 917 | 254 | 17.53 |
| HET-HET | *fancm zip4* | 1355 | 90 | 100 | 931 | 234 | 15.17 |
| HET-HET | *fancm zip4* | 1719 | 132 | 131 | 1176 | 280 | 16.69 |
| HET-HET | *fancm zip4* | 1446 | 115 | 101 | 989 | 241 | 16.26 |
| HET-HET | *fancm zip4* | 1826 | 148 | 139 | 1206 | 333 | 17.20 |
| HET-HET | *fancm zip4* | 1775 | 144 | 140 | 1178 | 313 | 17.54 |
| HET-HOM | *fancm zip4* | 2236 | 169 | 176 | 1530 | 361 | 16.85 |
| HET-HOM | *fancm zip4* | 1985 | 95 | 111 | 1396 | 383 | 10.98 |
| HET-HOM | *fancm zip4* | 1684 | 111 | 125 | 1150 | 298 | 15.16 |
| HET-HOM | *fancm zip4* | 2245 | 159 | 133 | 1532 | 421 | 13.98 |
| HET-HOM | *fancm zip4* | 1560 | 87 | 100 | 1095 | 278 | 12.81 |
| HET-HOM | *fancm zip4* | 1808 | 103 | 88 | 1280 | 337 | 11.19 |
| HET-HOM | *fancm zip4* | 1909 | 146 | 121 | 1299 | 343 | 15.13 |
| HET-HOM | *fancm zip4* | 1458 | 110 | 116 | 967 | 265 | 16.93 |
| HET-HOM | *fancm zip4* | 2173 | 175 | 144 | 1471 | 383 | 15.95 |
| HET-HOM | *fancm zip4* | 1321 | 70 | 82 | 922 | 247 | 12.26 |
| HET-HOM | *fancm zip4* | 311 | 15 | 16 | 235 | 45 | 10.52 |
| HET-HOM | *fancm zip4* | 1823 | 117 | 137 | 1231 | 338 | 15.07 |
| HET-HOM | *fancm zip4* | 1872 | 99 | 133 | 1285 | 355 | 13.27 |
| HET-HOM | *fancm zip4* | 1166 | 96 | 68 | 798 | 204 | 15.22 |
| HET-HOM | *fancm zip4* | 877 | 56 | 60 | 613 | 148 | 14.24 |
| HOM-HET | *fancm zip4* | 2164 | 334 | 307 | 1282 | 241 | 36.16 |
| HOM-HET | *fancm zip4* | 1662 | 244 | 255 | 1004 | 159 | 36.79 |
| HOM-HET | *fancm zip4* | 1889 | 284 | 304 | 1098 | 203 | 38.56 |

| | | | | | | | |
|---|---|---|---|---|---|---|---|
| HOM-HET | *fancm zip4* | 806 | 125 | 119 | 488 | 74 | 37.19 |
| HOM-HET | *fancm zip4* | 935 | 129 | 148 | 569 | 89 | 36.17 |
| HOM-HET | *fancm zip4* | 1751 | 256 | 264 | 1056 | 175 | 36.28 |
| HOM-HET | *fancm zip4* | 1985 | 285 | 311 | 1177 | 212 | 36.79 |
| HOM-HET | *fancm zip4* | 2010 | 303 | 280 | 1235 | 192 | 35.20 |
| HOM-HET | *fancm zip4* | 1517 | 242 | 217 | 906 | 152 | 37.16 |
| HOM-HET | *fancm zip4* | 728 | 114 | 121 | 427 | 66 | 40.47 |
| HOM-HET | *fancm zip4* | 1992 | 286 | 281 | 1240 | 185 | 34.37 |
| HOM-HET | *fancm zip4* | 1978 | 294 | 315 | 1164 | 205 | 38.01 |
| HOM-HET | *fancm zip4* | 1944 | 276 | 336 | 1152 | 180 | 39.14 |
| HOM-HOM | *msh2 fancm zip4* | 1707 | 208 | 244 | 1065 | 190 | 31.41 |
| HOM-HOM | *msh2 fancm zip4* | 2373 | 307 | 349 | 1466 | 251 | 33.13 |
| HOM-HOM | *msh2 fancm zip4* | 1857 | 242 | 296 | 1128 | 191 | 35.15 |
| HOM-HOM | *msh2 fancm zip4* | 1681 | 231 | 213 | 1052 | 185 | 31.32 |
| HOM-HOM | *msh2 fancm zip4* | 1824 | 244 | 268 | 1131 | 181 | 33.77 |
| HOM-HOM | *msh2 fancm zip4* | 1850 | 261 | 264 | 1097 | 228 | 34.24 |
| HOM-HOM | *msh2 fancm zip4* | 1542 | 215 | 210 | 964 | 153 | 33.01 |
| HOM-HOM | *msh2 fancm zip4* | 1946 | 278 | 271 | 1154 | 243 | 33.99 |
| HOM-HOM | *msh2 fancm zip4* | 1878 | 255 | 272 | 1151 | 200 | 33.76 |
| HOM-HOM | *msh2 fancm zip4* | 1792 | 239 | 247 | 1127 | 179 | 32.35 |
| HOM-HOM | *msh2 fancm zip4* | 2267 | 333 | 283 | 1378 | 273 | 32.43 |
| HOM-HOM | *msh2 fancm zip4* | 1914 | 277 | 230 | 1194 | 213 | 31.43 |
| HET-HET | *msh2 fancm zip4* | 2204 | 250 | 250 | 1374 | 330 | 26.09 |
| HET-HET | *msh2 fancm zip4* | 1998 | 227 | 196 | 1270 | 305 | 24.07 |
| HET-HET | *msh2 fancm zip4* | 1574 | 215 | 209 | 952 | 198 | 32.08 |
| HET-HET | *msh2 fancm zip4* | 1805 | 162 | 205 | 1178 | 260 | 22.97 |
| HET-HET | *msh2 fancm zip4* | 2079 | 266 | 231 | 1313 | 269 | 27.76 |
| HET-HET | *msh2 fancm zip4* | 1956 | 206 | 197 | 1287 | 266 | 23.32 |
| HET-HET | *msh2 fancm zip4* | 1714 | 215 | 228 | 1040 | 231 | 30.50 |
| HET-HET | *msh2 fancm zip4* | 1668 | 214 | 203 | 1014 | 237 | 29.29 |
| HET-HET | *msh2 fancm zip4* | 1857 | 241 | 218 | 1145 | 253 | 28.89 |
| HET-HET | *msh2 fancm zip4* | 1617 | 188 | 174 | 1010 | 245 | 25.69 |
| HET-HET | *msh2 fancm zip4* | 1341 | 157 | 142 | 840 | 202 | 25.56 |
| HET-HET | *msh2 fancm zip4* | 1254 | 160 | 164 | 773 | 157 | 30.48 |
| HET-HET | *msh2 fancm zip4* | 1360 | 142 | 180 | 849 | 189 | 27.44 |
| HET-HET | *msh2 fancm zip4* | 964 | 122 | 103 | 596 | 143 | 26.98 |
| HET-HET | *msh2 fancm zip4* | 1356 | 164 | 170 | 823 | 199 | 28.77 |
| HET-HET | *msh2 fancm zip4* | 2013 | 229 | 263 | 1231 | 290 | 28.50 |
| HET-HET | *msh2 fancm zip4* | 1640 | 187 | 213 | 1014 | 226 | 28.43 |
| HET-HET | *msh2 fancm zip4* | 1150 | 146 | 137 | 702 | 165 | 28.74 |
| HET-HET | *msh2 fancm zip4* | 1623 | 182 | 177 | 1018 | 246 | 25.33 |

| | | | | | | | |
|---|---|---|---|---|---|---|---|
| HET-HET | *msh2 fancm zip4* | 2104 | 311 | 255 | 1271 | 267 | 32.03 |
| HET-HET | *msh2 fancm zip4* | 1984 | 250 | 221 | 1256 | 257 | 27.53 |
| HET-HET | *msh2 fancm zip4* | 1984 | 271 | 246 | 1192 | 275 | 30.80 |
| HET-HET | *msh2 fancm zip4* | 1984 | 220 | 249 | 1265 | 250 | 27.39 |
| HET-HET | *msh2 fancm zip4* | 2015 | 235 | 250 | 1221 | 309 | 27.99 |
| HET-HOM | *msh2 fancm zip4* | 1648 | 169 | 174 | 1078 | 227 | 23.60 |
| HET-HOM | *msh2 fancm zip4* | 2107 | 232 | 209 | 1364 | 302 | 23.75 |
| HET-HOM | *msh2 fancm zip4* | 2107 | 237 | 224 | 1330 | 316 | 25.01 |
| HET-HOM | *msh2 fancm zip4* | 970 | 96 | 106 | 620 | 148 | 23.61 |
| HET-HOM | *msh2 fancm zip4* | 2273 | 230 | 248 | 1428 | 367 | 23.88 |
| HET-HOM | *msh2 fancm zip4* | 2009 | 222 | 201 | 1274 | 312 | 23.91 |
| HET-HOM | *msh2 fancm zip4* | 1341 | 157 | 144 | 867 | 173 | 25.77 |
| HET-HOM | *msh2 fancm zip4* | 1727 | 185 | 197 | 1083 | 262 | 25.33 |
| HET-HOM | *msh2 fancm zip4* | 1214 | 124 | 154 | 763 | 173 | 26.38 |
| HET-HOM | *msh2 fancm zip4* | 1991 | 199 | 209 | 1256 | 327 | 23.18 |
| HET-HOM | *msh2 fancm zip4* | 1864 | 176 | 198 | 1197 | 293 | 22.62 |
| HET-HOM | *msh2 fancm zip4* | 2219 | 225 | 225 | 1414 | 355 | 22.90 |
| HET-HOM | *msh2 fancm zip4* | 1038 | 107 | 107 | 678 | 146 | 23.34 |
| HET-HOM | *msh2 fancm zip4* | 2099 | 239 | 215 | 1347 | 298 | 24.67 |
| HET-HOM | *msh2 fancm zip4* | 2209 | 222 | 210 | 1422 | 355 | 21.97 |
| HET-HOM | *msh2 fancm zip4* | 2115 | 227 | 228 | 1346 | 314 | 24.52 |
| HET-HOM | *msh2 fancm zip4* | 1845 | 189 | 191 | 1192 | 273 | 23.31 |
| HOM-HET | *msh2 fancm zip4* | 1813 | 284 | 289 | 1054 | 186 | 39.35 |
| HOM-HET | *msh2 fancm zip4* | 2292 | 343 | 337 | 1388 | 224 | 36.23 |
| HOM-HET | *msh2 fancm zip4* | 1982 | 292 | 294 | 1183 | 213 | 36.07 |
| HOM-HET | *msh2 fancm zip4* | 745 | 111 | 108 | 444 | 82 | 35.81 |
| HOM-HET | *msh2 fancm zip4* | 1869 | 312 | 291 | 1083 | 183 | 40.44 |
| HOM-HET | *msh2 fancm zip4* | 1934 | 269 | 284 | 1179 | 202 | 34.57 |
| HOM-HET | *msh2 fancm zip4* | 953 | 146 | 136 | 571 | 100 | 36.11 |
| HOM-HET | *msh2 fancm zip4* | 1411 | 206 | 223 | 832 | 150 | 37.40 |
| HOM-HET | *msh2 fancm zip4* | 1471 | 215 | 222 | 873 | 161 | 36.29 |
| HOM-HET | *msh2 fancm zip4* | 1670 | 273 | 293 | 949 | 155 | 43.24 |
| HOM-HET | *msh2 fancm zip4* | 1089 | 174 | 152 | 649 | 114 | 36.65 |

**Supplementary Table 8.** *420* fluorescent seed count data for the lines with *HEI10* overexpression (*HEI10-OE*). *HEI10-OE* construct was introduced into wild type*, msh2* and *msh2 fancm zip4* mutants in recombinant lines with differing pattern of heterozygosity. The following recombinant backgrounds were used: 'HOM-HOM' that are Col/Col inbred throughout the genome, 'HET-HET' that are Col/Ct heterozygous throughout the genome, 'HET-HOM' where the *420* region is Col/Ct heterozygous and the remainder of chromosome 3 is Col/Col homozygous, and 'HOM-HET' where *420* is Col/Col homozygous and the remainder of chr3 is Col/Ct heterozygous. Genetic distance in centimorgans (cM) is calculated as $cM = 100\times(1-(1-2(NG+NR)/NT)^{1/2})$, where NG is the number of green alone seeds, NR is the number of red alone seeds and NT is the total number of seeds of all classes analysed.

| Cross | Genotype | Total | Green alone | Red alone | Red+ Green | Non-colour | cM |
|---|---|---|---|---|---|---|---|
| HOM-HOM | Wild type | 3746 | 330 | 361 | 2374 | 681 | 20.56 |
| HOM-HOM | Wild type | 3492 | 351 | 343 | 2268 | 530 | 22.38 |
| HOM-HOM | Wild type | 3964 | 338 | 349 | 2629 | 648 | 19.17 |
| HOM-HOM | Wild type | 3983 | 347 | 417 | 2548 | 671 | 21.49 |
| HOM-HOM | Wild type | 3225 | 324 | 276 | 2089 | 536 | 20.76 |
| HOM-HOM | Wild type | 4623 | 385 | 412 | 3069 | 757 | 19.06 |
| HOM-HOM | Wild type | 868 | 73 | 80 | 579 | 136 | 19.53 |
| HOM-HOM | Wild type | 3586 | 324 | 321 | 2305 | 636 | 19.98 |
| HOM-HOM | Wild type | 3880 | 381 | 313 | 2519 | 667 | 19.86 |
| HOM-HOM | *HEI10-OE* | 1742 | 310 | 267 | 1006 | 159 | 41.90 |
| HOM-HOM | *HEI10-OE* | 3795 | 668 | 611 | 2181 | 335 | 42.91 |
| HOM-HOM | *HEI10-OE* | 2044 | 313 | 325 | 1201 | 205 | 38.70 |
| HOM-HOM | *HEI10-OE* | 3514 | 591 | 535 | 2048 | 340 | 40.07 |
| HOM-HOM | *HEI10-OE* | 3586 | 630 | 546 | 2085 | 325 | 41.34 |
| HOM-HOM | *HEI10-OE* | 1507 | 218 | 229 | 889 | 171 | 36.22 |
| HOM-HOM | *HEI10-OE* | 2060 | 335 | 363 | 1180 | 182 | 43.23 |
| HOM-HOM | *HEI10-OE* | 2193 | 377 | 346 | 1260 | 210 | 41.64 |
| HOM-HOM | *HEI10-OE* | 2408 | 440 | 381 | 1391 | 196 | 43.60 |
| HOM-HOM | *HEI10-OE* | 2362 | 391 | 392 | 1371 | 208 | 41.95 |
| HOM-HOM | *HEI10-OE* | 2197 | 381 | 368 | 1259 | 189 | 43.59 |
| HOM-HOM | *HEI10-OE* | 2293 | 385 | 390 | 1304 | 214 | 43.08 |
| HOM-HOM | *HEI10-OE* | 2268 | 369 | 379 | 1313 | 207 | 41.66 |
| HOM-HOM | *HEI10-OE* | 2256 | 367 | 384 | 1304 | 201 | 42.19 |
| HOM-HOM | *HEI10-OE* | 2247 | 400 | 369 | 1298 | 180 | 43.83 |
| HOM-HOM | *HEI10-OE* | 2248 | 390 | 357 | 1305 | 196 | 42.09 |
| HOM-HOM | *HEI10-OE* | 2318 | 361 | 412 | 1367 | 178 | 42.29 |
| HOM-HOM | *msh2 HEI10-OE* | 1229 | 185 | 183 | 754 | 107 | 36.66 |
| HOM-HOM | *msh2 HEI10-OE* | 2014 | 276 | 369 | 1177 | 192 | 40.04 |
| HOM-HOM | *msh2 HEI10-OE* | 2126 | 322 | 294 | 1301 | 209 | 35.15 |
| HOM-HOM | *msh2 HEI10-OE* | 2094 | 318 | 302 | 1257 | 217 | 36.14 |

| | | | | | | | |
|---|---|---|---|---|---|---|---|
| HOM-HOM | *msh2 HEI10-OE* | 1723 | 237 | 287 | 1029 | 170 | 37.41 |
| HOM-HOM | *msh2 HEI10-OE* | 2045 | 307 | 313 | 1198 | 227 | 37.26 |
| HOM-HOM | *msh2 HEI10-OE* | 1259 | 191 | 194 | 742 | 132 | 37.68 |
| HET-HET | Wild type | 4260 | 373 | 328 | 2825 | 734 | 18.09 |
| HET-HET | Wild type | 3519 | 273 | 266 | 2335 | 645 | 16.71 |
| HET-HET | Wild type | 4099 | 318 | 324 | 2736 | 721 | 17.13 |
| HET-HET | Wild type | 4302 | 400 | 326 | 2833 | 743 | 18.61 |
| HET-HET | Wild type | 3346 | 257 | 239 | 2194 | 656 | 16.12 |
| HET-HET | Wild type | 2056 | 191 | 155 | 1355 | 355 | 18.55 |
| HET-HET | Wild type | 3556 | 277 | 327 | 2345 | 607 | 18.74 |
| HET-HET | Wild type | 3718 | 295 | 321 | 2433 | 669 | 18.23 |
| HET-HET | Wild type | 1869 | 142 | 158 | 1234 | 335 | 17.60 |
| HET-HET | Wild type | 3390 | 293 | 316 | 2272 | 509 | 19.96 |
| HET-HET | Wild type | 2718 | 210 | 237 | 1810 | 461 | 18.08 |
| HET-HET | Wild type | 3744 | 303 | 308 | 2454 | 679 | 17.93 |
| HET-HET | *HEI10-OE* | 3842 | 624 | 549 | 2283 | 386 | 37.60 |
| HET-HET | *HEI10-OE* | 3782 | 646 | 562 | 2185 | 389 | 39.90 |
| HET-HET | *HEI10-OE* | 3827 | 540 | 626 | 2223 | 438 | 37.50 |
| HET-HET | *HEI10-OE* | 2518 | 438 | 377 | 1446 | 257 | 40.61 |
| HET-HET | *HEI10-OE* | 2352 | 336 | 358 | 1432 | 226 | 35.98 |
| HET-HET | *HEI10-OE* | 1537 | 233 | 252 | 902 | 150 | 39.26 |
| HET-HET | *HEI10-OE* | 3240 | 448 | 541 | 1902 | 349 | 37.59 |
| HET-HET | *HEI10-OE* | 3108 | 513 | 453 | 1844 | 298 | 38.49 |
| HET-HET | *HEI10-OE* | 3793 | 603 | 596 | 2159 | 435 | 39.35 |
| HET-HET | *msh2 HEI10-OE* | 2118 | 305 | 280 | 1263 | 270 | 33.10 |
| HET-HET | *msh2 HEI10-OE* | 2156 | 370 | 309 | 1277 | 200 | 39.16 |
| HET-HET | *msh2 HEI10-OE* | 2058 | 323 | 335 | 1170 | 230 | 39.95 |
| HET-HET | *msh2 HEI10-OE* | 1458 | 237 | 246 | 829 | 146 | 41.91 |
| HET-HET | *msh2 HEI10-OE* | 2139 | 352 | 318 | 1243 | 226 | 38.88 |
| HET-HET | *msh2 HEI10-OE* | 1520 | 223 | 213 | 912 | 172 | 34.71 |
| HET-HET | *msh2 HEI10-OE* | 1213 | 168 | 203 | 724 | 118 | 37.69 |
| HET-HET | *msh2 fancm zip4 HEI10-OE* | 2036 | 273 | 240 | 1245 | 278 | 29.57 |
| HET-HET | *msh2 fancm zip4 HEI10-OE* | 1688 | 227 | 181 | 1055 | 225 | 28.13 |
| HET-HET | *msh2 fancm zip4 HEI10-OE* | 1939 | 239 | 208 | 1212 | 280 | 26.59 |
| HET-HOM | Wild type | 3896 | 475 | 422 | 2396 | 603 | 26.55 |
| HET-HOM | Wild type | 4073 | 446 | 392 | 2628 | 607 | 23.29 |
| HET-HOM | Wild type | 3741 | 458 | 410 | 2332 | 541 | 26.79 |
| HET-HOM | Wild type | 3841 | 416 | 410 | 2380 | 635 | 24.51 |

| | | | | | | | |
|---|---|---|---|---|---|---|---|
| HET-HOM | Wild type | 3931 | 416 | 462 | 2468 | 585 | 25.62 |
| HET-HOM | Wild type | 3376 | 366 | 376 | 2177 | 457 | 25.14 |
| HET-HOM | Wild type | 3997 | 462 | 438 | 2509 | 588 | 25.86 |
| HET-HOM | Wild type | 3037 | 357 | 345 | 1896 | 439 | 26.67 |
| HET-HOM | Wild type | 3652 | 430 | 432 | 2269 | 521 | 27.34 |
| HET-HOM | Wild type | 3532 | 402 | 358 | 2232 | 540 | 24.52 |
| HET-HOM | Wild type | 1411 | 142 | 136 | 940 | 193 | 22.16 |
| HET-HOM | Wild type | 3504 | 351 | 411 | 2196 | 546 | 24.83 |
| HET-HOM | *HEI10-OE* | 2047 | 380 | 362 | 1132 | 173 | 47.56 |
| HET-HOM | *HEI10-OE* | 1886 | 324 | 336 | 1086 | 140 | 45.22 |
| HET-HOM | *HEI10-OE* | 1944 | 360 | 317 | 1099 | 168 | 44.91 |
| HET-HOM | *HEI10-OE* | 1481 | 300 | 238 | 823 | 120 | 47.71 |
| HET-HOM | *HEI10-OE* | 1432 | 263 | 236 | 815 | 118 | 44.95 |
| HET-HOM | *HEI10-OE* | 1967 | 348 | 327 | 1125 | 167 | 43.99 |
| HET-HOM | *HEI10-OE* | 1883 | 342 | 338 | 1061 | 142 | 47.30 |
| HET-HOM | *HEI10-OE* | 1744 | 310 | 304 | 966 | 164 | 45.61 |
| HET-HOM | *HEI10-OE* | 3824 | 737 | 662 | 2175 | 250 | 48.20 |
| HET-HOM | *HEI10-OE* | 4103 | 729 | 721 | 2263 | 390 | 45.85 |
| HET-HOM | *HEI10-OE* | 4061 | 730 | 749 | 2187 | 395 | 47.88 |
| HET-HOM | *msh2 HEI10-OE* | 1887 | 303 | 263 | 1131 | 190 | 36.75 |
| HET-HOM | *msh2 HEI10-OE* | 1512 | 247 | 235 | 877 | 153 | 39.80 |
| HET-HOM | *msh2 HEI10-OE* | 1546 | 240 | 228 | 933 | 145 | 37.19 |
| HET-HOM | *msh2 HEI10-OE* | 1207 | 191 | 192 | 721 | 103 | 39.55 |
| HET-HOM | *msh2 HEI10-OE* | 1892 | 278 | 258 | 1144 | 212 | 34.17 |
| HET-HOM | *msh2 HEI10-OE* | 1377 | 208 | 158 | 852 | 159 | 31.56 |
| HET-HOM | *msh2 HEI10-OE* | 2049 | 274 | 295 | 1243 | 237 | 33.32 |
| HET-HOM | *msh2 fancm zip4 HEI10-OE* | 1622 | 177 | 165 | 1063 | 217 | 23.95 |
| HET-HOM | *msh2 fancm zip4 HEI10-OE* | 1884 | 201 | 185 | 1243 | 255 | 23.17 |
| HET-HOM | *msh2 fancm zip4 HEI10-OE* | 1391 | 152 | 168 | 900 | 171 | 26.52 |
| HET-HOM | *msh2 fancm zip4 HEI10-OE* | 1936 | 208 | 235 | 1227 | 266 | 26.36 |
| HET-HOM | *msh2 fancm zip4 HEI10-OE* | 2138 | 214 | 220 | 1415 | 289 | 22.93 |
| HET-HOM | *msh2 fancm zip4 HEI10-OE* | 1318 | 159 | 153 | 812 | 194 | 27.44 |
| HET-HOM | *msh2 fancm zip4 HEI10-OE* | 1225 | 126 | 135 | 809 | 155 | 24.25 |
| HET-HOM | *msh2 fancm zip4 HEI10-OE* | 1968 | 239 | 219 | 1256 | 254 | 26.89 |
| HOM-HET | Wild type | 3101 | 148 | 155 | 2110 | 688 | 10.30 |
| HOM-HET | Wild type | 3736 | 244 | 258 | 2491 | 743 | 14.49 |

| | | | | | | | |
|---|---|---|---|---|---|---|---|
| HOM-HET | Wild type | 1852 | 123 | 134 | 1265 | 330 | 15.00 |
| HOM-HET | Wild type | 3188 | 216 | 197 | 2168 | 607 | 13.92 |
| HOM-HET | Wild type | 3614 | 194 | 198 | 2475 | 747 | 11.51 |
| HOM-HET | Wild type | 3441 | 175 | 155 | 2343 | 768 | 10.10 |
| HOM-HET | Wild type | 720 | 51 | 54 | 471 | 144 | 15.84 |
| HOM-HET | Wild type | 2567 | 142 | 171 | 1737 | 517 | 13.04 |
| HOM-HET | Wild type | 3457 | 189 | 234 | 2345 | 689 | 13.09 |
| HOM-HET | Wild type | 1692 | 79 | 131 | 1152 | 330 | 13.30 |
| HOM-HET | Wild type | 3643 | 180 | 212 | 2561 | 690 | 11.41 |
| HOM-HET | Wild type | 2652 | 177 | 167 | 1762 | 546 | 13.94 |
| HOM-HET | Wild type | 2652 | 177 | 167 | 1762 | 546 | 13.94 |
| HOM-HET | Wild type | 3562 | 190 | 210 | 2411 | 751 | 11.94 |
| HOM-HET | Wild type | 2794 | 149 | 136 | 1901 | 608 | 10.78 |
| HOM-HET | Wild type | 3877 | 211 | 233 | 2651 | 782 | 12.20 |
| HOM-HET | Wild type | 3316 | 208 | 199 | 2235 | 674 | 13.14 |
| HOM-HET | *HEI10-OE* | 1480 | 232 | 167 | 917 | 164 | 32.12 |
| HOM-HET | *HEI10-OE* | 2229 | 287 | 294 | 1420 | 228 | 30.81 |
| HOM-HET | *HEI10-OE* | 1621 | 204 | 209 | 1011 | 197 | 29.97 |
| HOM-HET | *HEI10-OE* | 1658 | 205 | 215 | 999 | 239 | 29.76 |
| HOM-HET | *HEI10-OE* | 1612 | 191 | 198 | 1030 | 193 | 28.07 |
| HOM-HET | *HEI10-OE* | 1600 | 188 | 199 | 1008 | 205 | 28.15 |
| HOM-HET | *HEI10-OE* | 1044 | 108 | 131 | 659 | 146 | 26.37 |
| HOM-HET | *HEI10-OE* | 969 | 108 | 105 | 638 | 118 | 25.14 |
| HOM-HET | *HEI10-OE* | 2028 | 269 | 234 | 1270 | 255 | 29.01 |
| HOM-HET | *HEI10-OE* | 1767 | 219 | 221 | 1114 | 213 | 29.15 |
| HOM-HET | *HEI10-OE* | 1882 | 250 | 260 | 1173 | 199 | 32.32 |
| HOM-HET | *HEI10-OE* | 1805 | 233 | 247 | 1112 | 213 | 31.58 |
| HOM-HET | *HEI10-OE* | 2078 | 243 | 269 | 1314 | 252 | 28.78 |
| HOM-HET | *HEI10-OE* | 870 | 120 | 102 | 552 | 96 | 30.02 |
| HOM-HET | *HEI10-OE* | 2109 | 272 | 251 | 1345 | 241 | 29.00 |
| HOM-HET | *HEI10-OE* | 1800 | 184 | 235 | 1152 | 229 | 26.89 |
| HOM-HET | *HEI10-OE* | 860 | 99 | 116 | 543 | 102 | 29.29 |
| HOM-HET | *HEI10-OE* | 4093 | 548 | 538 | 2505 | 502 | 31.49 |
| HOM-HET | *HEI10-OE* | 3123 | 373 | 415 | 1916 | 419 | 29.62 |
| HOM-HET | *HEI10-OE* | 3337 | 425 | 405 | 2078 | 429 | 29.11 |
| HOM-HET | *HEI10-OE* | 2110 | 246 | 252 | 1339 | 273 | 27.34 |
| HOM-HET | *HEI10-OE* | 2695 | 369 | 303 | 1678 | 345 | 29.20 |
| HOM-HET | *HEI10-OE* | 4172 | 481 | 562 | 2544 | 585 | 29.29 |
| HOM-HET | *HEI10-OE* | 3797 | 525 | 519 | 2275 | 478 | 32.91 |
| HOM-HET | *HEI10-OE* | 3965 | 521 | 493 | 2392 | 559 | 30.11 |

| | | | | | | | |
|---|---|---|---|---|---|---|---|
| HOM-HET | *HEI10-OE* | 4129 | 539 | 532 | 2581 | 477 | 30.63 |
| HOM-HET | *HEI10-OE* | 4121 | 510 | 588 | 2579 | 444 | 31.65 |
| HOM-HET | *HEI10-OE* | 4128 | 491 | 594 | 2563 | 480 | 31.13 |
| HOM-HET | *HEI10-OE* | 4083 | 561 | 625 | 2419 | 478 | 35.27 |
| HOM-HET | *HEI10-OE* | 4006 | 468 | 561 | 2453 | 524 | 30.27 |
| HOM-HET | *msh2 HEI10-OE* | 1481 | 219 | 221 | 917 | 124 | 36.30 |
| HOM-HET | *msh2 HEI10-OE* | 1397 | 220 | 235 | 804 | 138 | 40.96 |
| HOM-HET | *msh2 HEI10-OE* | 2004 | 295 | 335 | 1177 | 197 | 39.07 |
| HOM-HET | *msh2 HEI10-OE* | 1560 | 241 | 279 | 910 | 130 | 42.26 |
| HOM-HET | *msh2 HEI10-OE* | 2073 | 294 | 344 | 1251 | 184 | 37.99 |
| HOM-HET | *msh2 HEI10-OE* | 2012 | 322 | 309 | 1169 | 212 | 38.95 |
| HOM-HET | *msh2 HEI10-OE* | 1875 | 333 | 292 | 1084 | 166 | 42.26 |
| HOM-HET | *msh2 HEI10-OE* | 2057 | 305 | 341 | 1210 | 201 | 39.02 |
| HOM-HET | *msh2 HEI10-OE* | 2059 | 341 | 313 | 1195 | 210 | 39.61 |
| HOM-HET | *msh2 HEI10-OE* | 1794 | 309 | 301 | 1012 | 172 | 43.44 |

**Supplementary Table 9.** *3.9* fluorescent seed count data for the wild type. *fancm zip4* and *msh2 fancm zip4* mutants in recombinant lines with differing pattern of heterozygosity. The following recombinant backgrounds were used: 'HOM-HOM' that are Col/Col inbred throughout the genome; 'HET-HET' that are Col/Ct heterozygous throughout the genome; ''HET-HOM' where the subtelomeric part of chromosome 3 is Col/Ct heterozygous and the remainder of chromosome 3 is Col/Col homozygous. Genetic distance in centimorgans (cM) is calculated as $cM = 100\times(1-(1-2(NG+NR)/NT)^{1/2})$, where NG is the number of green alone seeds, NR is the number of red alone seeds and NT is the total number of seeds of all classes analysed.

| | Genotype | Total | Green alone | Red alone | Red+ Green | Non-colour | cM |
|---|---|---|---|---|---|---|---|
| HOM-HOM | Wild type | 2376 | 211 | 155 | 1597 | 413 | 16.82 |
| HOM-HOM | Wild type | 986 | 70 | 68 | 676 | 172 | 15.14 |
| HOM-HOM | Wild type | 2386 | 208 | 167 | 1600 | 411 | 17.20 |
| HOM-HOM | Wild type | 2602 | 202 | 230 | 1716 | 454 | 18.27 |
| HOM-HOM | Wild type | 2619 | 223 | 206 | 1781 | 409 | 18.00 |
| HOM-HOM | Wild type | 2620 | 213 | 217 | 1724 | 466 | 18.04 |
| HOM-HOM | Wild type | 2475 | 210 | 184 | 1657 | 424 | 17.44 |
| HOM-HOM | Wild type | 2499 | 185 | 199 | 1683 | 432 | 16.77 |
| HOM-HOM | Wild type | 2377 | 226 | 157 | 1585 | 409 | 17.67 |
| HOM-HOM | Wild type | 2479 | 208 | 170 | 1641 | 460 | 16.63 |
| HOM-HOM | Wild type | 2416 | 208 | 201 | 1567 | 440 | 18.67 |
| HOM-HOM | Wild type | 2356 | 216 | 182 | 1538 | 420 | 18.63 |
| HET-HET | Wild type | 2098 | 210 | 186 | 1369 | 333 | 21.10 |
| HET-HET | Wild type | 1113 | 95 | 116 | 728 | 174 | 21.21 |
| HET-HET | Wild type | 2027 | 153 | 189 | 1365 | 320 | 18.60 |
| HET-HET | Wild type | 2395 | 199 | 230 | 1566 | 400 | 19.89 |
| HET-HET | Wild type | 2001 | 206 | 181 | 1293 | 321 | 21.69 |
| HET-HET | Wild type | 472 | 45 | 36 | 310 | 81 | 18.96 |
| HET-HOM | Wild type | 2315 | 169 | 203 | 1524 | 419 | 17.62 |
| HET-HOM | Wild type | 2497 | 195 | 142 | 1695 | 465 | 14.56 |
| HET-HOM | Wild type | 1890 | 138 | 129 | 1278 | 345 | 15.30 |
| HET-HOM | Wild type | 2201 | 156 | 159 | 1528 | 358 | 15.52 |
| HET-HOM | Wild type | 2540 | 187 | 208 | 1700 | 445 | 17.00 |
| HET-HOM | Wild type | 2324 | 164 | 149 | 1561 | 450 | 14.52 |
| HET-HOM | Wild type | 2481 | 179 | 150 | 1682 | 470 | 14.28 |
| HET-HOM | Wild type | 2537 | 217 | 181 | 1673 | 466 | 17.16 |
| HOM-HOM | *fancm zip4* | 1717 | 118 | 129 | 1168 | 302 | 15.60 |
| HOM-HOM | *fancm zip4* | 983 | 81 | 58 | 673 | 171 | 15.31 |
| HOM-HOM | *fancm zip4* | 2192 | 160 | 167 | 1466 | 399 | 16.24 |
| HOM-HOM | *fancm zip4* | 1834 | 126 | 160 | 1216 | 332 | 17.05 |
| HOM-HOM | *fancm zip4* | 2219 | 148 | 173 | 1529 | 369 | 15.70 |
| HOM-HOM | *fancm zip4* | 2230 | 163 | 138 | 1516 | 413 | 14.56 |
| HOM-HOM | *fancm zip4* | 2250 | 146 | 142 | 1546 | 416 | 13.74 |
| HOM-HOM | *fancm zip4* | 1539 | 82 | 109 | 1049 | 299 | 13.29 |
| HOM-HOM | *fancm zip4* | 2014 | 126 | 150 | 1373 | 365 | 14.80 |
| HOM-HOM | *fancm zip4* | 2057 | 129 | 142 | 1398 | 388 | 14.18 |
| HOM-HOM | *fancm zip4* | 2092 | 198 | 234 | 1340 | 320 | 23.38 |
| HOM-HOM | *fancm zip4* | 1380 | 80 | 79 | 955 | 266 | 12.28 |
| HOM-HOM | *fancm zip4* | 703 | 47 | 37 | 491 | 128 | 12.76 |
| HOM-HOM | *fancm zip4* | 1171 | 67 | 95 | 791 | 218 | 14.95 |

| | | | | | | | |
|---|---|---|---|---|---|---|---|
| HET-HET | *fancm zip4* | 1864 | 70 | 59 | 1302 | 433 | 7.18 |
| HET-HET | *fancm zip4* | 545 | 24 | 26 | 383 | 112 | 9.64 |
| HET-HET | *fancm zip4* | 831 | 35 | 28 | 587 | 181 | 7.89 |
| HET-HET | *fancm zip4* | 1904 | 67 | 47 | 1348 | 442 | 6.18 |
| HET-HET | *fancm zip4* | 1849 | 56 | 85 | 1326 | 382 | 7.94 |
| HET-HET | *fancm zip4* | 1857 | 41 | 52 | 1318 | 446 | 5.14 |
| HET-HET | *fancm zip4* | 1689 | 65 | 76 | 1191 | 357 | 8.73 |
| HET-HET | *fancm zip4* | 1956 | 62 | 49 | 1396 | 449 | 5.85 |
| HET-HET | *fancm zip4* | 904 | 28 | 32 | 643 | 201 | 6.87 |
| HET-HET | *fancm zip4* | 2096 | 56 | 56 | 1491 | 493 | 5.49 |
| HET-HET | *fancm zip4* | 964 | 32 | 45 | 677 | 210 | 8.33 |
| HET-HET | *fancm zip4* | 1322 | 59 | 47 | 948 | 268 | 8.37 |
| HET-HOM | *fancm zip4* | 2138 | 175 | 137 | 1444 | 382 | 15.85 |
| HET-HOM | *fancm zip4* | 2063 | 136 | 165 | 1382 | 380 | 15.85 |
| HET-HOM | *fancm zip4* | 1768 | 128 | 102 | 1194 | 344 | 13.99 |
| HET-HOM | *fancm zip4* | 2149 | 142 | 150 | 1436 | 421 | 14.66 |
| HET-HOM | *fancm zip4* | 2100 | 157 | 147 | 1441 | 355 | 15.71 |
| HET-HOM | *fancm zip4* | 1321 | 87 | 97 | 905 | 232 | 15.06 |
| HOM-HOM | *msh2 fancm zip4* | 1326 | 93 | 99 | 896 | 238 | 15.71 |
| HOM-HOM | *msh2 fancm zip4* | 1878 | 136 | 142 | 1281 | 319 | 16.10 |
| HOM-HOM | *msh2 fancm zip4* | 1837 | 122 | 139 | 1219 | 357 | 15.39 |
| HOM-HOM | *msh2 fancm zip4* | 1433 | 80 | 124 | 969 | 260 | 15.43 |
| HOM-HOM | *msh2 fancm zip4* | 2174 | 194 | 131 | 1439 | 410 | 16.27 |
| HOM-HOM | *msh2 fancm zip4* | 2221 | 138 | 144 | 1486 | 453 | 13.63 |
| HOM-HOM | *msh2 fancm zip4* | 2181 | 126 | 152 | 1489 | 414 | 13.68 |
| HOM-HOM | *msh2 fancm zip4* | 2341 | 157 | 162 | 1618 | 404 | 14.71 |
| HOM-HOM | *msh2 fancm zip4* | 1984 | 154 | 161 | 1304 | 365 | 17.39 |
| HOM-HOM | *msh2 fancm zip4* | 2052 | 154 | 114 | 1411 | 373 | 14.05 |
| HET-HET | *msh2 fancm zip4* | 1456 | 55 | 48 | 1044 | 309 | 7.34 |
| HET-HET | *msh2 fancm zip4* | 2022 | 41 | 38 | 1481 | 462 | 3.99 |
| HET-HET | *msh2 fancm zip4* | 1877 | 69 | 45 | 1333 | 430 | 6.27 |
| HET-HET | *msh2 fancm zip4* | 1959 | 48 | 58 | 1399 | 454 | 5.57 |
| HET-HET | *msh2 fancm zip4* | 2020 | 59 | 31 | 1489 | 441 | 4.56 |
| HET-HET | *msh2 fancm zip4* | 1671 | 48 | 53 | 1217 | 353 | 6.24 |
| HET-HET | *msh2 fancm zip4* | 1755 | 55 | 41 | 1244 | 415 | 5.63 |
| HET-HET | *msh2 fancm zip4* | 2166 | 102 | 41 | 1548 | 475 | 6.84 |
| HET-HET | *msh2 fancm zip4* | 2205 | 46 | 41 | 1611 | 507 | 4.03 |
| HET-HET | *msh2 fancm zip4* | 2051 | 67 | 51 | 1442 | 491 | 5.93 |
| HET-HOM | *msh2 fancm zip4* | 1918 | 106 | 123 | 1346 | 343 | 12.75 |
| HET-HOM | *msh2 fancm zip4* | 2015 | 139 | 149 | 1365 | 362 | 15.49 |
| HET-HOM | *msh2 fancm zip4* | 1777 | 117 | 111 | 1220 | 329 | 13.78 |
| HET-HOM | *msh2 fancm zip4* | 1679 | 139 | 118 | 1127 | 295 | 16.70 |
| HET-HOM | *msh2 fancm zip4* | 1697 | 111 | 117 | 1178 | 291 | 14.48 |
| HET-HOM | *msh2 fancm zip4* | 2303 | 168 | 146 | 1539 | 450 | 14.72 |
| HET-HOM | *msh2 fancm zip4* | 1890 | 108 | 157 | 1290 | 335 | 15.17 |
| HET-HOM | *msh2 fancm zip4* | 1098 | 72 | 59 | 757 | 210 | 12.74 |
| HET-HOM | *msh2 fancm zip4* | 2399 | 162 | 164 | 1675 | 398 | 14.66 |
| HET-HOM | *msh2 fancm zip4* | 2379 | 174 | 168 | 1600 | 437 | 15.59 |

**Supplementary Table 10.** *CEN3* crossover frequency in the wild type and *msh2* mutant in recombinant lines with differing pattern of heterozygosity. The following recombinant backgrounds were used: 'HOM-HOM' that are Col/Col inbred throughout the genome, 'HET-HET' that are Col/Ct heterozygous throughout the genome, 'HET-HOM' where the subtelomeric region of chromosome 3 is Col/Ct heterozygous and the remainder of chr3 is Col/Col homozygous, and 'HOM-HET' where subtelomeric region of chr3 is Col/Col homozygous and the remainder of chr3 is Col/Ct heterozygous. Genetic distance in centimorgans (cM) is calculated as cM = $100\times(N_Y/(N_Y+N_{R+Y}))$, where $N_Y$ is the number of yellow alone pollen, and $N_{Y+R}$ is the number of red and yellow pollen.

| Cross | Genotype | Total | Red alone | Yellow alone | Red+Yellow | Non-colour | cM |
|---|---|---|---|---|---|---|---|
| HOM-HOM | Wild type | 3402 | 264 | 147 | 1343 | 1648 | 10.94 |
| HOM-HOM | Wild type | 14293 | 937 | 606 | 6258 | 6492 | 9.68 |
| HOM-HOM | Wild type | 17145 | 993 | 794 | 7252 | 8106 | 10.94 |
| HOM-HOM | Wild type | 4356 | 375 | 215 | 1855 | 1911 | 11.59 |
| HOM-HOM | Wild type | 5689 | 383 | 255 | 2746 | 2305 | 9.28 |
| HOM-HOM | Wild type | 2113 | 99 | 65 | 694 | 1255 | 9.36 |
| HOM-HOM | Wild type | 2307 | 126 | 103 | 1071 | 1007 | 9.61 |
| HOM-HOM | *msh2* | 2200 | 169 | 100 | 915 | 1016 | 10.92 |
| HOM-HOM | *msh2* | 2295 | 191 | 94 | 893 | 1117 | 10.52 |
| HOM-HOM | *msh2* | 3229 | 252 | 154 | 1337 | 1486 | 11.51 |
| HOM-HOM | *msh2* | 2142 | 158 | 97 | 893 | 994 | 10.86 |
| HOM-HOM | *msh2* | 3242 | 231 | 169 | 1444 | 1398 | 11.71 |
| HOM-HOM | *msh2* | 1954 | 145 | 106 | 982 | 721 | 10.79 |
| HET-HET | Wild type | 3656 | 228 | 126 | 1237 | 2065 | 10.18 |
| HET-HET | Wild type | 3175 | 195 | 95 | 1095 | 1790 | 8.67 |
| HET-HET | Wild type | 13563 | 697 | 544 | 5800 | 6522 | 9.37 |
| HET-HET | Wild type | 13698 | 812 | 650 | 6598 | 5638 | 9.85 |
| HET-HET | Wild type | 12981 | 673 | 628 | 5840 | 5840 | 10.75 |
| HET-HET | Wild type | 2133 | 82 | 88 | 752 | 1211 | 11.71 |
| HET-HET | Wild type | 5593 | 282 | 180 | 2326 | 2805 | 7.73 |
| HET-HET | Wild type | 4044 | 213 | 103 | 1302 | 2426 | 7.91 |
| HET-HET | Wild type | 5573 | 260 | 188 | 1591 | 3534 | 11.81 |
| HET-HET | Wild type | 6126 | 596 | 255 | 2251 | 3024 | 11.32 |
| HET-HET | *msh2* | 14462 | 340 | 256 | 7080 | 6786 | 3.61 |
| HET-HET | *msh2* | 12150 | 290 | 210 | 5170 | 6480 | 4.06 |
| HET-HET | *msh2* | 3258 | 63 | 61 | 1443 | 1691 | 4.22 |
| HET-HET | *msh2* | 3984 | 105 | 116 | 1884 | 1879 | 6.15 |
| HET-HET | *msh2* | 3748 | 82 | 83 | 2030 | 1553 | 4.08 |
| HET-HET | *msh2* | 10291 | 235 | 240 | 6054 | 3762 | 3.96 |
| HET-HET | *msh2* | 6209 | 137 | 152 | 3148 | 2772 | 4.82 |
| HET-HET | *msh2* | 13477 | 296 | 319 | 8408 | 4454 | 3.79 |
| HET-HOM | Wild type | 3399 | 214 | 106 | 925 | 2154 | 11.45 |
| HET-HOM | Wild type | 2968 | 259 | 105 | 1092 | 1512 | 9.61 |
| HET-HOM | Wild type | 9094 | 498 | 365 | 4864 | 3367 | 7.51 |
| HET-HOM | Wild type | 9212 | 525 | 355 | 4840 | 3492 | 7.33 |

| | | | | | | | |
|---|---|---|---|---|---|---|---|
| HET-HOM | Wild type | 8893 | 622 | 302 | 3465 | 4504 | 8.71 |
| HET-HOM | *msh2* | 8933 | 608 | 325 | 4221 | 3779 | 7.69 |
| HET-HOM | *msh2* | 17666 | 1443 | 656 | 6950 | 8617 | 9.43 |
| HET-HOM | *msh2* | 3958 | 278 | 162 | 1459 | 2059 | 11.11 |
| HET-HOM | *msh2* | 6731 | 649 | 255 | 3395 | 2432 | 7.51 |
| HET-HOM | *msh2* | 4086 | 275 | 186 | 1958 | 1667 | 9.49 |
| HET-HOM | *msh2* | 13993 | 917 | 557 | 8348 | 4171 | 6.67 |
| HET-HOM | *msh2* | 14127 | 674 | 592 | 8659 | 4202 | 6.83 |
| HET-HOM | *msh2* | 10330 | 815 | 338 | 4886 | 4291 | 6.91 |
| HOM-HET | Wild type | 1831 | 140 | 77 | 656 | 958 | 11.73 |
| HOM-HET | Wild type | 11047 | 688 | 556 | 4464 | 5339 | 12.45 |
| HOM-HET | Wild type | 6188 | 326 | 303 | 2368 | 3191 | 12.79 |
| HOM-HET | Wild type | 9297 | 564 | 464 | 4733 | 3536 | 9.81 |
| HOM-HET | *msh2* | 3346 | 148 | 66 | 1696 | 1436 | 3.89 |
| HOM-HET | *msh2* | 9555 | 193 | 162 | 6353 | 2847 | 2.54 |
| HOM-HET | *msh2* | 6420 | 176 | 82 | 2376 | 3786 | 3.45 |
| HOM-HET | *msh2* | 6479 | 172 | 80 | 2352 | 3875 | 3.41 |
| HOM-HET | *msh2* | 7468 | 200 | 108 | 2459 | 4701 | 4.39 |
| HOM-HET | *msh2* | 8940 | 441 | 155 | 3597 | 4747 | 4.31 |
| HOM-HET | *msh2* | 9039 | 376 | 155 | 4159 | 4349 | 3.72 |

**Supplementary Table 11.** Characteristics of a CTL panel (*2309-1*, *3090*, *3092*, *2696*, *2697*, *2698*, *3.13*, *3.9*) overlapping norther arm and centromeric region of chromosome 3. CTLs were created by Wu et al. (2015).

| **CTL name** | **Size (Mb)** | **Single-color reporter line** | | **Coordinates (bp)** | |
|---|---|---|---|---|---|
| | | **Left** | **Right** | **Left** | **Right** |
| *2309-1* | 1.36 | CR92 | CG50 | 1 171 464 | 2 528 742 |
| *3090* | 0.77 | CG50 | CR352 | 2 528 742 | 3 300 254 |
| *3092* | 1.05 | CG21 | CR73 | 3 280 460 | 4 330 342 |
| *2696* | 1.27 | CR73 | CG24 | 4 330 342 | 5 603 768 |
| *2697* | 1.16 | CG24 | CR1045 | 5 603 768 | 6 761 383 |
| *2698* | 1.26 | CR1045 | CG43 | 6 761 383 | 8 017 809 |
| *3.13* | 2.25 | CR729 | CG906 | 8 072 294 | 10 318 203 |
| *3.9* | 6.24 | CG17 | CR55 | 9 741 508 | 15 980 483 |

**Supplementary Table 12.** Fluorescent seed count data for the CTLs panel (*2309-1*, *3090*, *3092*, *2696*, *2697*, *2698*, *3.13*, *3.9*) for the wild type and *msh2* mutants in recombinant lines with differing pattern of heterozygosity. The following recombinant backgrounds were used: 'HOM-HOM' that are Col/Col inbred throughout the genome, 'HET-HET' that are Col/Ct heterozygous throughout the genome, 'HET-HOM' where the subtelomeric part of chromosome 3 is Col/Ct heterozygous and the remainder of chromosome 3 is Col/Col homozygous. Genetic distance in centimorgans (cM) is calculated as $cM = 100\times(1-(1-2(NG+NR)/NT)^{1/2})$, where NG is the number of green alone seeds, NR is the number of red alone seeds and NT is the total number of seeds of all classes analysed.

| CTL | Cross | Genotype | Total | Green alone | Red alone | Red+ Green | Non-colour | cM |
|---|---|---|---|---|---|---|---|---|
| *2309-1* | HOM-HOM | Wild type | 2598 | 95 | 91 | 1856 | 556 | 7.44 |
| *2309-1* | HOM-HOM | Wild type | 1789 | 37 | 73 | 1290 | 389 | 6.35 |
| *2309-1* | HOM-HOM | Wild type | 2188 | 77 | 80 | 1528 | 503 | 7.45 |
| *2309-1* | HOM-HOM | Wild type | 1619 | 57 | 56 | 1163 | 343 | 7.24 |
| *2309-1* | HOM-HOM | Wild type | 1682 | 47 | 71 | 1201 | 363 | 7.28 |
| *2309-1* | HOM-HOM | Wild type | 2108 | 59 | 98 | 1518 | 433 | 7.75 |
| *2309-1* | HOM-HOM | Wild type | 977 | 37 | 41 | 706 | 193 | 8.33 |
| *2309-1* | HOM-HOM | Wild type | 2433 | 108 | 93 | 1696 | 536 | 8.63 |
| *2309-1* | HOM-HOM | *msh2* | 2406 | 98 | 76 | 1679 | 553 | 7.51 |
| *2309-1* | HOM-HOM | *msh2* | 2192 | 110 | 54 | 1569 | 459 | 7.78 |
| *2309-1* | HOM-HOM | *msh2* | 1827 | 56 | 81 | 1311 | 379 | 7.80 |
| *2309-1* | HOM-HOM | *msh2* | 1583 | 50 | 77 | 1126 | 330 | 8.37 |
| *2309-1* | HOM-HOM | *msh2* | 1132 | 29 | 40 | 815 | 248 | 6.29 |
| *2309-1* | HOM-HOM | *msh2* | 1913 | 94 | 64 | 1339 | 416 | 8.63 |
| *2309-1* | HOM-HOM | *msh2* | 1868 | 76 | 106 | 1292 | 394 | 10.27 |
| *2309-1* | HOM-HOM | *msh2* | 1998 | 90 | 75 | 1384 | 449 | 8.63 |
| *3090* | HOM-HOM | Wild type | 1815 | 61 | 62 | 1280 | 412 | 7.02 |
| *3090* | HOM-HOM | Wild type | 2534 | 54 | 85 | 1796 | 599 | 5.64 |
| *3090* | HOM-HOM | Wild type | 2117 | 69 | 83 | 1511 | 454 | 7.46 |
| *3090* | HOM-HOM | Wild type | 2326 | 67 | 77 | 1676 | 506 | 6.40 |
| *3090* | HOM-HOM | Wild type | 2068 | 84 | 76 | 1458 | 450 | 8.06 |
| *3090* | HOM-HOM | Wild type | 1505 | 48 | 72 | 1062 | 323 | 8.32 |
| *3090* | HOM-HOM | Wild type | 2441 | 67 | 62 | 1718 | 594 | 5.43 |
| *3090* | HOM-HOM | *msh2* | 2597 | 72 | 88 | 1837 | 600 | 6.36 |
| *3090* | HOM-HOM | *msh2* | 2518 | 87 | 70 | 1769 | 592 | 6.44 |
| *3090* | HOM-HOM | *msh2* | 2074 | 98 | 70 | 1488 | 418 | 8.46 |
| *3090* | HOM-HOM | *msh2* | 2185 | 98 | 65 | 1563 | 459 | 7.76 |
| *3090* | HOM-HOM | *msh2* | 2233 | 122 | 50 | 1582 | 479 | 8.02 |
| *3090* | HOM-HOM | *msh2* | 2367 | 116 | 69 | 1673 | 509 | 8.15 |
| *3092* | HOM-HOM | Wild type | 2210 | 71 | 64 | 1591 | 484 | 6.31 |
| *3092* | HOM-HOM | Wild type | 2389 | 56 | 56 | 1695 | 582 | 4.80 |
| *3092* | HOM-HOM | Wild type | 1925 | 70 | 69 | 1355 | 431 | 7.50 |
| *3092* | HOM-HOM | Wild type | 2516 | 51 | 53 | 1812 | 600 | 4.22 |
| *3092* | HOM-HOM | Wild type | 2278 | 73 | 57 | 1614 | 534 | 5.88 |
| *3092* | HOM-HOM | Wild type | 1422 | 37 | 46 | 1040 | 299 | 6.02 |

| | | | | | | | | |
|---|---|---|---|---|---|---|---|---|
| *3092* | HOM-HOM | Wild type | 2536 | 62 | 75 | 1822 | 577 | 5.56 |
| *3092* | HOM-HOM | *msh2* | 2345 | 111 | 68 | 1657 | 509 | 7.95 |
| *3092* | HOM-HOM | *msh2* | 2083 | 68 | 83 | 1462 | 470 | 7.53 |
| *3092* | HOM-HOM | *msh2* | 2572 | 97 | 78 | 1812 | 585 | 7.05 |
| *3092* | HOM-HOM | *msh2* | 2546 | 73 | 119 | 1792 | 562 | 7.85 |
| *3092* | HOM-HOM | *msh2* | 2374 | 77 | 77 | 1666 | 554 | 6.71 |
| *3092* | HOM-HOM | *msh2* | 2389 | 87 | 80 | 1671 | 551 | 7.25 |
| *2696* | HOM-HOM | Wild type | 1587 | 41 | 51 | 1155 | 340 | 5.98 |
| *2696* | HOM-HOM | Wild type | 2442 | 63 | 81 | 1716 | 582 | 6.08 |
| *2696* | HOM-HOM | Wild type | 2407 | 93 | 64 | 1695 | 555 | 6.75 |
| *2696* | HOM-HOM | Wild type | 2418 | 66 | 84 | 1705 | 563 | 6.41 |
| *2696* | HOM-HOM | Wild type | 2157 | 62 | 86 | 1561 | 448 | 7.11 |
| *2696* | HOM-HOM | Wild type | 1278 | 42 | 46 | 925 | 265 | 7.14 |
| *2696* | HOM-HOM | *msh2* | 2106 | 128 | 70 | 1471 | 437 | 9.89 |
| *2696* | HOM-HOM | *msh2* | 1963 | 111 | 65 | 1354 | 433 | 9.41 |
| *2696* | HOM-HOM | *msh2* | 2217 | 125 | 79 | 1552 | 461 | 9.67 |
| *2696* | HOM-HOM | *msh2* | 1677 | 80 | 105 | 1156 | 336 | 11.72 |
| *2696* | HOM-HOM | *msh2* | 2016 | 123 | 71 | 1409 | 413 | 10.14 |
| *2696* | HOM-HOM | *msh2* | 2233 | 105 | 50 | 1601 | 477 | 7.20 |
| *2697* | HOM-HOM | Wild type | 2622 | 66 | 86 | 1859 | 611 | 5.98 |
| *2697* | HOM-HOM | Wild type | 2483 | 73 | 95 | 1781 | 534 | 7.01 |
| *2697* | HOM-HOM | Wild type | 2365 | 124 | 97 | 1662 | 482 | 9.83 |
| *2697* | HOM-HOM | Wild type | 1763 | 45 | 76 | 1247 | 395 | 7.12 |
| *2697* | HOM-HOM | Wild type | 1955 | 76 | 58 | 1375 | 446 | 7.11 |
| *2697* | HOM-HOM | Wild type | 2309 | 58 | 85 | 1637 | 529 | 6.40 |
| *2697* | HOM-HOM | Wild type | 1926 | 61 | 99 | 1461 | 305 | 8.68 |
| *2697* | HOM-HOM | Wild type | 2144 | 60 | 80 | 1645 | 359 | 6.76 |
| *2697* | HOM-HOM | Wild type | 2229 | 74 | 69 | 1718 | 368 | 6.64 |
| *2697* | HOM-HOM | Wild type | 2305 | 83 | 107 | 1675 | 440 | 8.61 |
| *2697* | HOM-HOM | Wild type | 2465 | 74 | 114 | 1760 | 517 | 7.94 |
| *2697* | HOM-HOM | Wild type | 2506 | 102 | 78 | 1820 | 506 | 7.46 |
| *2697* | HOM-HOM | Wild type | 2278 | 63 | 63 | 1688 | 464 | 5.69 |
| *2697* | HOM-HOM | Wild type | 2571 | 104 | 87 | 1853 | 527 | 7.73 |
| *2697* | HOM-HOM | Wild type | 2295 | 41 | 91 | 1689 | 474 | 5.93 |
| *2697* | HOM-HOM | Wild type | 2200 | 84 | 89 | 1597 | 430 | 8.20 |
| *2697* | HOM-HOM | *msh2* | 1028 | 30 | 27 | 811 | 160 | 5.71 |
| *2697* | HOM-HOM | *msh2* | 515 | 13 | 18 | 388 | 96 | 6.21 |
| *2697* | HOM-HOM | *msh2* | 509 | 20 | 28 | 361 | 100 | 9.92 |
| *2697* | HOM-HOM | *msh2* | 528 | 6 | 22 | 412 | 88 | 5.45 |
| *2698* | HOM-HOM | Wild type | 2301 | 97 | 98 | 1617 | 489 | 8.87 |
| *2698* | HOM-HOM | Wild type | 2354 | 68 | 143 | 1658 | 485 | 9.41 |
| *2698* | HOM-HOM | Wild type | 1029 | 37 | 30 | 730 | 232 | 6.74 |
| *2698* | HOM-HOM | Wild type | 1825 | 77 | 75 | 1292 | 381 | 8.71 |
| *2698* | HOM-HOM | Wild type | 921 | 26 | 44 | 658 | 193 | 7.91 |
| *2698* | HOM-HOM | Wild type | 2029 | 79 | 72 | 1448 | 430 | 7.74 |
| *2698* | HOM-HOM | *msh2* | 1894 | 80 | 75 | 1371 | 368 | 8.55 |

| | | | | | | | | |
|---|---|---|---|---|---|---|---|---|
| *2698* | HOM-HOM | *msh2* | 2352 | 72 | 74 | 1704 | 502 | 6.41 |
| *2698* | HOM-HOM | *msh2* | 1623 | 58 | 66 | 1168 | 331 | 7.96 |
| *2698* | HOM-HOM | *msh2* | 2472 | 79 | 82 | 1810 | 501 | 6.74 |
| *2698* | HOM-HOM | *msh2* | 2338 | 99 | 76 | 1698 | 465 | 7.79 |
| *2698* | HOM-HOM | *msh2* | 1524 | 63 | 56 | 1104 | 301 | 8.14 |
| *2698* | HOM-HOM | *msh2* | 1482 | 51 | 51 | 1085 | 295 | 7.14 |
| *2698* | HOM-HOM | *msh2* | 2323 | 110 | 91 | 1682 | 440 | 9.06 |
| *2698* | HOM-HOM | *msh2* | 2229 | 92 | 70 | 1611 | 456 | 7.55 |
| *3.13* | HOM-HOM | Wild type | 2598 | 159 | 127 | 1772 | 540 | 11.69 |
| *3.13* | HOM-HOM | Wild type | 2370 | 165 | 124 | 1627 | 454 | 13.04 |
| *3.13* | HOM-HOM | Wild type | 2630 | 165 | 131 | 1792 | 542 | 11.97 |
| *3.13* | HOM-HOM | Wild type | 2314 | 111 | 145 | 1591 | 467 | 11.75 |
| *3.13* | HOM-HOM | Wild type | 2332 | 132 | 161 | 1616 | 423 | 13.47 |
| *3.13* | HOM-HOM | Wild type | 2201 | 130 | 112 | 1519 | 440 | 11.68 |
| *3.13* | HOM-HOM | Wild type | 2201 | 111 | 105 | 1547 | 438 | 10.35 |
| *3.13* | HOM-HOM | Wild type | 2425 | 150 | 133 | 1645 | 497 | 12.44 |
| *3.13* | HOM-HOM | Wild type | 2518 | 128 | 170 | 1718 | 502 | 12.63 |
| *3.13* | HOM-HOM | Wild type | 2359 | 97 | 151 | 1630 | 481 | 11.13 |
| *3.13* | HOM-HOM | Wild type | 2302 | 133 | 104 | 1619 | 446 | 10.89 |
| *3.13* | HOM-HOM | Wild type | 2519 | 152 | 134 | 1725 | 508 | 12.08 |
| *3.13* | HOM-HOM | Wild type | 2398 | 145 | 102 | 1668 | 483 | 10.89 |
| *3.13* | HOM-HOM | Wild type | 2486 | 156 | 131 | 1683 | 516 | 12.30 |
| *3.13* | HOM-HOM | Wild type | 2336 | 130 | 133 | 1601 | 472 | 11.98 |
| *3.13* | HOM-HOM | *msh2* | 2386 | 158 | 121 | 1640 | 467 | 12.47 |
| *3.13* | HOM-HOM | *msh2* | 1801 | 105 | 118 | 1222 | 356 | 13.26 |
| *3.13* | HOM-HOM | *msh2* | 2542 | 141 | 161 | 1753 | 487 | 12.68 |
| *3.13* | HOM-HOM | *msh2* | 2669 | 166 | 138 | 1872 | 493 | 12.13 |
| *3.13* | HOM-HOM | *msh2* | 2210 | 149 | 184 | 1475 | 402 | 16.42 |
| *3.13* | HOM-HOM | *msh2* | 2595 | 161 | 126 | 1746 | 562 | 11.75 |
| *3.13* | HOM-HOM | *msh2* | 2669 | 139 | 145 | 1837 | 548 | 11.28 |
| *3.13* | HOM-HOM | *msh2* | 1176 | 63 | 71 | 817 | 225 | 12.13 |
| *3.9* | HOM-HOM | Wild type | 2376 | 211 | 155 | 1597 | 413 | 16.82 |
| *3.9* | HOM-HOM | Wild type | 986 | 70 | 68 | 676 | 172 | 15.14 |
| *3.9* | HOM-HOM | Wild type | 2386 | 208 | 167 | 1600 | 411 | 17.20 |
| *3.9* | HOM-HOM | Wild type | 2602 | 202 | 230 | 1716 | 454 | 18.27 |
| *3.9* | HOM-HOM | Wild type | 2619 | 223 | 206 | 1781 | 409 | 18.00 |
| *3.9* | HOM-HOM | Wild type | 2620 | 213 | 217 | 1724 | 466 | 18.04 |
| *3.9* | HOM-HOM | Wild type | 2475 | 210 | 184 | 1657 | 424 | 17.44 |
| *3.9* | HOM-HOM | Wild type | 2499 | 185 | 199 | 1683 | 432 | 16.77 |
| *3.9* | HOM-HOM | Wild type | 2377 | 226 | 157 | 1585 | 409 | 17.67 |
| *3.9* | HOM-HOM | Wild type | 2479 | 208 | 170 | 1641 | 460 | 16.63 |
| *3.9* | HOM-HOM | Wild type | 2416 | 208 | 201 | 1567 | 440 | 18.67 |
| *3.9* | HOM-HOM | Wild type | 2356 | 216 | 182 | 1538 | 420 | 18.63 |
| *3.9* | HOM-HOM | *msh2* | 2477 | 201 | 193 | 1617 | 466 | 17.42 |
| *3.9* | HOM-HOM | *msh2* | 2592 | 228 | 186 | 1718 | 460 | 17.50 |
| *3.9* | HOM-HOM | *msh2* | 2610 | 212 | 178 | 1790 | 430 | 16.27 |

| | | | | | | | | |
|---|---|---|---|---|---|---|---|---|
| *3.9* | HOM-HOM | *msh2* | 2610 | 247 | 171 | 1737 | 455 | 17.56 |
| *3.9* | HOM-HOM | *msh2* | 1591 | 131 | 158 | 1037 | 265 | 20.21 |
| *3.9* | HOM-HOM | *msh2* | 2214 | 217 | 213 | 1435 | 349 | 21.80 |
| *3.9* | HOM-HOM | *msh2* | 2405 | 198 | 221 | 1569 | 417 | 19.28 |
| *3.9* | HOM-HOM | *msh2* | 2610 | 221 | 200 | 1714 | 475 | 17.70 |
| *2309-1* | HET-HET | Wild type | 2224 | 69 | 77 | 1628 | 450 | 6.80 |
| *2309-1* | HET-HET | Wild type | 2224 | 98 | 67 | 1609 | 450 | 7.72 |
| *2309-1* | HET-HET | Wild type | 1614 | 64 | 60 | 1166 | 324 | 8.00 |
| *2309-1* | HET-HET | Wild type | 2340 | 75 | 61 | 1674 | 530 | 5.99 |
| *2309-1* | HET-HET | Wild type | 2430 | 77 | 66 | 1751 | 536 | 6.07 |
| *2309-1* | HET-HET | Wild type | 2350 | 40 | 88 | 1700 | 522 | 5.60 |
| *2309-1* | HET-HET | Wild type | 2172 | 89 | 107 | 1550 | 426 | 9.47 |
| *2309-1* | HET-HET | *msh2* | 1919 | 82 | 78 | 1333 | 426 | 8.72 |
| *2309-1* | HET-HET | *msh2* | 2022 | 92 | 85 | 1392 | 453 | 9.17 |
| *2309-1* | HET-HET | *msh2* | 1866 | 85 | 77 | 1348 | 356 | 9.10 |
| *3090* | HET-HET | Wild type | 2501 | 64 | 77 | 1764 | 596 | 5.81 |
| *3090* | HET-HET | Wild type | 2421 | 63 | 83 | 1705 | 570 | 6.22 |
| *3090* | HET-HET | Wild type | 2416 | 42 | 101 | 1742 | 531 | 6.11 |
| *3090* | HET-HET | Wild type | 2429 | 57 | 78 | 1781 | 513 | 5.72 |
| *3090* | HET-HET | Wild type | 2392 | 81 | 79 | 1688 | 544 | 6.93 |
| *3090* | HET-HET | Wild type | 2379 | 57 | 137 | 1679 | 506 | 8.52 |
| *3090* | HET-HET | *msh2* | 2228 | 57 | 121 | 1558 | 492 | 8.34 |
| *3090* | HET-HET | *msh2* | 1778 | 59 | 101 | 1247 | 371 | 9.44 |
| *3090* | HET-HET | *msh2* | 2200 | 76 | 76 | 1536 | 512 | 7.17 |
| *3090* | HET-HET | *msh2* | 1567 | 68 | 73 | 1075 | 351 | 9.44 |
| *3090* | HET-HET | *msh2* | 999 | 33 | 37 | 702 | 227 | 7.27 |
| *3090* | HET-HET | *msh2* | 1454 | 53 | 43 | 1011 | 347 | 6.84 |
| *3090* | HET-HET | *msh2* | 1477 | 47 | 51 | 1038 | 341 | 6.87 |
| *3092* | HET-HET | Wild type | 2299 | 51 | 80 | 1666 | 502 | 5.87 |
| *3092* | HET-HET | Wild type | 2226 | 70 | 51 | 1620 | 485 | 5.59 |
| *3092* | HET-HET | Wild type | 2270 | 87 | 66 | 1620 | 497 | 6.98 |
| *3092* | HET-HET | Wild type | 2243 | 69 | 72 | 1644 | 458 | 6.50 |
| *3092* | HET-HET | Wild type | 2528 | 108 | 56 | 1816 | 548 | 6.71 |
| *3092* | HET-HET | Wild type | 2319 | 80 | 79 | 1647 | 513 | 7.11 |
| *3092* | HET-HET | Wild type | 1864 | 87 | 86 | 1327 | 364 | 9.76 |
| *3092* | HET-HET | Wild type | 794 | 28 | 16 | 581 | 169 | 5.70 |
| *3092* | HET-HET | *msh2* | 2518 | 107 | 99 | 1746 | 566 | 8.55 |
| *3092* | HET-HET | *msh2* | 2410 | 75 | 95 | 1687 | 553 | 7.32 |
| *3092* | HET-HET | *msh2* | 2157 | 96 | 78 | 1525 | 458 | 8.42 |
| *3092* | HET-HET | *msh2* | 2379 | 80 | 99 | 1674 | 526 | 7.83 |
| *3092* | HET-HET | *msh2* | 2478 | 88 | 95 | 1733 | 562 | 7.68 |
| *3092* | HET-HET | *msh2* | 2407 | 89 | 88 | 1670 | 560 | 7.65 |
| *3092* | HET-HET | *msh2* | 2214 | 93 | 76 | 1596 | 449 | 7.95 |
| *2696* | HET-HET | Wild type | 2408 | 68 | 85 | 1704 | 551 | 6.57 |
| *2696* | HET-HET | Wild type | 2381 | 62 | 89 | 1677 | 553 | 6.56 |
| *2696* | HET-HET | Wild type | 2399 | 74 | 70 | 1731 | 524 | 6.19 |

| | | | | | | | | |
|---|---|---|---|---|---|---|---|---|
| *2696* | HET-HET | Wild type | 2463 | 61 | 86 | 1762 | 554 | 6.16 |
| *2696* | HET-HET | Wild type | 2369 | 63 | 113 | 1665 | 528 | 7.73 |
| *2696* | HET-HET | Wild type | 2461 | 54 | 82 | 1790 | 535 | 5.69 |
| *2696* | HET-HET | Wild type | 2450 | 48 | 75 | 1741 | 586 | 5.15 |
| *2696* | HET-HET | *msh2* | 2368 | 104 | 85 | 1643 | 536 | 8.33 |
| *2696* | HET-HET | *msh2* | 2164 | 71 | 61 | 1553 | 479 | 6.30 |
| *2696* | HET-HET | *msh2* | 2447 | 125 | 88 | 1706 | 528 | 9.12 |
| *2696* | HET-HET | *msh2* | 2261 | 84 | 82 | 1586 | 509 | 7.63 |
| *2696* | HET-HET | *msh2* | 2466 | 81 | 110 | 1761 | 514 | 8.07 |
| *2696* | HET-HET | *msh2* | 2470 | 121 | 68 | 1741 | 540 | 7.97 |
| *2696* | HET-HET | *msh2* | 2430 | 72 | 95 | 1701 | 562 | 7.13 |
| *2696* | HET-HET | *msh2* | 2414 | 110 | 77 | 1687 | 540 | 8.07 |
| *2696* | HET-HET | *msh2* | 2347 | 112 | 62 | 1679 | 494 | 7.71 |
| *2696* | HET-HET | *msh2* | 2303 | 76 | 63 | 1667 | 497 | 6.23 |
| *2696* | HET-HET | *msh2* | 2444 | 100 | 104 | 1693 | 547 | 8.73 |
| *2697* | HET-HET | Wild type | 2474 | 84 | 53 | 1806 | 531 | 5.70 |
| *2697* | HET-HET | Wild type | 2466 | 45 | 85 | 1789 | 547 | 5.42 |
| *2697* | HET-HET | Wild type | 2412 | 70 | 45 | 1709 | 588 | 4.89 |
| *2697* | HET-HET | Wild type | 2494 | 51 | 74 | 1824 | 545 | 5.14 |
| *2697* | HET-HET | Wild type | 2399 | 46 | 69 | 1745 | 539 | 4.91 |
| *2697* | HET-HET | Wild type | 2399 | 54 | 64 | 1712 | 569 | 5.05 |
| *2697* | HET-HET | Wild type | 2463 | 43 | 56 | 1767 | 597 | 4.10 |
| *2697* | HET-HET | Wild type | 2461 | 42 | 59 | 1805 | 555 | 4.19 |
| *2697* | HET-HET | *msh2* | 495 | 13 | 35 | 379 | 68 | 10.22 |
| *2697* | HET-HET | *msh2* | 573 | 17 | 44 | 409 | 103 | 11.28 |
| *2697* | HET-HET | *msh2* | 566 | 15 | 26 | 407 | 118 | 7.53 |
| *2697* | HET-HET | *msh2* | 1092 | 40 | 55 | 824 | 173 | 9.12 |
| *2697* | HET-HET | *msh2* | 1294 | 40 | 78 | 989 | 187 | 9.58 |
| *2697* | HET-HET | *msh2* | 593 | 29 | 34 | 408 | 122 | 11.26 |
| *2698* | HET-HET | Wild type | 2403 | 59 | 86 | 1754 | 504 | 6.23 |
| *2698* | HET-HET | Wild type | 2482 | 84 | 106 | 1787 | 505 | 7.97 |
| *2698* | HET-HET | Wild type | 2266 | 92 | 76 | 1642 | 456 | 7.71 |
| *2698* | HET-HET | Wild type | 2456 | 86 | 101 | 1777 | 492 | 7.93 |
| *2698* | HET-HET | Wild type | 2506 | 97 | 95 | 1786 | 528 | 7.98 |
| *2698* | HET-HET | Wild type | 2343 | 89 | 75 | 1699 | 480 | 7.26 |
| *2698* | HET-HET | Wild type | 2545 | 80 | 81 | 1867 | 517 | 6.54 |
| *2698* | HET-HET | Wild type | 2396 | 94 | 70 | 1726 | 506 | 7.10 |
| *2698* | HET-HET | *msh2* | 2156 | 82 | 76 | 1502 | 496 | 7.62 |
| *2698* | HET-HET | *msh2* | 2096 | 78 | 89 | 1470 | 459 | 8.31 |
| *2698* | HET-HET | *msh2* | 1540 | 58 | 60 | 1087 | 335 | 7.98 |
| *2698* | HET-HET | *msh2* | 1940 | 88 | 50 | 1375 | 427 | 7.39 |
| *2698* | HET-HET | *msh2* | 2093 | 95 | 78 | 1449 | 471 | 8.64 |
| *2698* | HET-HET | *msh2* | 1367 | 66 | 59 | 956 | 286 | 9.61 |
| *2698* | HET-HET | *msh2* | 1558 | 58 | 93 | 1082 | 325 | 10.21 |
| *2698* | HET-HET | *msh2* | 1654 | 86 | 60 | 1155 | 353 | 9.26 |
| *2698* | HET-HET | *msh2* | 814 | 31 | 29 | 576 | 178 | 7.66 |

| | | | | | | | | |
|---|---|---|---|---|---|---|---|---|
| *2698* | HET-HET | *msh2* | 2016 | 112 | 72 | 1405 | 427 | 9.59 |
| *2698* | HET-HET | *msh2* | 1418 | 73 | 64 | 978 | 303 | 10.18 |
| *3.13* | HET-HET | Wild type | 2294 | 115 | 162 | 1575 | 442 | 12.91 |
| *3.13* | HET-HET | Wild type | 2518 | 176 | 171 | 1688 | 483 | 14.89 |
| *3.13* | HET-HET | Wild type | 2084 | 130 | 127 | 1422 | 405 | 13.20 |
| *3.13* | HET-HET | Wild type | 1240 | 74 | 92 | 836 | 238 | 14.43 |
| *3.13* | HET-HET | *msh2* | 2386 | 164 | 118 | 1628 | 476 | 12.61 |
| *3.13* | HET-HET | *msh2* | 2260 | 150 | 155 | 1510 | 445 | 14.55 |
| *3.13* | HET-HET | *msh2* | 2520 | 162 | 113 | 1729 | 516 | 11.58 |
| *3.13* | HET-HET | *msh2* | 2192 | 106 | 125 | 1523 | 438 | 11.16 |
| *3.13* | HET-HET | *msh2* | 2319 | 144 | 133 | 1608 | 434 | 12.76 |
| *3.13* | HET-HET | *msh2* | 2321 | 156 | 107 | 1604 | 454 | 12.06 |
| *3.13* | HET-HET | *msh2* | 1981 | 115 | 118 | 1343 | 405 | 12.55 |
| *3.13* | HET-HET | *msh2* | 2202 | 153 | 115 | 1515 | 419 | 13.02 |
| *3.13* | HET-HET | *msh2* | 1898 | 146 | 104 | 1300 | 348 | 14.18 |
| *3.13* | HET-HET | *msh2* | 1185 | 74 | 72 | 815 | 224 | 13.19 |
| *3.13* | HET-HET | *msh2* | 1747 | 130 | 103 | 1183 | 331 | 14.37 |
| *3.13* | HET-HET | *msh2* | 1774 | 121 | 104 | 1190 | 359 | 13.61 |
| *3.9* | HET-HET | Wild type | 2098 | 210 | 186 | 1369 | 333 | 21.10 |
| *3.9* | HET-HET | Wild type | 1113 | 95 | 116 | 728 | 174 | 21.21 |
| *3.9* | HET-HET | Wild type | 2027 | 153 | 189 | 1365 | 320 | 18.60 |
| *3.9* | HET-HET | Wild type | 2395 | 199 | 230 | 1566 | 400 | 19.89 |
| *3.9* | HET-HET | Wild type | 2001 | 206 | 181 | 1293 | 321 | 21.69 |
| *3.9* | HET-HET | Wild type | 472 | 45 | 36 | 310 | 81 | 18.96 |
| *3.9* | HET-HET | *msh2* | 2002 | 135 | 128 | 1336 | 403 | 14.14 |
| *3.9* | HET-HET | *msh2* | 2122 | 144 | 129 | 1418 | 431 | 13.82 |
| *3.9* | HET-HET | *msh2* | 2095 | 144 | 132 | 1407 | 412 | 14.18 |
| *3.9* | HET-HET | *msh2* | 2047 | 147 | 129 | 1352 | 419 | 14.54 |
| *3.9* | HET-HET | *msh2* | 1458 | 102 | 94 | 974 | 288 | 14.49 |
| *3.9* | HET-HET | *msh2* | 1682 | 99 | 129 | 1160 | 294 | 14.62 |
| *2309-1* | HET-HOM | Wild type | 2501 | 76 | 129 | 1765 | 531 | 8.56 |
| *2309-1* | HET-HOM | Wild type | 2543 | 86 | 96 | 1849 | 512 | 7.43 |
| *2309-1* | HET-HOM | Wild type | 2543 | 97 | 88 | 1857 | 501 | 7.56 |
| *2309-1* | HET-HOM | Wild type | 2619 | 110 | 125 | 1845 | 539 | 9.42 |
| *2309-1* | HET-HOM | Wild type | 2424 | 93 | 106 | 1706 | 519 | 8.58 |
| *2309-1* | HET-HOM | Wild type | 2465 | 78 | 101 | 1789 | 497 | 7.55 |
| *2309-1* | HET-HOM | Wild type | 2376 | 107 | 85 | 1699 | 485 | 8.44 |
| *2309-1* | HET-HOM | *msh2* | 2231 | 73 | 134 | 1569 | 455 | 9.75 |
| *2309-1* | HET-HOM | *msh2* | 2140 | 63 | 85 | 1548 | 444 | 7.17 |
| *2309-1* | HET-HOM | *msh2* | 2345 | 64 | 64 | 1727 | 490 | 5.62 |
| *2309-1* | HET-HOM | *msh2* | 2339 | 66 | 100 | 1644 | 529 | 7.37 |
| *2309-1* | HET-HOM | *msh2* | 2203 | 145 | 91 | 1517 | 450 | 11.36 |
| *2309-1* | HET-HOM | *msh2* | 2307 | 105 | 85 | 1618 | 499 | 8.61 |
| *2309-1* | HET-HOM | *msh2* | 2202 | 62 | 84 | 1575 | 481 | 6.87 |
| *3090* | HET-HOM | Wild type | 2735 | 101 | 109 | 1919 | 606 | 8.00 |
| *3090* | HET-HOM | Wild type | 2601 | 81 | 118 | 1820 | 582 | 7.97 |

| | | | | | | | | |
|---|---|---|---|---|---|---|---|---|
| *3090* | HET-HOM | Wild type | 2553 | 111 | 80 | 1794 | 568 | 7.78 |
| *3090* | HET-HOM | Wild type | 2546 | 110 | 122 | 1816 | 498 | 9.57 |
| *3090* | HET-HOM | Wild type | 2653 | 107 | 88 | 1877 | 581 | 7.64 |
| *3090* | HET-HOM | Wild type | 2458 | 105 | 71 | 1727 | 555 | 7.44 |
| *3090* | HET-HOM | Wild type | 2355 | 96 | 55 | 1664 | 540 | 6.63 |
| *3090* | HET-HOM | *msh2* | 2129 | 91 | 105 | 1488 | 445 | 9.67 |
| *3090* | HET-HOM | *msh2* | 2189 | 124 | 103 | 1499 | 463 | 10.97 |
| *3090* | HET-HOM | *msh2* | 1408 | 73 | 46 | 982 | 307 | 8.84 |
| *3090* | HET-HOM | *msh2* | 1857 | 108 | 65 | 1288 | 396 | 9.80 |
| *3090* | HET-HOM | *msh2* | 2216 | 87 | 91 | 1573 | 465 | 8.38 |
| *3090* | HET-HOM | *msh2* | 1887 | 63 | 57 | 1379 | 388 | 6.58 |
| *3092* | HET-HOM | Wild type | 2515 | 76 | 83 | 1790 | 566 | 6.54 |
| *3092* | HET-HOM | Wild type | 2423 | 90 | 67 | 1722 | 544 | 6.70 |
| *3092* | HET-HOM | Wild type | 2335 | 83 | 72 | 1662 | 518 | 6.87 |
| *3092* | HET-HOM | Wild type | 2411 | 115 | 53 | 1706 | 537 | 7.23 |
| *3092* | HET-HOM | Wild type | 2584 | 70 | 81 | 1859 | 574 | 6.03 |
| *3092* | HET-HOM | Wild type | 2479 | 74 | 93 | 1761 | 551 | 6.98 |
| *3092* | HET-HOM | Wild type | 2497 | 74 | 65 | 1773 | 585 | 5.73 |
| *3092* | HET-HOM | *msh2* | 2346 | 77 | 72 | 1682 | 515 | 6.57 |
| *3092* | HET-HOM | *msh2* | 1545 | 54 | 48 | 1124 | 319 | 6.84 |
| *3092* | HET-HOM | *msh2* | 2315 | 96 | 85 | 1604 | 530 | 8.15 |
| *3092* | HET-HOM | *msh2* | 2357 | 102 | 79 | 1627 | 549 | 8.00 |
| *3092* | HET-HOM | *msh2* | 2287 | 102 | 88 | 1626 | 471 | 8.68 |
| *3092* | HET-HOM | *msh2* | 2213 | 82 | 90 | 1540 | 501 | 8.10 |
| *2696* | HET-HOM | Wild type | 2511 | 105 | 135 | 1752 | 519 | 10.06 |
| *2696* | HET-HOM | Wild type | 2402 | 103 | 131 | 1673 | 495 | 10.27 |
| *2696* | HET-HOM | Wild type | 2402 | 145 | 145 | 1623 | 489 | 12.91 |
| *2696* | HET-HOM | Wild type | 2505 | 141 | 127 | 1714 | 523 | 11.34 |
| *2696* | HET-HOM | Wild type | 2452 | 114 | 124 | 1684 | 530 | 10.23 |
| *2696* | HET-HOM | Wild type | 2402 | 157 | 91 | 1670 | 484 | 10.92 |
| *2696* | HET-HOM | Wild type | 2473 | 136 | 129 | 1701 | 507 | 11.36 |
| *2696* | HET-HOM | Wild type | 2452 | 148 | 110 | 1699 | 495 | 11.14 |
| *2696* | HET-HOM | *msh2* | 1547 | 53 | 49 | 1120 | 325 | 6.83 |
| *2696* | HET-HOM | *msh2* | 1969 | 77 | 61 | 1407 | 424 | 7.27 |
| *2696* | HET-HOM | *msh2* | 2407 | 118 | 71 | 1712 | 506 | 8.19 |
| *2696* | HET-HOM | *msh2* | 1951 | 65 | 66 | 1391 | 429 | 6.96 |
| *2696* | HET-HOM | *msh2* | 1888 | 93 | 57 | 1353 | 385 | 8.29 |
| *2696* | HET-HOM | *msh2* | 2464 | 111 | 86 | 1715 | 552 | 8.34 |
| *2696* | HET-HOM | *msh2* | 2350 | 111 | 53 | 1667 | 519 | 7.24 |
| *2696* | HET-HOM | *msh2* | 2387 | 117 | 75 | 1667 | 528 | 8.40 |
| *2696* | HET-HOM | *msh2* | 2316 | 99 | 80 | 1639 | 498 | 8.05 |
| *2696* | HET-HOM | *msh2* | 2192 | 81 | 65 | 1539 | 507 | 6.90 |
| *2696* | HET-HOM | *msh2* | 2417 | 108 | 60 | 1719 | 530 | 7.21 |
| *2696* | HET-HOM | *msh2* | 1808 | 78 | 48 | 1287 | 395 | 7.23 |
| *2696* | HET-HOM | *msh2* | 2180 | 97 | 88 | 1535 | 460 | 8.88 |
| *2697* | HET-HOM | Wild type | 2453 | 78 | 90 | 1766 | 519 | 7.10 |

| | | | | | | | | |
|---|---|---|---|---|---|---|---|---|
| *2697* | HET-HOM | Wild type | 2437 | 141 | 70 | 1717 | 509 | 9.07 |
| *2697* | HET-HOM | Wild type | 2426 | 60 | 91 | 1711 | 564 | 6.43 |
| *2697* | HET-HOM | Wild type | 2522 | 92 | 82 | 1831 | 517 | 7.16 |
| *2697* | HET-HOM | Wild type | 2406 | 120 | 61 | 1708 | 517 | 7.83 |
| *2697* | HET-HOM | Wild type | 2558 | 86 | 89 | 1863 | 520 | 7.09 |
| *2697* | HET-HOM | Wild type | 2448 | 59 | 103 | 1768 | 518 | 6.85 |
| *2697* | HET-HOM | *msh2* | 706 | 13 | 29 | 517 | 147 | 6.14 |
| *2697* | HET-HOM | *msh2* | 545 | 8 | 24 | 439 | 74 | 6.05 |
| *2697* | HET-HOM | *msh2* | 529 | 24 | 34 | 371 | 100 | 11.64 |
| *2697* | HET-HOM | *msh2* | 569 | 10 | 15 | 410 | 134 | 4.49 |
| *2697* | HET-HOM | *msh2* | 541 | 27 | 32 | 383 | 99 | 11.58 |
| *2697* | HET-HOM | *msh2* | 675 | 16 | 28 | 487 | 144 | 6.75 |
| *2697* | HET-HOM | *msh2* | 625 | 19 | 34 | 450 | 122 | 8.87 |
| *2698* | HET-HOM | Wild type | 2612 | 109 | 53 | 1854 | 596 | 6.41 |
| *2698* | HET-HOM | Wild type | 2667 | 69 | 90 | 1929 | 579 | 6.15 |
| *2698* | HET-HOM | Wild type | 2612 | 56 | 105 | 1893 | 558 | 6.37 |
| *2698* | HET-HOM | Wild type | 2503 | 64 | 76 | 1822 | 541 | 5.76 |
| *2698* | HET-HOM | Wild type | 2646 | 45 | 108 | 1915 | 578 | 5.96 |
| *2698* | HET-HOM | Wild type | 2644 | 66 | 73 | 1945 | 560 | 5.40 |
| *2698* | HET-HOM | Wild type | 2558 | 85 | 64 | 1857 | 552 | 6.01 |
| *2698* | HET-HOM | *msh2* | 2086 | 94 | 108 | 1451 | 433 | 10.20 |
| *2698* | HET-HOM | *msh2* | 2160 | 116 | 87 | 1517 | 440 | 9.89 |
| *2698* | HET-HOM | *msh2* | 2206 | 126 | 74 | 1543 | 463 | 9.52 |
| *2698* | HET-HOM | *msh2* | 2165 | 125 | 121 | 1508 | 411 | 12.09 |
| *2698* | HET-HOM | *msh2* | 2231 | 120 | 83 | 1562 | 466 | 9.56 |
| *2698* | HET-HOM | *msh2* | 2052 | 98 | 80 | 1456 | 418 | 9.09 |
| *2698* | HET-HOM | *msh2* | 2178 | 82 | 97 | 1544 | 455 | 8.59 |
| *3.13* | HET-HOM | Wild type | 2561 | 80 | 45 | 1869 | 567 | 5.01 |
| *3.13* | HET-HOM | Wild type | 2512 | 52 | 72 | 1821 | 567 | 5.06 |
| *3.13* | HET-HOM | Wild type | 2512 | 68 | 45 | 1833 | 566 | 4.60 |
| *3.13* | HET-HOM | Wild type | 2447 | 132 | 116 | 1711 | 488 | 10.71 |
| *3.13* | HET-HOM | Wild type | 2292 | 109 | 108 | 1644 | 431 | 9.96 |
| *3.13* | HET-HOM | Wild type | 2376 | 62 | 74 | 1742 | 498 | 5.90 |
| *3.13* | HET-HOM | *msh2* | 2278 | 149 | 166 | 1535 | 428 | 14.94 |
| *3.13* | HET-HOM | *msh2* | 2488 | 136 | 152 | 1741 | 459 | 12.34 |
| *3.13* | HET-HOM | *msh2* | 2467 | 104 | 161 | 1716 | 486 | 11.39 |
| *3.13* | HET-HOM | *msh2* | 2473 | 142 | 148 | 1687 | 496 | 12.51 |
| *3.13* | HET-HOM | *msh2* | 2142 | 126 | 126 | 1443 | 447 | 12.55 |
| *3.13* | HET-HOM | *msh2* | 2200 | 131 | 139 | 1476 | 454 | 13.14 |
| *3.13* | HET-HOM | *msh2* | 2305 | 129 | 154 | 1567 | 455 | 13.14 |
| *3.13* | HET-HOM | *msh2* | 2034 | 111 | 119 | 1369 | 435 | 12.03 |
| *3.13* | HET-HOM | *msh2* | 2472 | 137 | 137 | 1760 | 438 | 11.78 |
| *3.9* | HET-HOM | Wild type | 2315 | 169 | 203 | 1524 | 419 | 17.62 |
| *3.9* | HET-HOM | Wild type | 2497 | 195 | 142 | 1695 | 465 | 14.56 |
| *3.9* | HET-HOM | Wild type | 1890 | 138 | 129 | 1278 | 345 | 15.30 |
| *3.9* | HET-HOM | Wild type | 2201 | 156 | 159 | 1528 | 358 | 15.52 |

| | | | | | | | | |
|---|---|---|---|---|---|---|---|---|
| *3.9* | HET-HOM | Wild type | 2540 | 187 | 208 | 1700 | 445 | 17.00 |
| *3.9* | HET-HOM | Wild type | 2324 | 164 | 149 | 1561 | 450 | 14.52 |
| *3.9* | HET-HOM | Wild type | 2481 | 179 | 150 | 1682 | 470 | 14.28 |
| *3.9* | HET-HOM | Wild type | 2537 | 217 | 181 | 1673 | 466 | 17.16 |
| *3.9* | HET-HOM | *msh2* | 2242 | 242 | 177 | 1475 | 348 | 20.87 |
| *3.9* | HET-HOM | *msh2* | 2346 | 172 | 183 | 1585 | 406 | 16.49 |
| *3.9* | HET-HOM | *msh2* | 2070 | 183 | 141 | 1379 | 367 | 17.12 |
| *3.9* | HET-HOM | *msh2* | 1863 | 200 | 129 | 1219 | 315 | 19.58 |
| *3.9* | HET-HOM | *msh2* | 2169 | 217 | 152 | 1444 | 356 | 18.77 |
| *3.9* | HET-HOM | *msh2* | 2391 | 191 | 201 | 1595 | 404 | 18.02 |
| *3.9* | HET-HOM | *msh2* | 2363 | 217 | 164 | 1572 | 410 | 17.69 |
| *3.9* | HET-HOM | *msh2* | 2100 | 185 | 138 | 1418 | 359 | 16.79 |

**Supplementary Table 13.** *I3bc* FTL flow cytometry count data in wild type and *msh2* mutant in Col/Col inbred and Col/Ct hybrid. There are eight possible fluorescent pollen classes: BYR, indicating pollen with blue (B), yellow (Y) and red (R) fluorescence, byr, indicating non-fluorescent pollen, bYr, ByR, BYr, byR, bYR, Byr.

| Cross | Genotype | Replicate | BYR | byr | bYr | ByR | BYr | byR | bYR | Byr | Total |
|---|---|---|---|---|---|---|---|---|---|---|---|
| Col/Col | Wild type | 1 | 12627 | 12318 | 68 | 134 | 1004 | 959 | 2274 | 2277 | 31661 |
| Col/Col | Wild type | 2 | 15514 | 15509 | 100 | 222 | 1486 | 1542 | 2704 | 2808 | 39885 |
| Col/Col | Wild type | 3 | 16684 | 16530 | 86 | 173 | 1520 | 1495 | 3474 | 3739 | 43701 |
| Col/Col | Wild type | 4 | 3407 | 3598 | 23 | 56 | 411 | 371 | 561 | 611 | 9038 |
| Col/Col | Wild type | 5 | 4375 | 4425 | 16 | 43 | 335 | 378 | 815 | 820 | 11207 |
| Col/Col | Wild type | 6 | 5936 | 5939 | 27 | 76 | 459 | 544 | 1041 | 1029 | 15051 |
| Col/Col | *msh2* | 1 | 8610 | 8566 | 75 | 90 | 1251 | 857 | 744 | 1215 | 21408 |
| Col/Col | *msh2* | 2 | 14947 | 13896 | 74 | 116 | 1592 | 1569 | 1261 | 1430 | 34885 |
| Col/Col | *msh2* | 3 | 6832 | 7603 | 39 | 76 | 915 | 845 | 1693 | 1598 | 19601 |
| Col/Col | *msh2* | 4 | 6091 | 6012 | 35 | 71 | 678 | 713 | 952 | 944 | 15496 |
| Col/Col | *msh2* | 5 | 7328 | 7187 | 54 | 127 | 870 | 902 | 1059 | 1143 | 18670 |
| Col/Col | *msh2* | 6 | 4945 | 4943 | 18 | 57 | 470 | 443 | 660 | 722 | 12258 |
| Col/Col | *msh2* | 7 | 6033 | 6044 | 30 | 79 | 678 | 639 | 912 | 942 | 15357 |
| Col/Col | *msh2* | 8 | 6397 | 6363 | 35 | 137 | 643 | 761 | 1206 | 1295 | 16837 |
| Col/Col | *msh2* | 9 | 4175 | 4014 | 14 | 47 | 445 | 478 | 604 | 692 | 10469 |
| Col/Ct | Wild type | 1 | 4329 | 4970 | 7 | 50 | 482 | 487 | 709 | 441 | 11475 |
| Col/Ct | Wild type | 2 | 4483 | 4454 | 17 | 42 | 518 | 571 | 792 | 766 | 11643 |
| Col/Ct | Wild type | 3 | 3876 | 3859 | 7 | 44 | 441 | 447 | 599 | 548 | 9821 |
| Col/Ct | Wild type | 4 | 8151 | 8129 | 37 | 166 | 1203 | 1272 | 1209 | 926 | 21093 |
| Col/Ct | Wild type | 5 | 8442 | 8549 | 24 | 137 | 1282 | 1283 | 1147 | 747 | 21611 |
| Col/Ct | Wild type | 6 | 6046 | 7541 | 18 | 97 | 1000 | 1004 | 910 | 709 | 17325 |
| Col/Ct | Wild type | 7 | 4971 | 5098 | 12 | 81 | 667 | 592 | 785 | 723 | 12929 |
| Col/Ct | Wild type | 8 | 4261 | 4352 | 7 | 52 | 475 | 512 | 769 | 684 | 11112 |
| Col/Ct | Wild type | 9 | 4088 | 3980 | 10 | 46 | 533 | 557 | 674 | 556 | 10444 |
| Col/Ct | *msh2* | 1 | 5025 | 6183 | 41 | 88 | 1088 | 1138 | 918 | 811 | 15292 |
| Col/Ct | *msh2* | 2 | 3735 | 4630 | 25 | 66 | 693 | 608 | 797 | 874 | 11428 |
| Col/Ct | *msh2* | 3 | 3820 | 3980 | 24 | 65 | 640 | 590 | 758 | 422 | 10299 |
| Col/Ct | *msh2* | 4 | 4510 | 5715 | 29 | 104 | 817 | 973 | 874 | 819 | 13841 |
| Col/Ct | *msh2* | 5 | 3917 | 4526 | 28 | 90 | 788 | 775 | 784 | 642 | 11550 |
| Col/Ct | *msh2* | 6 | 3278 | 3224 | 11 | 54 | 573 | 569 | 560 | 478 | 8747 |

**Supplementary Table 14.** *I3bc* FTL interference count data in wild type and *msh2* mutant in Col/Col inbred and Col/Ct hybrid. *I3bc* interference (1 – CoC) is calculated with the use of genetic distances of adjacent *I3b* and *I3c* intervals, to estimate the number of DCO pollen expected in the absence of interference. The ratio of “observed DCOs” [(*I3b* cM/100) x (*I3c* cM/100) x total pollen number] to expected “DCOs” gives the coefficient of coincidence (CoC).

| **Cross** | **Genotype** | **Replicate** | ***I3b* cM** | ***I3c* cM** | **Expected DCOs** | **Observed DCOs** | **1-CoC** |
|---|---|---|---|---|---|---|---|
| Col/Col | Wild type | 1 | 21.21 | 6.84 | 459 | 202 | 0.56 |
| Col/Col | Wild type | 2 | 22.22 | 8.40 | 744 | 322 | 0.57 |
| Col/Col | Wild type | 3 | 24.00 | 7.49 | 786 | 259 | 0.67 |
| Col/Col | Wild type | 4 | 22.49 | 9.53 | 194 | 79 | 0.59 |
| Col/Col | Wild type | 5 | 21.48 | 6.89 | 166 | 59 | 0.64 |
| Col/Col | Wild type | 6 | 21.10 | 7.35 | 233 | 103 | 0.56 |
| Col/Col | *msh2* | 1 | 19.00 | 10.62 | 432 | 165 | 0.62 |
| Col/Col | *msh2* | 2 | 16.78 | 9.61 | 562 | 190 | 0.66 |
| Col/Col | *msh2* | 3 | 25.77 | 9.57 | 483 | 115 | 0.76 |
| Col/Col | *msh2* | 4 | 21.90 | 9.66 | 328 | 106 | 0.68 |
| Col/Col | *msh2* | 5 | 22.25 | 10.46 | 435 | 181 | 0.58 |
| Col/Col | *msh2* | 6 | 19.33 | 8.06 | 191 | 75 | 0.61 |
| Col/Col | *msh2* | 7 | 21.36 | 9.29 | 305 | 109 | 0.64 |
| Col/Col | *msh2* | 8 | 24.21 | 9.36 | 382 | 172 | 0.55 |
| Col/Col | *msh2* | 9 | 21.78 | 9.40 | 214 | 61 | 0.72 |
| Col/Ct | Wild type | 1 | 18.47 | 8.94 | 189 | 57 | 0.70 |
| Col/Ct | Wild type | 2 | 22.73 | 9.86 | 261 | 59 | 0.77 |
| Col/Ct | Wild type | 3 | 20.72 | 9.56 | 195 | 51 | 0.74 |
| Col/Ct | Wild type | 4 | 21.86 | 12.70 | 585 | 203 | 0.65 |
| Col/Ct | Wild type | 5 | 20.63 | 12.61 | 562 | 161 | 0.71 |
| Col/Ct | Wild type | 6 | 20.91 | 12.23 | 443 | 115 | 0.74 |
| Col/Ct | Wild type | 7 | 21.40 | 10.46 | 289 | 93 | 0.68 |
| Col/Ct | Wild type | 8 | 21.96 | 9.41 | 230 | 59 | 0.74 |
| Col/Ct | Wild type | 9 | 22.21 | 10.97 | 255 | 56 | 0.78 |
| Col/Ct | *msh2* | 1 | 25.86 | 15.40 | 609 | 129 | 0.79 |
| Col/Ct | *msh2* | 2 | 26.01 | 12.18 | 362 | 91 | 0.75 |
| Col/Ct | *msh2* | 3 | 23.40 | 12.81 | 309 | 89 | 0.71 |
| Col/Ct | *msh2* | 4 | 25.16 | 13.89 | 484 | 133 | 0.73 |
| Col/Ct | *msh2* | 5 | 25.88 | 14.55 | 435 | 118 | 0.73 |
| Col/Ct | *msh2* | 6 | 24.92 | 13.80 | 301 | 65 | 0.78 |

**Supplementary Table 15.** List of oligonucleotides.

| ID | Primer sequence | Comments |
|---|---|---|
| **JZ-001** | TTAAACTCACTTGGCAATCGC | *msh2-1* SALK mutant in Col, use together with LBb1.3 |
| **JZ-002** | TAAGCTTTGCTGATTTGGCTG | |
| **JZ-14** | AACCACTGCTCTACGTCAGC | *msh2* CRISPR/Cas9 deletion mutants genotyping |
| **JZ-15** | TCGCTTCAGATGCTGTCTAA | |
| **zip4-F** | TTGCTACCTTGGGCTCTCTC | *zip4-2* SALK mutant in Col, use together with LBb1.3 |
| **zip4-R** | ATTCTGTTCTCGCTTTCCAG | |
| **zip4_CRISPR_F** | TCTCCCGCCAACAATCGAAA | *zip4-3* CRISPR/Cas9 deletion mutant genotyping in L*er* |
| **zip4_CRISPR_R** | CTGCAATCGTCGTCACCTCT | |
| **fancm-dCAPS-F** | ACAATATATGTTTCGTGCAGGTAAGAC ATTGGAAG | *fancm-1* mutant in Col, dCAPS primers: digestion with MboII enzyme |
| **fancm-dCAPS-R** | CACCAATAGATGTTGCGACAAT | |
| **fancm-Ler-L** | GCTTGGTGATCGACGAGGCACATCAA G | *fancm-10* mutant in L*er*, dCAPS primers: digestion with HindIII enzyme |
| **fancm-Ler-R** | ATTAGTTGGTCACCTTGATCAT | |
| **recq4a-F** | ATCAGAGCCACTCATTGTTG | *recq4a-4* mutant in Col, multiplex PCR |
| **recq4a-wt-R** | GTCCTGATCGTGTTGGACAG | |
| **recq4a-mut-R** | ATATTGACCATCATACTCATTGC | |
| **recq4b-wt-F** | TCAGAAAGTTGCTCTGCGTC | *recq4b-2* wild type allele in Col |
| **recq4b-wt-R** | ACCAAGACCCTGCATATTGC | |
| **recq4b-mut-F** | ACTAGAGATACTTCAGGAGCTGAGC | *recq4b-2* mutant allele in Col |
| **recq4b-mut-R** | GCTTTCTTCCCTTCCTTTCTC | |
| **recq4a_Ler_F** | GACCAAGGCAGATATGCCTGTGATACC TG | *recq4a* mutant in L*er*, dCAPS primers: digestion with ScrFI enzyme |
| **recq4a_Ler_R** | GTCCAGGGAAATTCACGACTGGTC | |
| **LBb1.3** | ATTTTGCCGATTTCGGAAC | Universal primer for SALK mutants |
| **HEI10_C2_F** | TAATACGACTCACTATAGGG | Genotyping of *HEI10-OE* construct presence in C2 line |
| **HEI10_C2_R** | GCCAGCAAGACAGAACAGTTC | |
| **ZIP4-gRNA1** | AAGTTGACTCAGCCGAGTT | gRNAs binding within 1st exon of *ZIP4* |
| **ZIP4-gRNA2** | AATCGTCGGAGATGGAACAC | |

**Supplementary Table 16.** Summary of the GBS libraries and crossovers identified.

| Genotype | Crossovers per chromosome | | | | | Total crossover number | Total reads | No. of individuals | Avg. reads per library |
|---|---|---|---|---|---|---|---|---|---|
| | Chr1 | Chr2 | Chr3 | Chr4 | Chr5 | | | | |
| *fancm zip4* | 194 | 195 | 209 | 264 | 199 | 1061 | 109121099 | 232 | 470349 |
| *msh2 fancm* | 487 | 357 | 364 | 316 | 452 | 1976 | 117537405 | 201 | 584763 |
| *msh2 fancm zip4* | 314 | 227 | 277 | 230 | 276 | 1324 | 69456952 | 184 | 377483 |
| *msh2 recq4ab* | 2232 | 1418 | 1781 | 1129 | 2013 | 8573 | 196367255 | 283 | 693877 |
